# Glucose and methylglyoxal alter plasma protein dynamics, immune traits and glucose homeostasis in a sex- and season-dependent manner in zebra finches

**DOI:** 10.64898/2026.07.14.737735

**Authors:** Adrián Moreno Borrallo, François Criscuolo, Fabrice Bertile

## Abstract

Birds maintain high circulating glucose levels compared with other vertebrates without developing the diabetic complications observed in mammals, yet the mechanisms underlying this resistance remain unclear. We explored the effects of chronic one-year glucose and methylglyoxal (a reactive dicarbonyl compound causing glycative stress) supplementation on physiological condition in zebra finches (*Taeniopygia guttata*), with emphasis on sex- and season-dependent variation in plasma biochemistry and immune traits. Ninety zebra finches were randomly assigned to control, glucose or methylglyoxal-supplemented drinking treatments. We analysed the effects on plasma proteins, metabolites (glucose, uric acid), tissue damage markers like aspartate aminotransferase (AST), creatine kinase (CK) and bile acids, electrolytes, and leukocyte profiles. Both supplementations increased plasma glucose concentrations, with methylglyoxal producing the strongest effect. The treatments, particularly glucose, disrupted plasma protein dynamics, preventing the spring increase in total proteins in both sexes and globulins in females. This suggests alterations in reproductive-related metabolism. Glucose supplementation elevated the heterophil-to-lymphocyte (H/L) ratio in May and August, consistent with elevated physiological stress. However, methylglyoxal supplementation reduced the H/L ratio in November and unexpectedly lowered plasma AST and CK concentrations in May, despite its well-established pro-oxidative properties, suggesting hormetic mechanisms. Overall, our findings show that glucose and methylglyoxal reshape physiology in zebra finches in a strongly sex- and season-dependent manner rather than simply inducing generalized metabolic damage. These results highlight the importance of considering both biological context and standardized haematological reference values when investigating glucose metabolism in birds.

## Introduction

Birds provide compelling models for studying adaptations in carbohydrate physiology, given their exceptionally high circulating glucose levels relative to other vertebrates (Polakof et al. 2011). This trait is particularly notable given that birds exhibit a body mass-adjusted lifespan exceeding that of similarly sized endothermic mammals (Lindstedt and Calder 1976), despite the potential deleterious effects of reducing sugars. These effects arise because reducing sugars, such as glucose, non-enzymatically react with certain nitrogen-containing compounds, such as proteins and DNA, a process known as glycation (Maillard 1912; Suji and Sivakami 2004). Glycation can impair protein function (e.g. Vetter and Indurthi 2011) and, together with oxidative stress, contributes to the formation of Advanced Glycation End-products (AGEs; Poulsen et al. 2013; Twarda-Clapa et al. 2022). In humans, AGEs and protein glycation, are increased in diabetes, and AGE tissue accumulation contributes to various pathologies (Cerami et al. 1986; Ulrich and Cerami 2001; Chaudhuri et al. 2018; Khalid et al. 2022) and influences mortality hazard (Nin et al. 2011). These effects are partly mediated by proinflammatory consequences of AGEs, primarily through the membrane Receptors of AGEs (RAGEs) (Hofmann et al. 1999; Ramasamy et al. 2011). However, little is known about how glucose and AGEs affect inflammation and immune function in birds, which appear to lack RAGEs (Eythrib 2013; Szwergold and Miller 2014a). A recent study in chicken shows that AGEs induce proinflammatory signalling in peripheral blood leukocytes *via* an unidentified receptor (Wein et al. 2020). Another study suggests that streptozotocin-induced increases in blood glucose and glycation levels do not elevate AGE formation in chicken, possibly due to enhanced scavenging by free amino acids (Szwergold and Miller 2014b). This is associated with a reduced or absent proinflammatory response in most tissues under streptozotocin induced hyperglycaemic conditions (Luo et al. 2025). Overall, birds seem to have evolved adaptations that mitigate the impact of glucose on oxidative stress (Vágási et al. 2024) and glycation (Zuck et al. 2017; Brun et al. 2022), although the underlying mechanisms remain speculative (Szwergold and Miller 2014a; 2014b; Anthony-Regnitz et al. 2020).

Although a few studies have linked glucose regulation to survival in birds (Montoya et al. 2022; 2018) and highlighted the ecological relevance of protein glycation, including age-related variation in haemoglobin glycation and a negative impact on survival (Récapet et al. 2016), experimental evidence remains scarce. Our previous work shows that glucose supplementation in drinking water increases albumin glycation and mortality in zebra finches, while methylglyoxal (a reactive dicarbonyl compound derived from glucose degradation and inducing AGE formation; Vašková et al. 2025), enhances DNA damage and increases plasma AGE levels and red blood cell apoptosis (Moreno Borrallo et al. 2026a, 2026b). Nevertheless, the mechanisms underlying the increased mortality remain unclear, as none of the measured physiological variables explained the elevated hazard (Moreno Borrallo et al. 2026a, 2026b).

In the current study, mostly exploratory, we further investigate the potential effects of glucose and methylglyoxal supplementation on haematological parameters and organ damage indicators to elucidate their impact on birds’ health. We analyse aspartate aminotransferase (AST), a transaminase enzyme whose plasma levels, combined with creatine kinase (CK), are used to diagnose acute hepatic damage and muscle damage (see Harr 2002 and references therein). Since AST is present in both hepatocytes and myocytes, and CK only in myocytes, elevated AST with normal CK levels suggests liver damage (Harr 2002). Comorbidity suggesting pancreatic and hepatic involvement has been reported in several cases of diabetes in birds (Weyer et al. 2024). However, these plasma enzymatic markers can be misleading, as they do not always correlate with cytological indicators of hepatic damage, particularly in cases of chronic fibrosis (Lumeij 1997). We therefore complement our analysis with total bile acids (TBA) measurements, as previously recommended (Cray et al. 2008).

We also measure plasma protein content (total protein, albumin and globulin), uric acid (a main protein catabolite with debated antioxidant properties, McGraw 2011; Skrip and McWilliams 2016; Peglow Pinz et al. 2025), and several electrolytes (phosphates, calcium, potassium and sodium), to evaluate kidney function and hydration status (Jones 1999). Finally, we assess immune function using the heterophil-to-lymphocyte ratio (H/L), monocyte counts and albumin- to-globulin ratio (A/G), which are markers of inflammation and stress (Harr 2002; Davis et al. 2008, 2018). In zebra finches, the H/L ratio is already known to vary with stressful conditions (Birkhead and Fletcher 1998; Udino et al. 2024a; 2024b; Colominas-Ciuró et al. 2024). Overall, our study therefore aims to explore further the effects of glucose and methylglyoxal supplementation on avian health and to identify additional mechanisms potentially mediating the relationship between glucose and mortality in birds.

## Material and methods

### General protocol

A total of 90 zebra finches, individually identified with leg rings and originating from diverse captive sources and breeds (see details in **ESM1**), were randomly assigned to three experimental groups (30 birds per group, 15 males, 15 females), using the *sample()* function in R (R Core Team 2025). We verified that the body mass and age (known for 86 out of 90 birds) were not significantly different across groups (Moreno Borrallo et al. 2026a). Birds were housed in outdoor aviaries at the DEPE – IPHC animal facilities (Strasbourg), with *ad libitum* access to food, grit and shelter (for housing details, see Moreno Borrallo et al. 2026a; for environmental information throughout the whole experiment, i.e. maximum and minimum daily temperature and humidity, see **ESM3**, and for photoperiod https://www.timeanddate.com/sun/france/strasbourg?month=9&year=2022 and https://www.timeanddate.com/sun/france/strasbourg?month=9&year=2023). Water was provided *ad libitum* and supplemented with either glucose (50 g/L) or methylglyoxal (8.33 g/L), or left without supplementation (control group). Supplement concentrations were determined based on a pilot study (see Moreno Borrallo et al. 2026a).

The experimentation ran from the 21^st^ of September 2022 to the 25^th^ of September 2023. During the experiment, 48 out of the 90 birds died (53.33 %), with mortality significantly higher in the glucose-supplemented group than in the other groups (Moreno Borrallo et al. 2026a). Several baseline measurements were taken in August 2022, with subsequent samplings every three months (for a detailed schedule, see Moreno Borrallo et al. 2026a; 2026b). For the H/L ratio (see below), blood samples were collected in November 2022, February, May and August 2023. For other haematological parameters, only November 2022 and May 2023 samples were analysed. Immunological assays were performed one week after the other parameter measurements.

Birds were captured from aviaries the day before blood sampling and housed overnight in small indoor cages (40.5 cm x 54.5 cm x 29.5 cm), with free access to water but not food. Blood samples (approximately 100 µL) were collected in the morning from the alar vein using lancets and heparinized capillary tubes, then placed in heparinized Microvette® tubes. They were kept on ice until processing and centrifuged at 3500 g for 10 min at 4° C to separate plasma and cells (see more details on Moreno Borrallo et al. 2026a; 2026b). All assays were conducted on fresh plasma on the same day.

### Plasma haematology

The following plasma parameters were determined: total proteins (TP; g/dL), albumin (g/dL), globulin (g/dL), albumin/globulin ratio (A/G), aspartate aminotransferase (AST; U/L), total bile acids (TBA; µmol/L), creatine kinase (CK; U/L), uric acid (UA; mg/dL), glucose (mg/dL), Ca^2+^ (mg/dL), phosphates (mg/dL), K^+^ (mmol/L) and Na^+^ (mmol/L). Measurements were performed using a veterinary analyser (Element RC®, Scil®) equipped with an avian-reptilian rotor (108085) using a colorimetric technique. A very similar veterinary machine (Zoetis VetScan VS2) has already been used for the determination of the exact same parameters in birds, including zebra finches (Xiong et al. 2023). Because of limited plasma volume and potential values exceeding quantification limits, all samples (n = 117) were diluted with sterile reverse osmosis water (2x, n = 78; 2.5x, n = 14; 4x, n = 16; 5x, n = 4; 8x, n = 5). Final (diluted) plasma volume was always 100 µL, as this is the requirement for the rotors. Final concentrations were corrected for the dilution factor.

For some parameters, the Element RC® machine sometimes returned values below (all except glucose, CK and Ca^2+^) or above (AST and UA) certain limits (reported as “>” or “<”), which were defined as Limits of Quantification (LOQ). After dilution correction, these values were included in the statistical analyses as censored data (i.e. as intervals instead of point values). For three parameters (total proteins, globulin and the A/G ratio), the machine sometimes returned no numerical output at all (”---”), which we treated as a null reading below the Limit of Detection (LOD). No external standards or calibrators were run: the analyser relies on its manufacturer’s internal calibration and only returns an output value (a number, “< X”, “> X” or “---”). We therefore approximated the LOD of each of these parameters as the minimum non-null raw value observed in our dataset multiplied by the corresponding dilution factor: since any “---” reading necessarily lies below the lowest value the machine was able to quantify in our data, treating it as censored below this bound is conservative and allows these samples to be retained in the Bayesian models as left-censored data rather than discarded.

### Immunology

Due to the limited blood volume available from zebra finches (about 100 µL per individual), we were unable to use density gradient centrifugation (e.g., Histopaque) to separate leukocytes from red blood cells. Moreover, no species-specific antibodies were available to validate cell counts directly from whole blood, and several tests with both mouse and chicken antibodies showed insufficient specificity for zebra finches’ cells (Criscuolo unpublished results). Therefore, an aliquot of whole blood (50 µL), collected on heparin, was divided into two parts: 10 µL was diluted in 390 µL of cell culture media (Dulbecco Modified Eagle Medium (DMEM) high Glucose, L-Glutamine, Sodium Pyruvate, Dominique Dutcher®, Brumath, France) and used in another study (Moreno Borrallo et al. 2026b). The remaining 40 µL were centrifugated at 1000 g for 2 min at 4°C to obtain plasma containing residual white blood cells. This allowed us to remove most red blood cells, which can otherwise confound flow cytometry signals based on morphology (Seliger et al. 2012). This plasma (10 µL) was then diluted into 90 µL of cell culture media, distributed in a 96-wells plate and analysed using an Accuri^TM^ C6 flow cytometer (BD, Oxford, UK). Leukocyte populations (heterophils, lymphocytes and monocytes) were identified based on cell size (FSC channel) and shape (SCC channel), following methods described in chickens (Seliger et al. 2012) and American kestrels (Jenkins et al. 2021, see Figure 5 therein). Repeatability estimates were established on 10 repeated samples, leading to an ICC of 0.504 for H/Lratio (Boostrap repeatability estimation, P = 0.060) and of 0.704 for H:WBC (heterophil:leukocyte) ratio (Boostrap repeatability estimation, P = 0.004).

### Statistics

All analyses were conducted in R (version 4.5.1) using RStudio (R Core Team 2025). Plasma parameters were analysed using Bayesian models (*brm()* function from the brms package, Bürkner 2017, 2018) to account for censored data. Models were fitted using four chains with 2000 iterations (1000 warmup, adapt delta = 0.9), except for TBA and AST, which required a higher adapt delta (0.95). Convergence was confirmed by inspecting trace plots (see **Figure ESM2.1**), and R-hat values (all = 1, Bürkner 2017). Priors were the default priors of the brms() package: flat priors for fixed effects and weakly informative Student-t priors (default scale) for the intercept and random effects. Models included treatment (control, glucose or methylglyoxal), measurement season (November or May), sex, and all their interactions. They also included chronological age at the start of the experiment and its interaction with each of the other variables separately, and body mass at the month of the measurement. Age and body mass were mean-centred to improve interpretability (Schielzeth 2010). To explore the potential effects of selective disappearance (Forslund and Pärt 1995), the effects of the biochemical and immunological variables on mortality hazard should be determined (as we do in Moreno Borrallo et al. 2026a, b). Unfortunately, to our knowledge, no R package is currently available to handle mortality hazard analyses for explanatory variables showing both left and right analytic censoring, which is the case of many of the variables we present here. For this reason, a categorical survival variable, taking the value of 0 if the individual was alive in May and 1 if it had died, was included in all models. Although this method is not the most appropriate, it allowed us to obtain a rough estimate of the month-to-month variations attributable to deaths and those resulting from actual seasonal effects. Model selection involved a model comparison procedure using the function loo() from the eponymous package (Vehtari et al. 2017). This method involves comparing the ELPD differences relative to the best model for all models with the same number of observations, then selecting the model with the highest ELPD difference (i.e., the difference closest to 0, with differences being negative), or the simplest model if no sufficient difference is observed between the ELPD differences of multiple models (i.e., if the absolute value of a model’s ELPD difference is less than its SD, it would not be considered as to have a better predictive value). The models to be compared are the full model, which includes all the above-mentioned variables, and all of its simplified versions. Given the very large number of possible combinations, it was not feasible from a computational standpoint to test them all; therefore, the set of models was limited to those obtained through a stepwise simplification process. Specifically, each simplification step eliminated an interaction or a variable whose credible interval included zero, until the credible intervals of all remaining variables or interactions excluded zero. The models always retained at least the “treatment*month” term in order to test for season dependent effects of the treatments. Details regarding ELPD difference values for the analysed models are given in **Table ESM2.2-24** (even numbers only).

Following data inspection using histogram plots and normality tests, log_10_ transformation was applied to the dependent variables, to better approach a normal distribution, except for glucose, that already followed a Gaussian distribution. Bird ID was included as a random factor for the intercept in all models. For both Na^+^ and Ca^2+^, after data inspection, the dilution factor was included into the model, as dilution seemed to constrain their values.

For immunological data, linear mixed models (*lmer()* function from the lme4 package, Bates et al. 2015) were used, with square root transformed H/L ratio as the dependent variable. Model selection was based on information criteria using the *dredge()* function (MuMIn package, Burnham and Anderson 2002). We included the same explanatory variables as described above. We also tested their effects on leukocyte counts (heterophils, lymphocytes and monocytes), adjusting for (centred) white blood cell (WBC) counts to better characterize the nature of the underlying effects. Repeatability was calculated using the *rptGaussian()* function from the rptR package (Stoffel et al. 2017), including all the original variables (both fixed and random).

Marginal means and slopes were estimated using *emmeans()* and *emmtrends()* functions (Lenth et al. 2019) for all models (i.e. both frequentists and Bayesian). Only the equivalent comparisons were considered (e.g. between months within a treatment group or between treatment groups within a given month, but not across different months and groups at the same time, and the same for sex). Figures show all subgroups present in the final model and their pairwise comparisons, as described above, regardless of the significance of the interactions themselves. Significance was set to α < 0.05 for frequentist models, while for Bayesian models, we considered credible intervals that excluded zero as sufficient evidence of support. Structure of all final models is reported in **ESM2**.

## Results

**Table 1** summarizes raw values for all variables. No effects of the survival variable was found for any dependent variable.

**Table 1.** Summary of variables assessed (haematology: N = 117 measures, 67 in November, 50 in May; immunology: N = 200 measures). Means excluded values beyond LOQ or LOD. Sodium Na^+^: values are poorly reliable due to dilution issues.

| Variable | Mean of measured values | Minimum | Maximum | Number of values below the lower LOQ | Number of values above the upper LOQ | Number of values below the LOD | Units |
|---|---|---|---|---|---|---|---|
| **Albumin** | 1.797 | 1.044 | 3.962 | 21 | - | - | g/dL |
| **Globulin** | 1.51 | 0.614 | 3.776 | 22 | - | 22 | g/dL |
| **A/G** | 3.055 | 0.695 | 10.196 | 22 | - | 22 | - |
| **Total proteins** | 3.066 | 1.502 | 7.109 | 6 | - | 1 | g/dL |
| **AST** | 651.468 | 229.61 | 2240.628 | - | 6 | - | U/L |
| **CK** | 715.674 | 58.529 | 7807.161 | - | - | - | U/L |
| **Total bile acids** | 47.519 | 5.118 | 340.864 | 3 | - | - | µmol/L |
| **Uric acid** | 11.731 | 0.873 | 43.68 | 2 | 1 | - | mg/dL |
| **Glucose** | 300.282 | 171.65 | 438.242 | - | - | - | mg/dL |
| **Phosphates** | 5.41 | 2.005 | 19.114 | 16 | - | - | mg/dL |
| **Ca<sup>2+</sup>** | 10.367 | 6.881 | 25.5 | - | - | - | mg/dL |
| **K<sup>+</sup>** | 5.081 | 3.007 | 10.474 | 26 | - | - | mmol/L |
| **Na<sup>+</sup>** | 271.888 | 182.524 | 774.618 | 5 | - | - | mmol/L |
| **H/L ratio** | 0.28 | 0.01 | 1.38 | - | - | - | - |

### Plasma proteins

No significant effects of treatment nor of any other tested variable were observed for albumin (**Tables ESM2.1-2**). The A/G ratio was higher in November than in May for all treatments (**Figure 1.A**; Control: November – May: estimate: 0.131, HPD_95_[0.004, 0.27]; Glucose: 0.218, HPD_95_[0.049, 0.406]; Methylglyoxal: 0.17, HPD_95_[0.044, 0.298]), but no other variable affected it (**Tables ESM2.3-4**).

**Figure 1.**
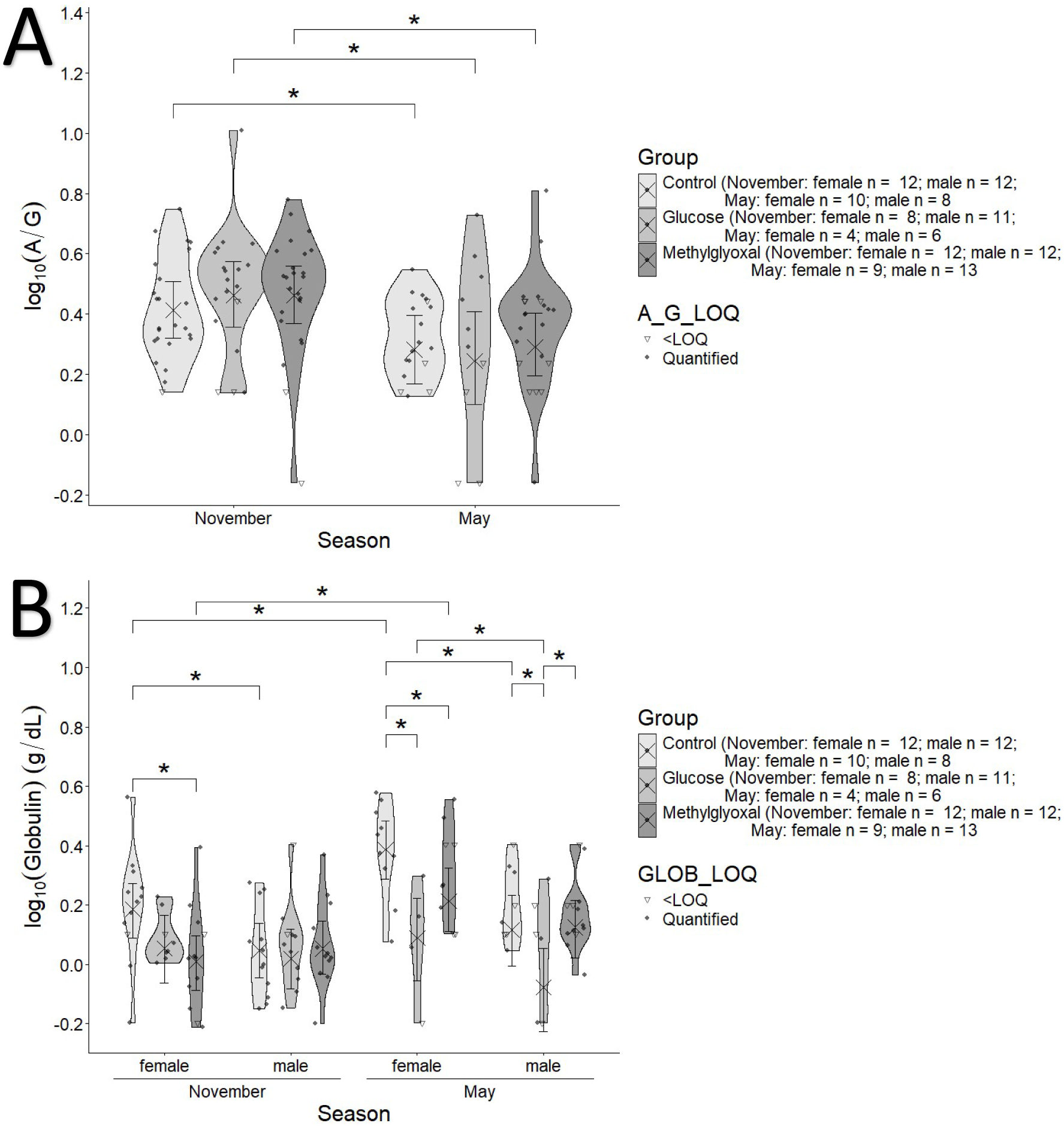
Effects of the glucose (50 g /L) and methylglyoxal (8.33 g/L) supplementation treatments, sex and seasons on (**A**) log10 transformed plasma globulin levels and (**B**) albumin to globulin ratio (A/G) of zebra finches. Filled circles represent quantified values, and empty triangles values below LOQ. Crosses and error bars represent model-estimated marginal means ±95% CI. Annotations indicating the support of the effects are based on pairwise contrasts performed separately between months within group and sex between treatments within month and sex or between sexes within season and treatment.

Globulin levels (**Table ESM2.5-6**; **Figure 1.B**) showed that glucose supplementation prevented the significant increase observed from November to May in the control and methylglyoxal treated females (control: May - November: 0.2034, HPD_95_[0.092, 0.31]; methylglyoxal: May - November: 0.204, HPD_95_[0.085, 0.327]). Indeed, in May, glucose-supplemented males had lower globulin levels than both control and methylglyoxal treated ones (Glucose - Control: -0.193, HPD_95_[-0.371, -0.03]; Glucose - Methylglyoxal: -0.204, HPD_95_[-0.373, -0.044]), while both glucose- and methylglyoxal-treated females had lower levels than control females (Glucose - Control: -0.298, HPD_95_[-0.464, -0.135]; Methylglyoxal - Control: -0.173, HPD_95_[-0.301, -0.035]). In November, control females had significantly higher globulin levels than males (female - male: 0.138, HPD_95_[0.016, 0.255]) and than their methylglyoxal counterpart (Control - Methylglyoxal: 0.175, HPD_95_[0.055, 0.3]). Moreover, females in May, showed values higher than those of males at that season for both control and glucose groups (control: female - male: 0.271, HPD_95_[0.142, 0.414]; glucose: female - male: 0.168, HPD_95_[0.0004, 0.321]).

Total plasma proteins (**Table ESM2.7-8**; **Figure 2**) followed a somewhat similar pattern, with glucose supplementation preventing the seasonal increase observed from November to May in the control and methylglyoxal groups for both sexes (**Figure 2.A.** female, Control: May - November: 0.082, HPD_95_[0.02, 0.144]; Methylglyoxal: 0.083, HPD_95_[0.024, 0.147]; male, Control: 0.082, HPD_95_[0.02, 0.144]; Methylglyoxal: 0.083, HPD_95_[0.024, 0.147]). Control females had higher total plasma protein levels than control males at both seasons (November: female - male: 0.133, HPD_95_[0.046, 0.22]; May: 0.133, HPD_95_[0.046, 0.22]), and within females, total plasma protein levels were also higher in the control group than in both the glucose- and methylglyoxal- supplemented groups at both seasons (November: Control - Glucose: 0.108, HPD_95_[0.004, 0.211]; Control - Methylglyoxal: 0.147, HPD_95_[0.053, 0.237]; May: Control - Glucose: 0.155, HPD_95_[0.038, 0.273]; Control - Methylglyoxal: 0.145, HPD_95_[0.043, 0.247]). Finally, total protein levels decreased with body mass (**Figure 2.B**; marginal slope: -0.018, HPD_95_[-0.032, -0.005]).

**Figure 2.**
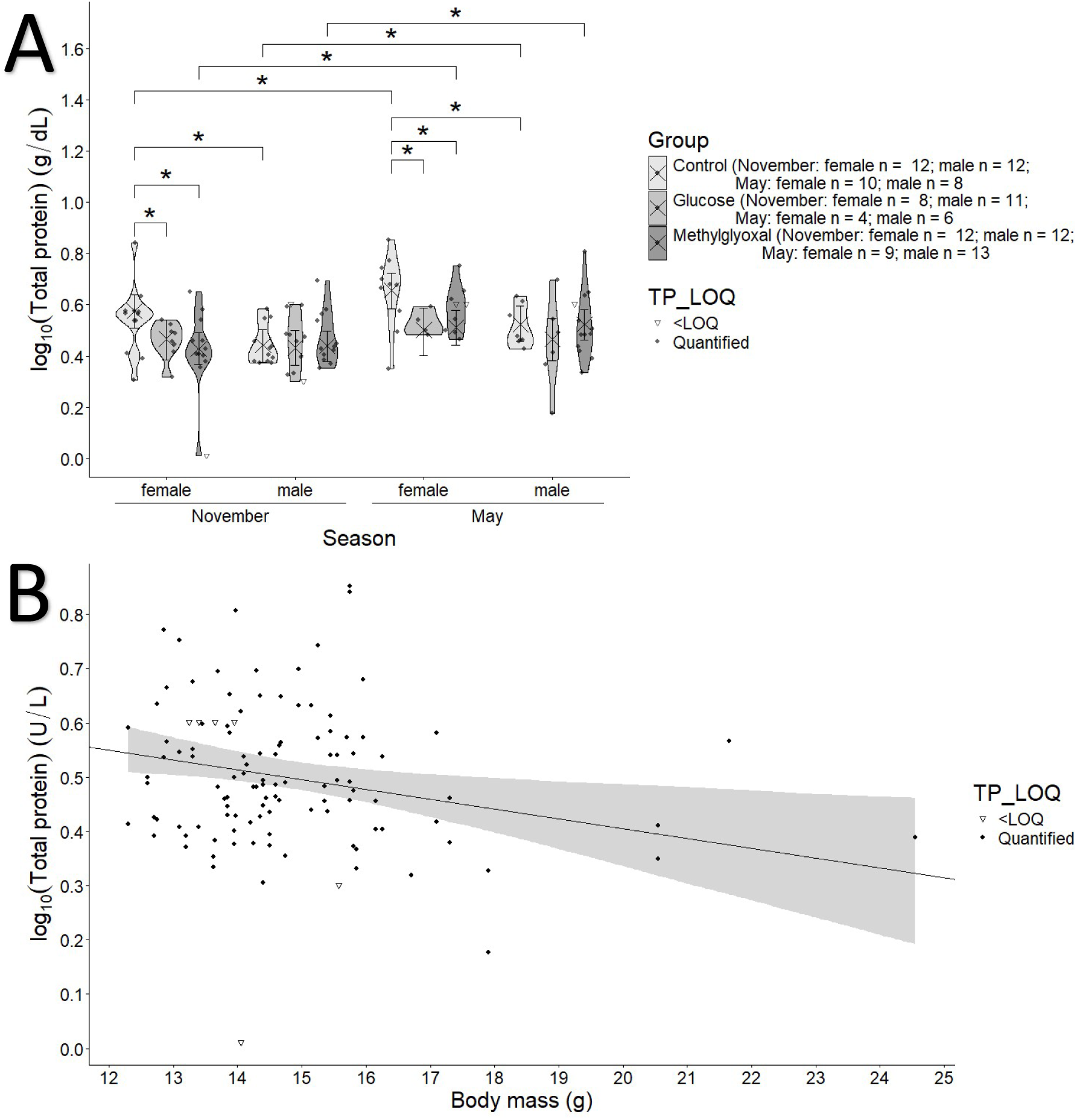
Effects of the glucose (50 g /L) and methylglyoxal (8.33 g/L) supplementation treatments across seasons and sexes (**A**) and body mass in grams (**B**) on log10 transformed plasma total protein levels of zebra finches. Filled circles represent quantified values, and empty triangles values below LOQ. Crosses and error bars represent model-estimated marginal means ±95% CI. Annotations indicating the support of the effects are based on pairwise contrasts performed separately between sexes within treatments and within seasons, between treatments within seasons and sexes or between seasons within treatments and sexes. Grey area in plot **B** represents estimated ±95% CI.

### Plasma enzymes

Plasma AST levels (**Table ESM2.9-10.**) significantly decreased from November to May for the methylglyoxal group (**Figure 3.A**; November - May: -0.216, HPD_95_[-0.325, -0.108]), reaching levels significantly lower than in both the control and glucose groups (Methylglyoxal - Control: - 0.195, HPD_95_[-0.318, -0.071]; Methylglyoxal - Glucose: -0.23, HPD_95_[-0.379, -0.093]). AST levels also decreased with age (**Figure 3.B**; marginal slope = -0.043, HPD_95_[-0.077, -0.006]).

**Figure 3.**
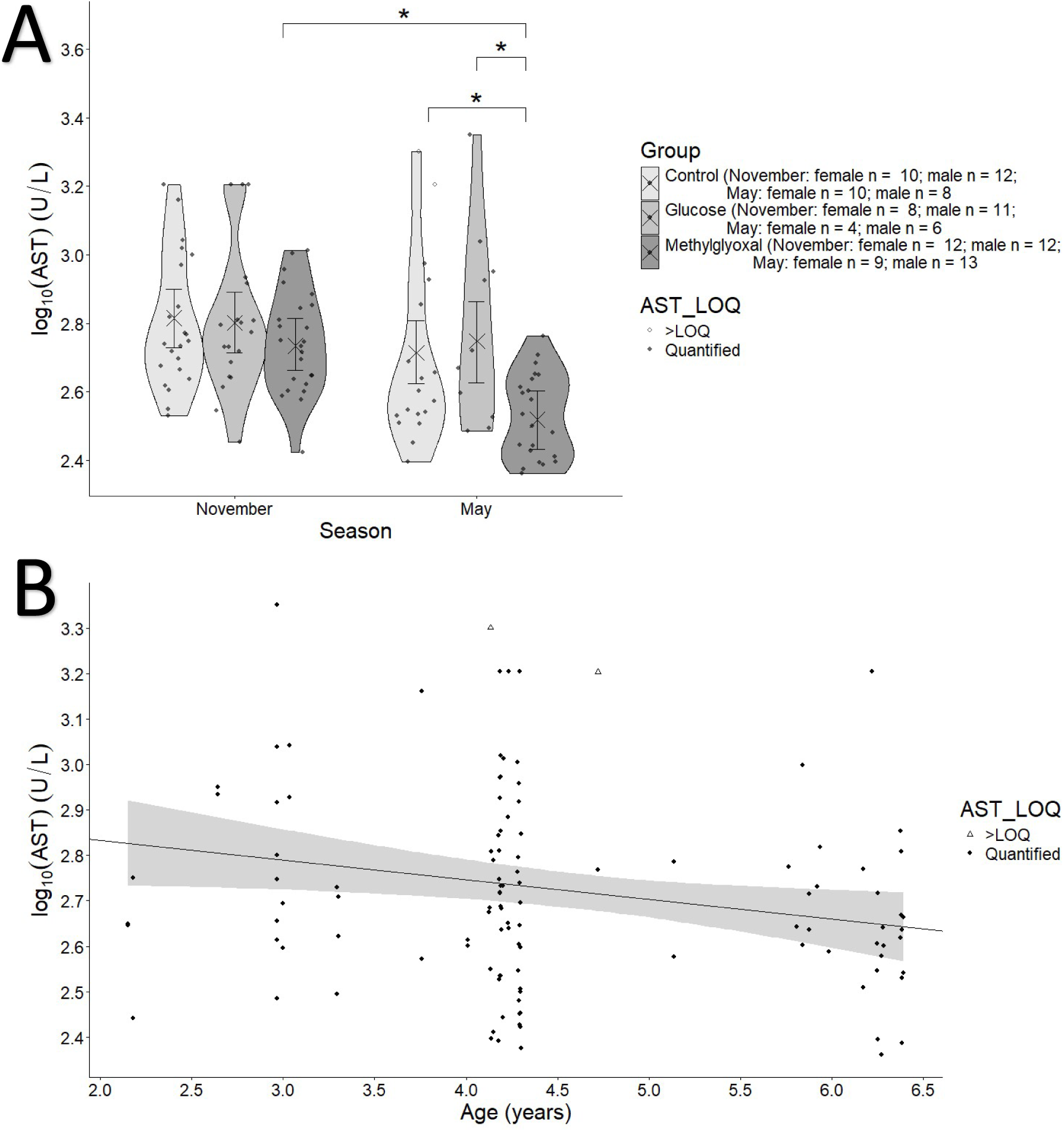
A. Effects of the glucose (50 g /L) and methylglyoxal (8.33 g/L) supplementation treatments across seasons on log10 transformed plasma aspartate aminotransferase levels of zebra finches. Crosses and error bars represent model-estimated marginal means ±95% CI. nnotations indicating the support of the effects are based on pairwise contrasts performed separately within treatments and within seasons. **C.** Variation in log10 transformed plasma aspartate aminotransferase levels with the age of the birds at the beginning of the experiment. The intercept and slope are marginalized across groups. Grey area in plot **B** represents estimated ±95% CI.

CK levels (**Table ESM2.11-12.**) significantly decreased from November to May in the methylglyoxal group (**Figure 4.A**; May - November: -0.315, HPD_95_[-0.5, -0.145]), reaching levels lower than both the control and glucose groups (Methylglyoxal - Control: -0.331, HPD_95_[-0.541, -0.127]; Methylglyoxal - Glucose: -0.267, HPD_95_[-0.511, -0.03]). CK levels also increased with age (**Figure 4.B**; marginal slope = 0.059, HPD_95_[0.002, 0.117]).

**Figure 4.**
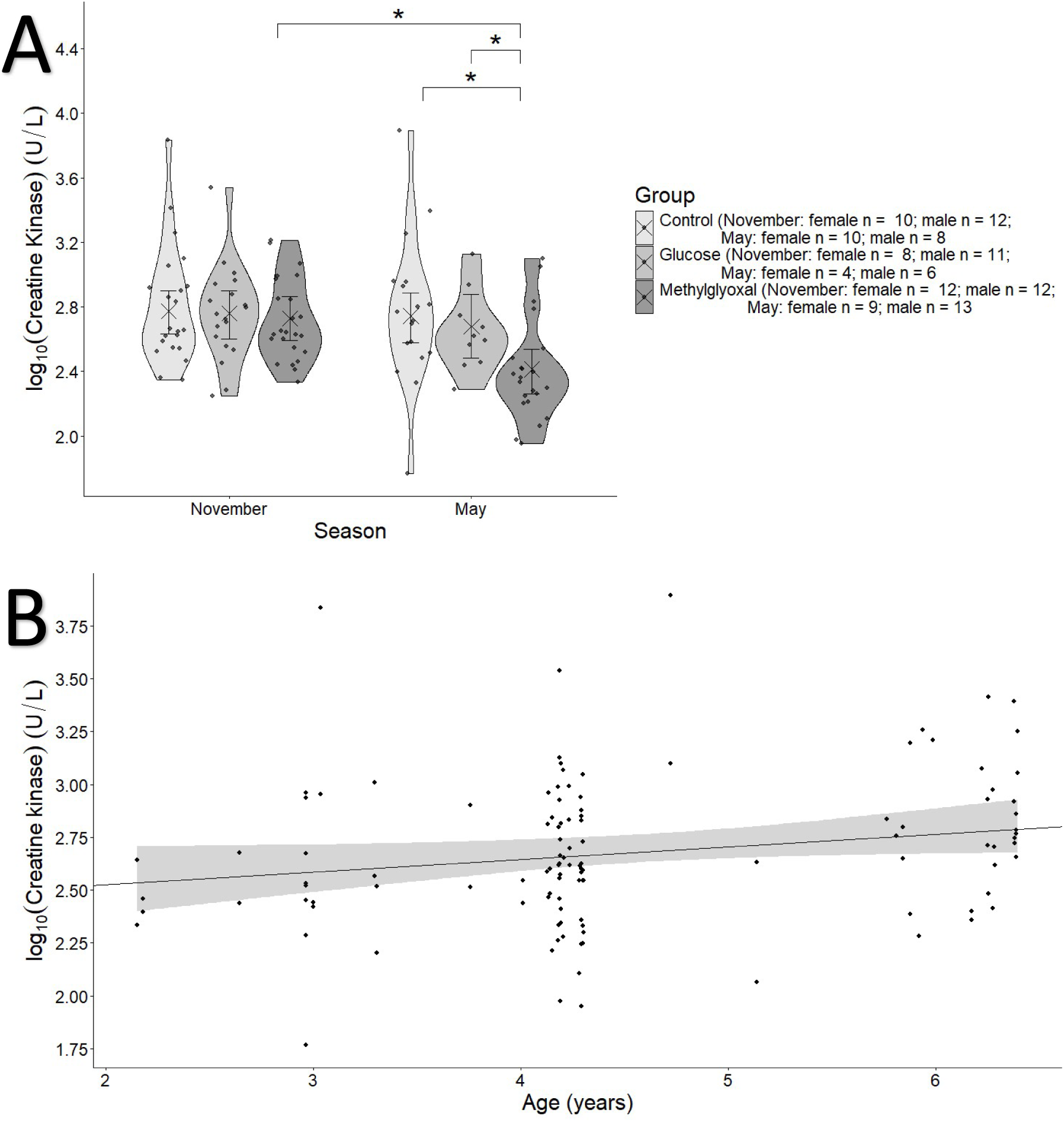
**A**. Effects of the glucose (50 g /L) and methylglyoxal (8.33 g/L) supplementation treatments across seasons on log10 transformed plasma creatine kinase levels of zebra finches. Crosses and error bars represent model-estimated marginal means ±95% CI. Annotations indicating the support of the effects are based on pairwise contrasts performed separately within seasons and within treatments. **B**. Variation in log10 transformed plasma creatine kinase levels with the age of the birds at the beginning of the experiment. Grey area in plot **B** represents estimated ±95% CI.

### Plasma glucose, uric acid and total bile acids

Plasma glucose levels (**Table ESM2.13-14.**) were significantly higher in May in both the glucose and methylglyoxal groups compared to controls (**Figure 5.A**; Glucose - Control: 49.4, HPD_95_[18.79, 83]; Methylglyoxal - Control: 70.1, HPD_95_[45.38, 98.1]), whereas in November, only the methylglyoxal group differed from controls (Methylglyoxal - Control: 34.2, HPD_95_[8.92, 58.1]). Plasma glucose positively correlated with body mass (**Figure 5.B.**; marginal slope = 6.25, HPD_95_[1.28, 11.3]).

**Figure 5.**
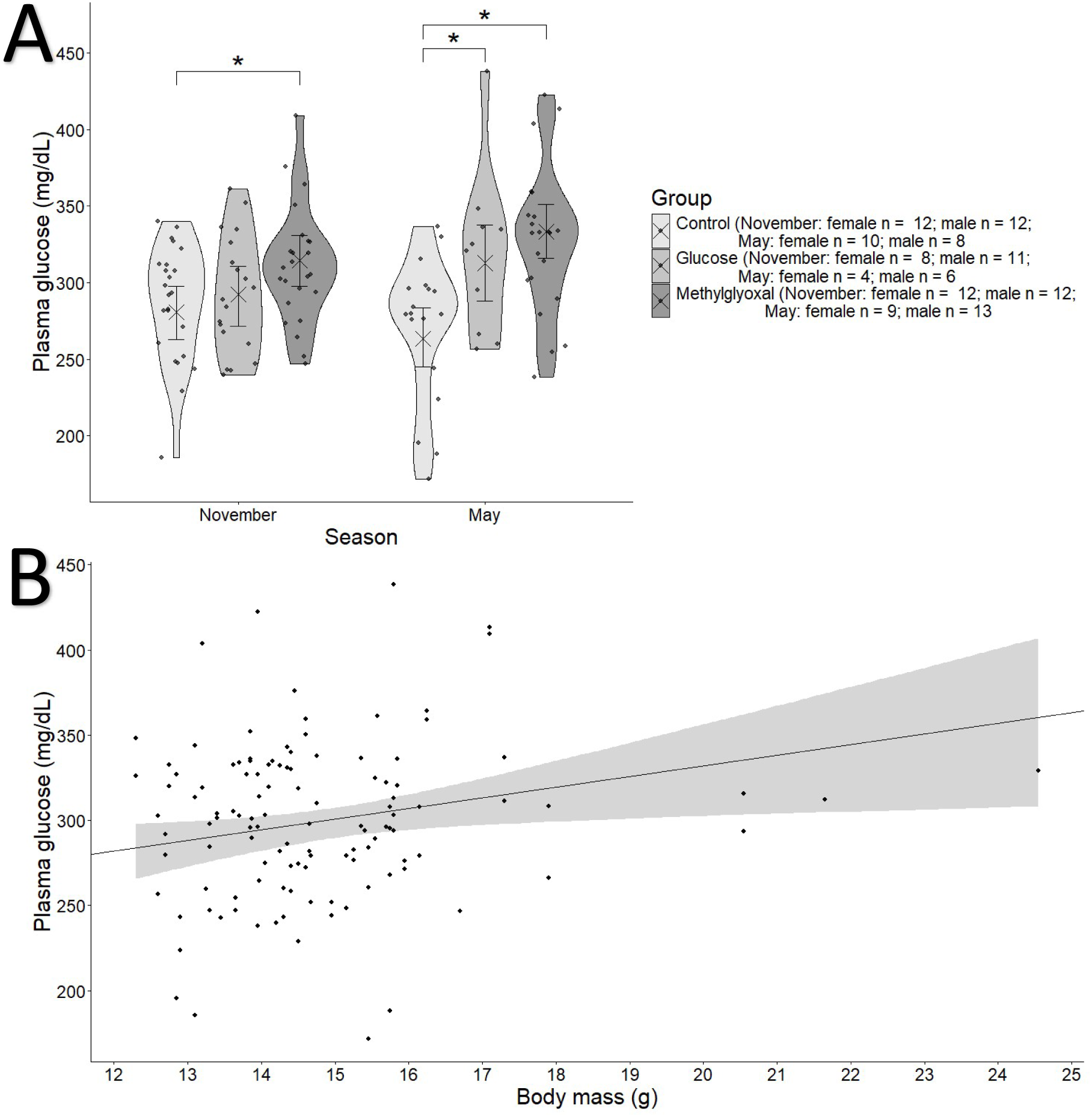
**A** Effects of the glucose (50 g /L) and methylglyoxal (8.33 g/L) supplementation treatments across seasons on plasma glucose levels of zebra finches. Crosses and error bars represent model-estimated marginal means ±95% CI. Annotations indicating the support of the effects are based on pairwise contrasts performed separately within seasons and within treatments. **B**. Variation in plasma glucose with body mass of the birds on the same season of the measure. Grey area in plot **B** represents estimated ±95% CI.

Uric acid (**Table ESM2.15-16**) decreased from November to May in methylglyoxal-supplemented females (**Figure 6.A.** May - November: -0.488, HPD_95_[-0.763, -0.234]), reaching values that were lower compared to both the control and glucose groups (; Methylglyoxal - Control: -0.688, HPD_95_[-0.983, -0.394]; Methylglyoxal - Glucose: -0.475, HPD_95_[-0.127, -0.863]). Values in females were also lower than in males within the methylglyoxal treated birds during May (female - male: -0.406, HPD_95_[-0.692, -0.143]). Uric acid also increased with age (**Figure 6.B**; marginal slope: 0.063, HPD_95_[0.007, 0.129]) and decreased with body mass (**Figure 6.C**; marginal slope: -0.054, HPD_95_[-0.1, -0.008]).

**Figure 6.**
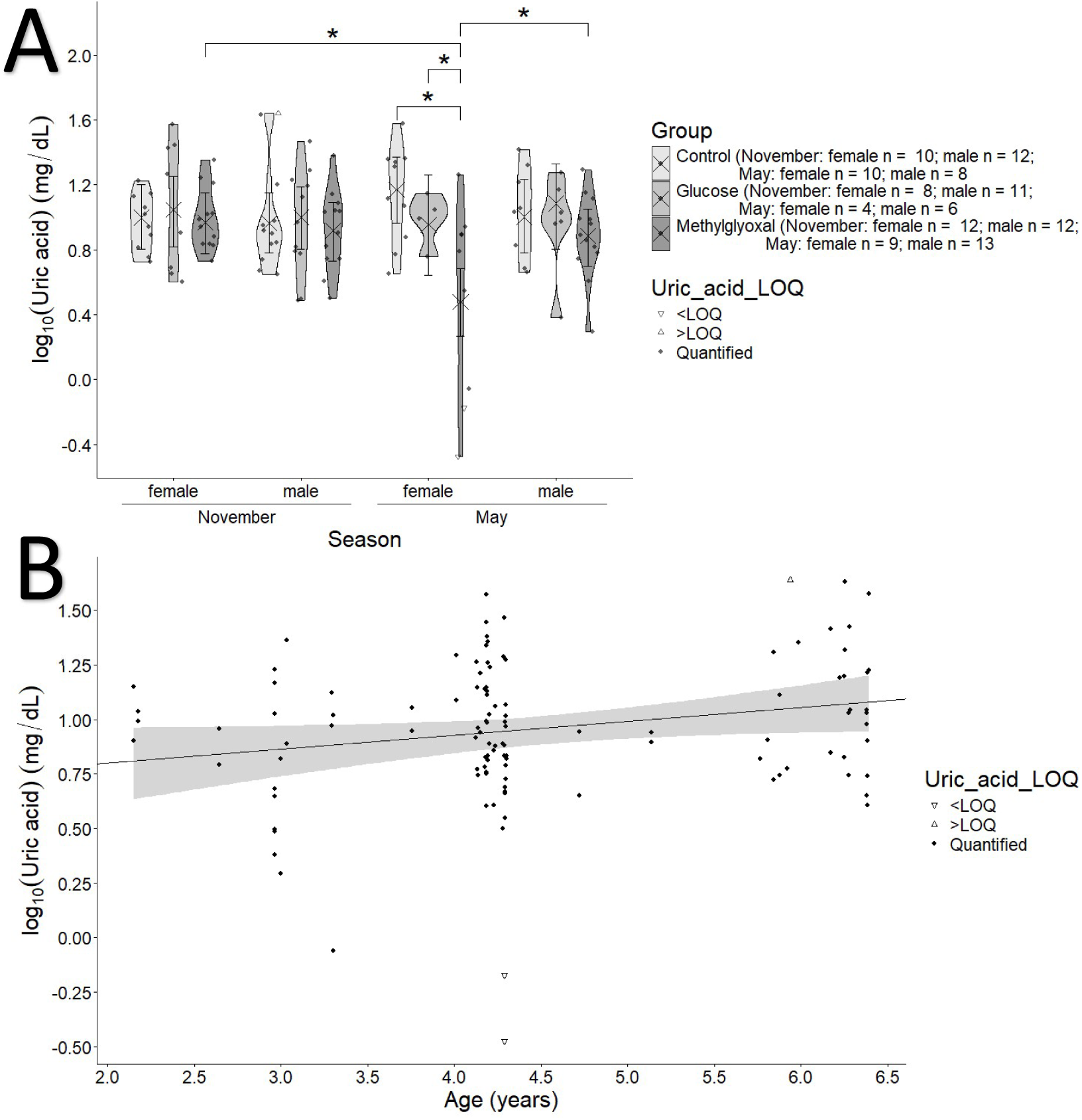
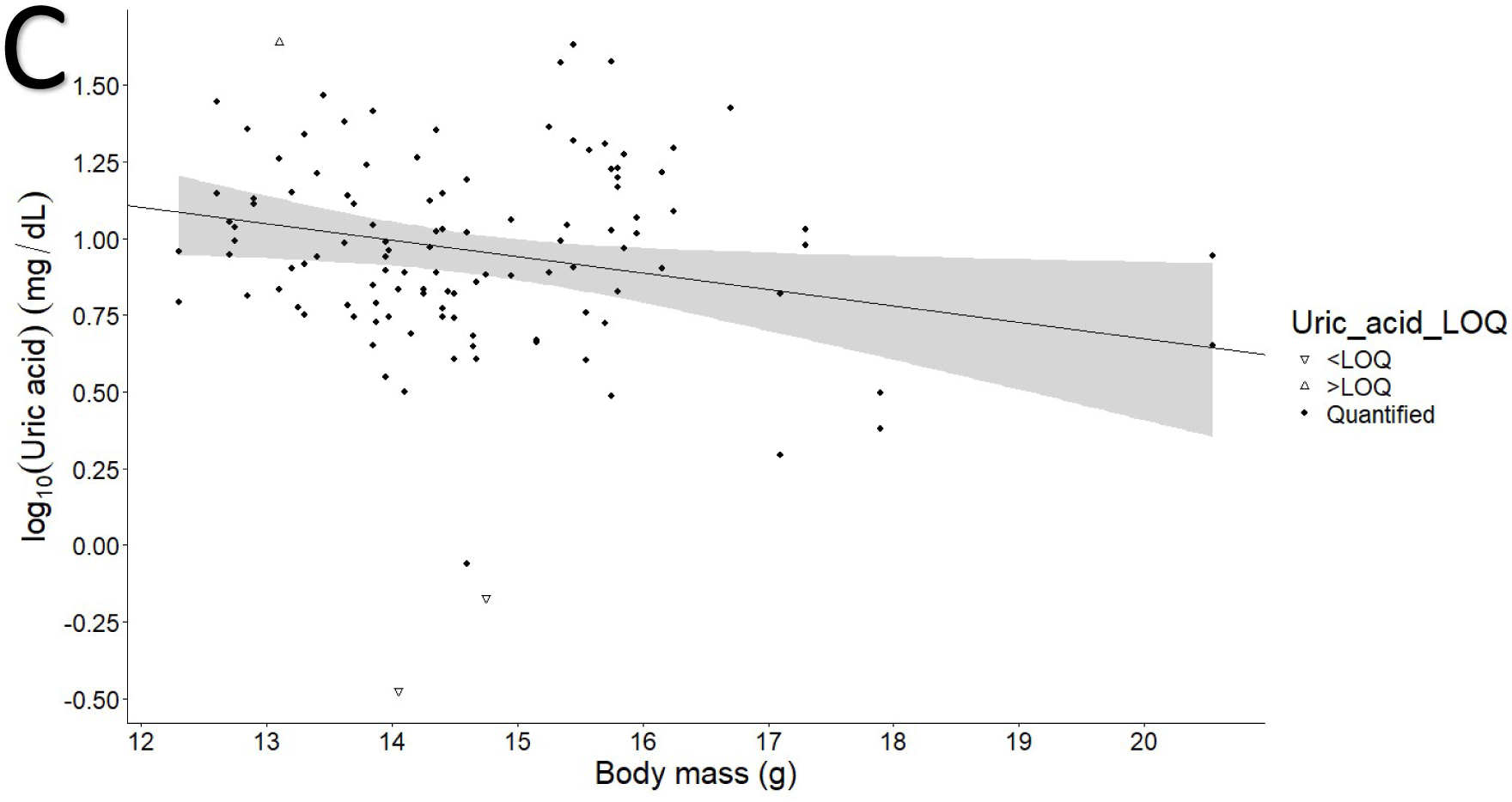
**A**. Effects of the glucose (50 g /L) and methylglyoxal (8.33 g/L) supplementation treatments across seasons and sexes on log10 transformed plasma uric acid levels of zebra finches. Crosses and error bars represent model-estimated marginal means ±95% CI. Annotations indicating the support of the effects are based on pairwise contrasts performed separately within each season and treatment, within each treatment and sex and within each season and sex. **B**. Marginal intercept and slope representing variation in log10 transformed plasma uric acid levels with the age of the birds at the beginning of the experiment. **C.** Marginal intercept and slope representing variation in log10 transformed plasma uric acid levels with the body mass of the birds at the season of the measurement. Grey area in plots **B** and **C** represent estimated ±95% CI.

Total bile acids (TBA) levels (**Table ESM2.17-18**) increased with glucose supplementation in both measurement sessions, in May, compared to both control and methylglyoxal groups (**Figure 7.A**; Glucose - Methylglyoxal: 0.327, HPD_95_[0.115, 0.546]; Glucose - Control: 0.32, HPD_95_[0.088, 0.531]). In November, this effect was observed only in comparison with the methylglyoxal group (Glucose - Methylglyoxal: 0.22, HPD_95_[0.053, 0.385]). TBA levels were significantly higher in November than in May in the control and methylglyoxal groups (Control: November - May: 0.345, HPD_95_[0.175, 0.512]; Methylglyoxal: November - May: 0.225, HPD_95_[0.081, 0.389]). TBA levels decreased with age only in the control group (**Figure 7.B**; slope: -0.135, HPD_95_[-0.21, - 0.055]).

**Figure 7.**
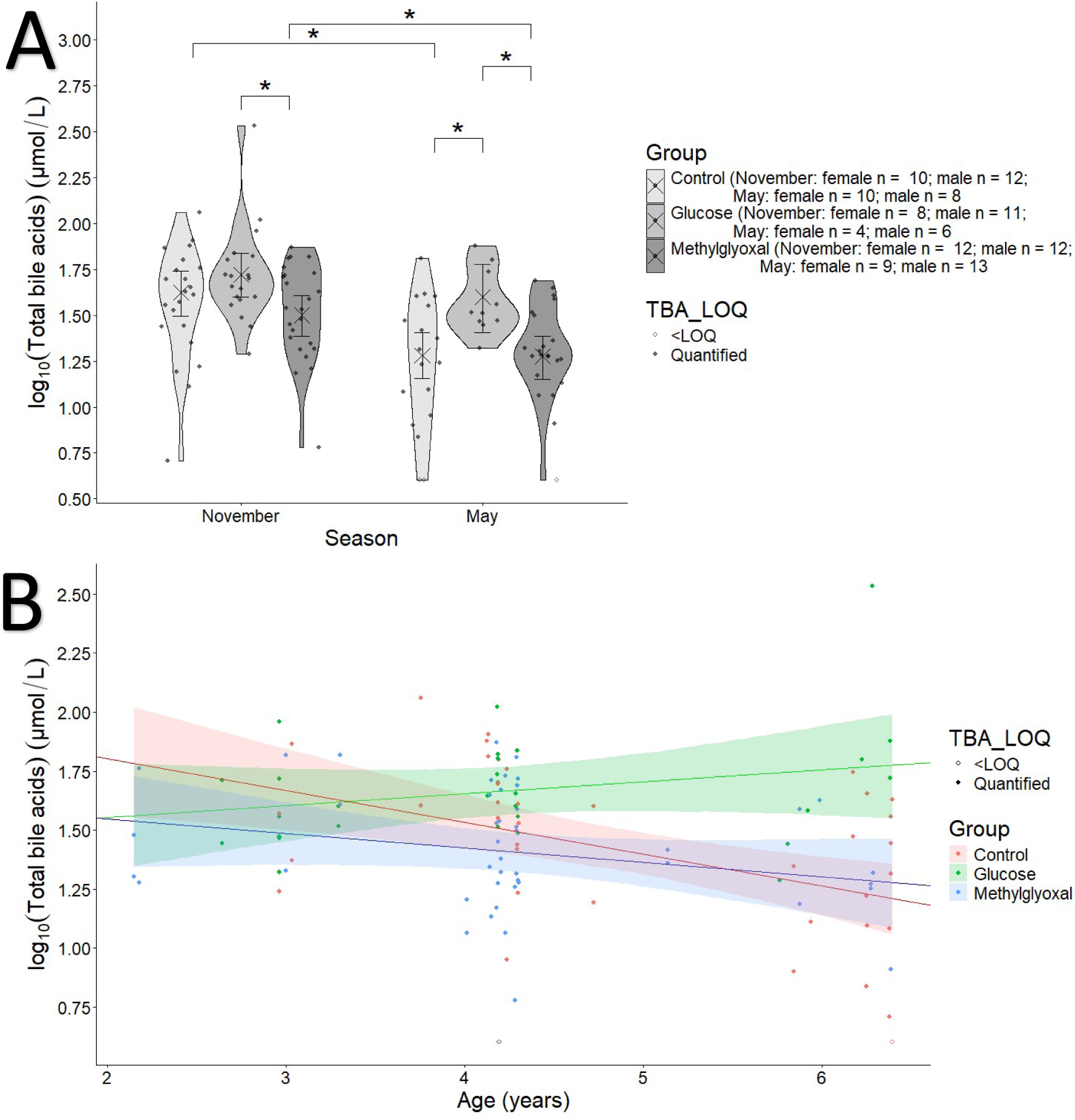
**A**. Effects of the glucose (50 g /L) and methylglyoxal (8.33 g/L) supplementation treatments across seasons on log10 transformed plasma total bile acid levels of zebra finches. Crosses and error bars represent model-estimated marginal means ±95% CI. Annotations indicating the support of the effects are based on pairwise contrasts performed separately within seasons and within treatments. **B**. Group marginal intercepts and slopes representing variation in log10 transformed plasma total bile acid levels with the age of the birds at the beginning of the experiment. Coloured areas in plot **B** represent estimated ±95% CI.

### Plasma electrolytes

Phosphate levels (**Table ESM2.19-20**) were lower in May in the methylglyoxal group compared to the control group (**Figure 8.A**; for both sexes: Methylglyoxal - Control: -0.186, HPD_95_[0.307, 0.059]). Males showed significantly lower phosphate levels in all groups in May than in November (**Figure 8.A**; May - November: Control: -0.138, HPD_95_[-0.271, -0.011]; Glucose: -0.19, HPD_95_[-0.336, -0.043]; Methylglyoxal: -0.287, HPD_95_[-0.409, -0.167]) and compared to females (**Figure 8.A**; for all treatments: male - female: -0.244, HPD_95_[-0.348, -0.128]).

**Figure 8.**
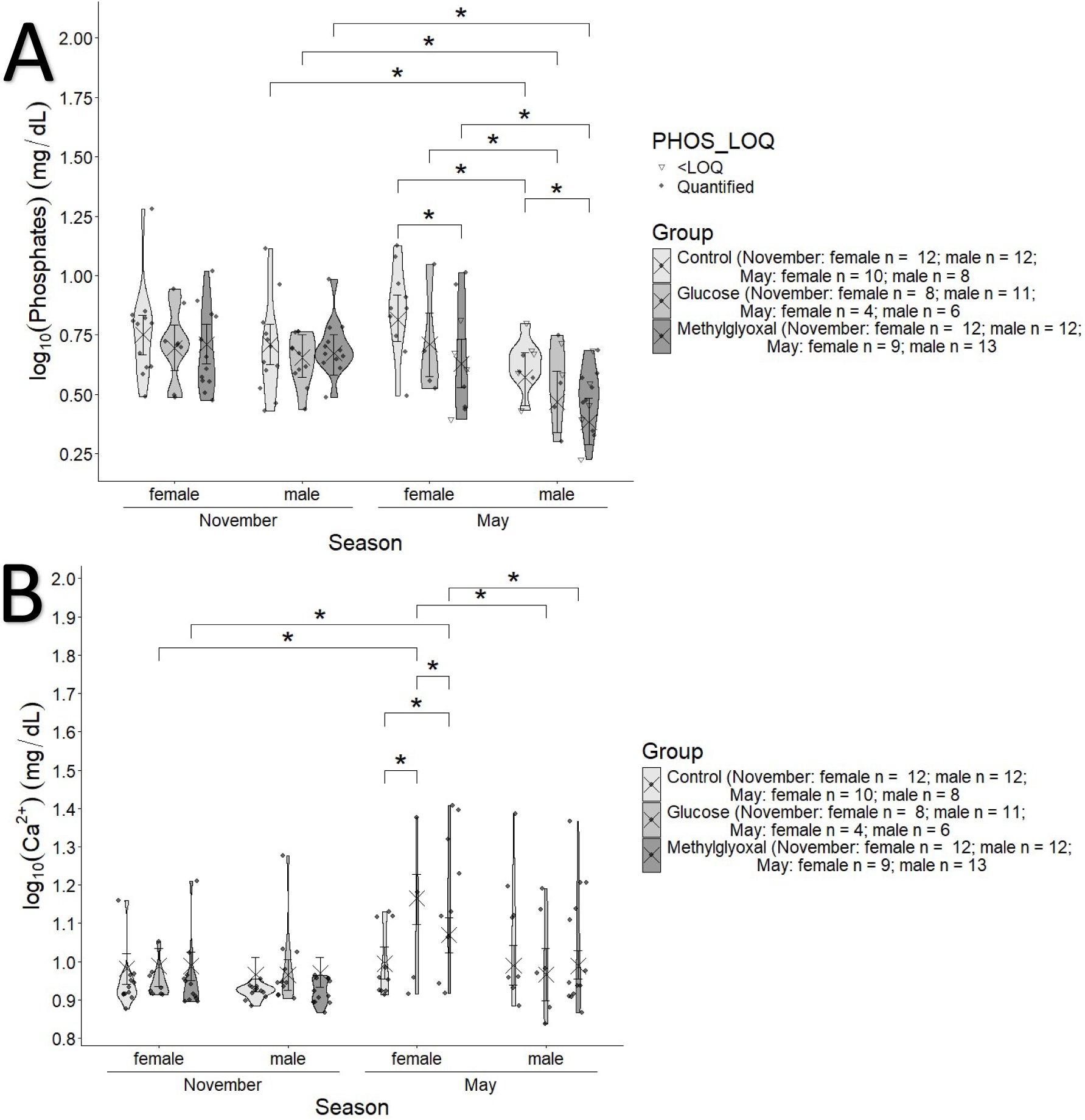
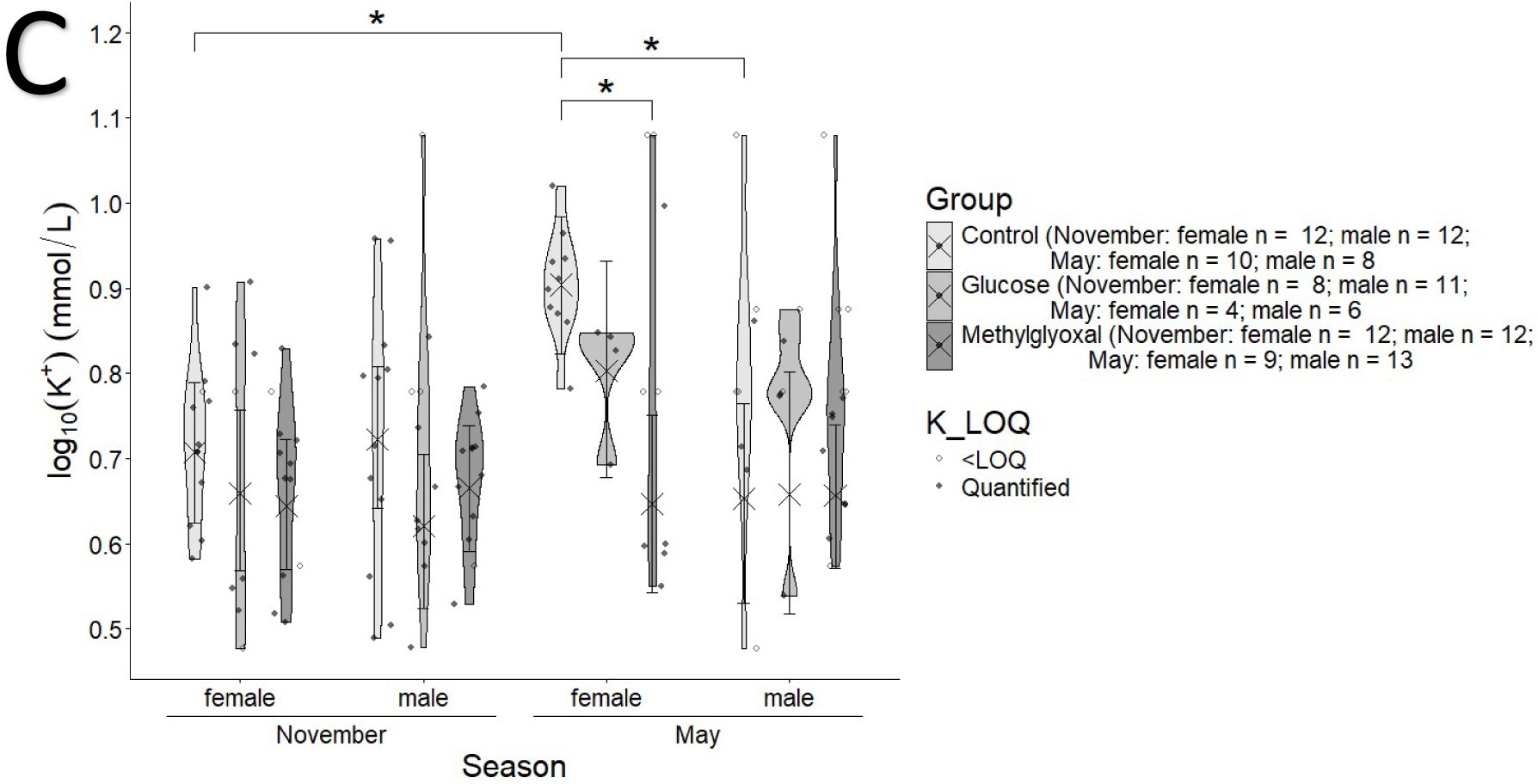
Effects of the glucose (50 g /L) and methylglyoxal (8.33 g/L) supplementation treatments across seasons and sexes on log10 transformed plasma (**A**) log10 transformed plasma phosphates levels, (**B**) calcium levels and (**C**) potassium levels of zebra finches. Crosses and error bars represent model-estimated marginal means ±95% CI. Annotations indicating the support of the effects are based on pairwise contrasts performed separately between sexes within each season and treatment, between seasons within each treatment and sex and between treatments within each season and sex.

Calcium levels (**Table ESM2.21-22**) were higher in May in females from both supplemented groups compared to controls (**Figure 8.B;** Glucose - Control: 0.164, HPD_95_[0.087, 0.24]; Methylglyoxal - Control: 0.076, HPD_95_[0.014, 0.133]) and in glucose-supplemented females in comparison to methylglyoxal (Glucose - Methylglyoxal: 0.088, HPD_95_[0.01, 0.175]). In glucose- and methylglyoxal-supplemented females, calcium levels were also higher in May than in November (May - November: Glucose: 0.177, HPD_95_[0.101, 0.252]; Methylglyoxal: 0.083, HPD_95_[0.022, 0.143]). Finally, females had higher calcium levels than males in both supplemented groups in May (Glucose: female - male: 0.175, HPD_95_[0.09, 0.258]; Methylglyoxal: female - male: 0.082, HPD_95_[0.022, 0.139]).

Potassium levels (**Table ESM2.23-24**) showed a somewhat similar but reversed sex- and season- dependent pattern, with increased values in May in control females compared to November (**Figure 8.C**; May - November: 0.198, HPD_95_[0.098, 0.295]) and males (**Figure 8.C**; female - male: 0.247, HPD_95_[0.112, 0.387]). This seasonal increase observed in females was prevented by methylglyoxal supplementation (Control - Methylglyoxal: 0.255, HPD_95_[0.129, 0.384]). For sodium levels, see **ESM2** (**Table ESM2.25-26** and **Figure ESM2.2**). For dilution effects on both calcium and sodium, see **ESM2** (**Figure ESM2.3**).

### Leukocyte profiles

The H/L ratio was significantly influenced by both supplementation type and season. Glucose supplementation led to a notable increase in the H/L ratio in May compared to both the control and methylglyoxal groups (**Figure 9.A**; Glucose - Control: marginal means contrast ± SE: 0.17 ± 0.071, t = 2.391, P = 0.046; Glucose - Methylglyoxal: 0.182 ± 0.07, t = 2.594, P = 0.027). In August, this increase was observed only relative to the control group (Glucose - Control: 0.215 ± 0.075, t = 2.866, P = 0.013). In contrast, methylglyoxal supplementation resulted in a significantly lower H/L ratio in November compared to both control and glucose groups (Methylglyoxal - Control: - 0.151 ± 0.051, t = -2.974, P = 0.009; Methylglyoxal - Glucose: -0.212 ± 0.058, t = -3.667, P = 0.001). The control group exhibited higher H/L ratios in November than in any other month (November - February: 0.254 ± 0.053, t = 4.842, P < 0.0001; November - May: 0.185 ± 0.055, t = 3.382, P = 0.005; November - August: 0.308 ± 0.055, t = 5.624, P < 0.0001). In glucose-supplemented birds, H/L ratios were higher in November compared to February (November - February: 0.256 ± 0.069, t = 3.693, P = 0.002).

**Figure 9.**
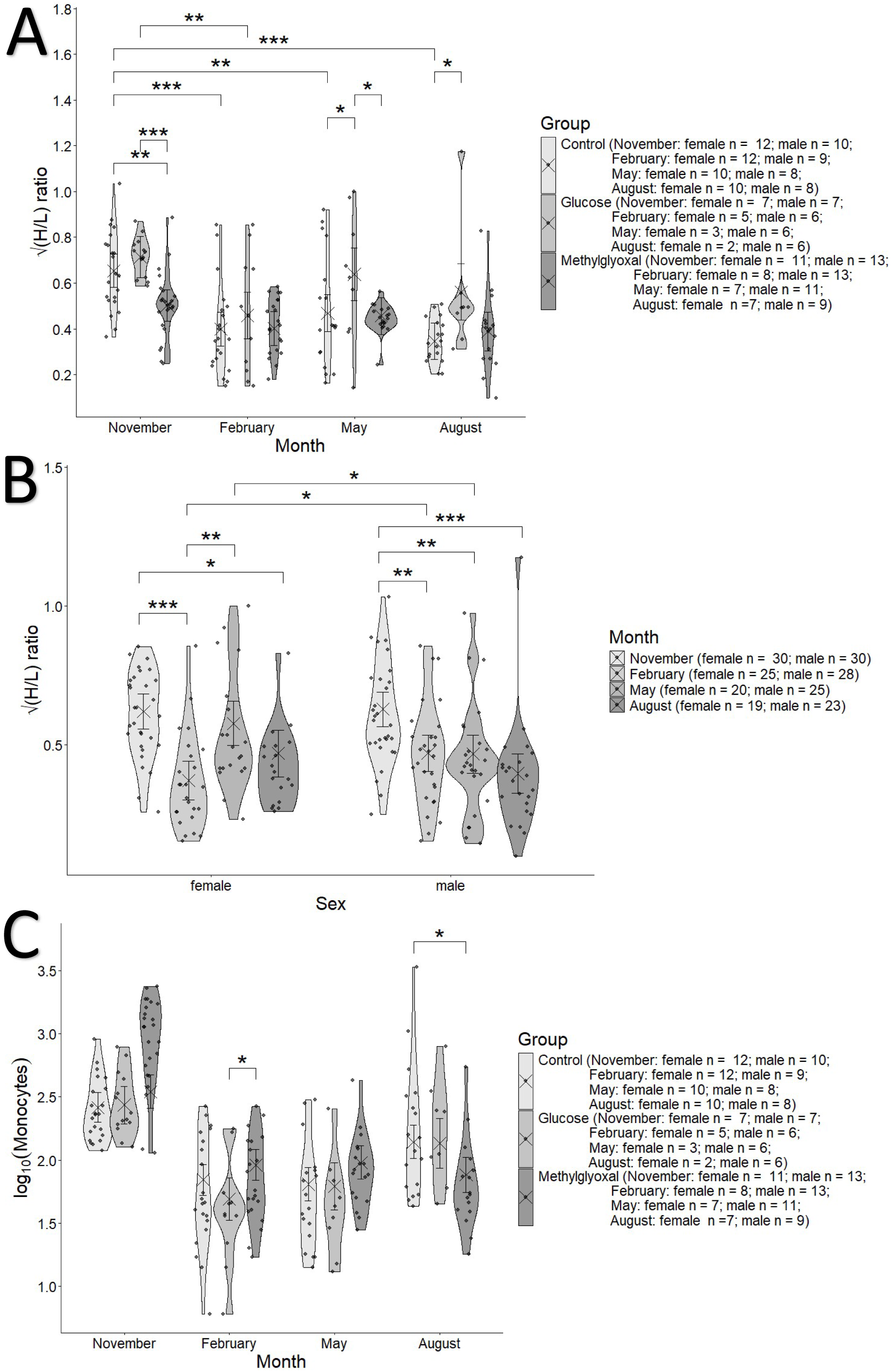
Effects of the glucose (50 g /L) and methylglyoxal (8.33 g/L) supplementation treatments across months (**A**) and sexes (**B**) on square root transformed heterophil to lymphocyte (H/L) count ratio and of the treatments across months on (**C**) the log10 transformed monocyte count (with the model adjusting for centred leukocyte count) of zebra finches. Crosses and error bars represent model-estimated marginal means ±95% CI. Significance annotations are based on pairwise contrasts performed separately within treatments (**A**, not noted for **C**) and within months (**A**, **B** and **C**) or sexes (**B**). The number of asterisks represent the level of significance (* P < 0.05; ** P < 0.01; *** P < 0.001).

Sex dependent seasonal variation in the H/L ratio was also evident (**Figure 9.B**). Males showed higher H/L ratios in November compared to all other months (November - February: 0.158 ± 0.046, t = 3.423, P = 0.004; November - May: 0.161 ± 0.047, t = 3.403, P = 0.005; November - August: 0.232 ± 0.048, t = 4.806, P < 0.0001). In females, H/L ratios were lower in February and August compared to November (February - November: -0.249 ± 0.048, t = -5.22, P < 0.0001; August - November: -0.151 ± 0.053, t = -2.833, P = 0.026). Females also differed from males depending on the month, with lower H/L ratios in February (female - male: -0.099 ± 0.048, t = - 2.057, P = 0.041) and higher values in May (female - male: 0.11 ± 0.053, t = 2.096, P = 0.037). An increase from February to May was also observed for females (May - February: 0.206 ± 0.054, t = 3.819, P = 0.001). Finally, the trait showed no repeatability (R = 0, SE = 0.039, CI_95_[0, 0.13], P = 1).

Heterophils counts (**Figure ESM2.4.A**) were not affected by treatments in November. However, in February and May, methylglyoxal supplementation increased heterophil counts compared to both control and glucose groups (February: Methylglyoxal - Control: marginal difference ± SE: 0.219 ± 0.087, t = 2.514, P = 0.034; Methylglyoxal - Glucose: marginal difference ± SE: 0.316 ± 0.105, t = 3.003, P = 0.008; May: Methylglyoxal - Control: marginal difference ± SE: 0.333 ± 0.094, t = 3.532, P = 0.001; Methylglyoxal - Glucose: marginal difference ± SE: 0.288 ± 0.116, t = 2.49, P = 0.036). In contrast, in August, methylglyoxal supplementation reduced heterophil counts compared to both other groups (Methylglyoxal - Control: marginal difference ± SE: -0.419 ± 0.098, t = -4.275, P = 0.0001; Methylglyoxal - Glucose: -0.744 ± 0.123, t = -6.027, P < 0.0001), whereas glucose supplementation increased heterophil counts compared to controls (Glucose - Control: 0.325 ± 0.12, t = 2.696, P = 0.021). Seasonal effects were also seen (see **ESM2** for details). Heterophil counts were positively associated with white blood cell (WBC) count (effect ± SE: 1.028*10^-4^ ± 1.264*10^-5^, t = 8.127, P = 4.5*10^-14^). The trait showed no repeatability (R = 0, SE = 0, CI_95_ = [0, 0], P = 1).

Lymphocyte counts (**Figure ESM2.4.B**) were unaffected by the treatments in November. In February, methylglyoxal supplementation increased lymphocyte counts compared to glucose- supplemented birds (Methylglyoxal - Glucose: 0.32 ± 0.099, t = 3.219, P = 0.004). In May, methylglyoxal supplementation also increased lymphocyte counts compared to both control and glucose groups (Methylglyoxal - Control: 0.275 ± 0.089, t = 3.079, P = 0.007; Methylglyoxal - Glucose: 0.437 ± 0.11; t = 3.995, P = 0.0003). In contrast, in August, methylglyoxal supplementation decreased lymphocyte counts compared to both control and glucose groups (Methylglyoxal - Control: -0.381 ± 0.093, t = -4.112, P = 0.0002; Methylglyoxal - Glucose: -0.354 ± 0.117, t = -3.037, P = 0.007). Seasonal effects were also seen (see **ESM2** for details). Lymphocyte counts were positively correlated with WBC count (estimate ± SE: 1.735*10^-4^ ± 1.195*10^-5^, t = 14.516, P < 2*10^-16^), but the trait showed no repeatability (R = 0, SE = 0, CI_95_ = [0, 0], P = 1).

Monocyte counts (**Figure 9.C**) were influenced by methylglyoxal supplementation in a season- dependant manner. In February, methylglyoxal indeed increased monocyte count compared to glucose-supplemented birds (Methylglyoxal - Glucose: 0.272 ± 0.105, t = 2.588, P = 0.028). In contrast, in August, methylglyoxal decreased monocyte counts compared to controls (Methylglyoxal - Control: -0.26 ± 0.098, t = -2.657, P = 0.023). Seasonal effects were also seen (see **ESM2** for details). Monocyte counts were positively correlated with total leukocyte counts (estimate ± SE: 1.563*10^-4^ ± 1.257*10^-5^, t = 12.432, P < 2*10^-16^). However, repeatability of monocyte counts was not significant (R = 0.036, SE = 0.05, CI_95_[0, 0.168], P = 0.274).

## Discussion

### Glucose upplementation prevents the seasonal plasma protein increase

Many of the assessed parameters showed sex dependent effects with seasonal variations. Total plasma protein and globulin levels notably followed similar patterns: control females exhibited higher levels than males, whereas this sex difference was abolished by both glucose and methylglyoxal supplementations in November. We also observed a May increase in both total plasma protein and globulin levels in comparison to November, which was prevented by glucose supplementation. The observed similar variation in globulin and total plasma protein levels likely derives from methodological characteristics of the veterinary analyser technique, which calculates the globulin fraction simply by subtracting albumin from total protein concentration. Given that albumin levels - one of the main plasma proteins - remained unaffected by our treatments, globulin fraction, as defined by the device, drive most plasma protein variations in our samples. Indeed, this seasonal increase in globulin levels likely explains the decrease in A/G from November to May, potentially reflecting the higher stress (Harr 2002; Davis et al. 2008, 2018) suffered by the birds in November (see also Moreno Borrallo et al. 2026a, b). The observed seasonal increase in the plasma protein levels variables, which at least for the globulin variable appears restricted to females, may reflect reproductive physiology. In birds, reproductive activity is associated with oestrogen-mediated changes in hepatic protein synthesis and plasma protein composition (Walzem 1996). This interpretation is further supported by the egg production and nest-building behaviours observed during that period. Similar patterns of elevated plasma proteins in prelaying or preovulatory females have been observed in American kestrels (*Falco sparverius*; Dawson and Bortolotti 1997), pigeons (*Columba livia*; Gayathri et al. 2004; Gayathri and Hegde 2006) and great tits (*Parus major*; Hõrak et al. 1998) followed by a decrease during incubation.

The mechanism behind supplementation-induced reduction in protein levels remains unclear. Sugar syrup supplementation in chickens shows no effect (Hussein et al. 2016) or a dose- dependent increase in total plasma proteins (Hussein et al. 2018). In these chickens (i.e. Hussein et al. 2016), supplementation was provided within the food itself and was associated with increased food intake. Our results may therefore be explained by reduced food intake in birds receiving glucose in drinking water (see Moreno Borrallo et al. 2026a).

### Supplementation effects on plasma electrolytes suggest modulation of reproductive physiology through alterations on energy balance

Both glucose and methylglyoxal supplementation increased calcium levels in females in May, with a stronger effect of glucose supplementation. This suggests that the experimentally induced increase in plasma glucose may have promoted reproductive investment in females, consistent with the opportunistic breeding strategy of zebra finches, which is partly driven by nutritional status (see e.g. Perfito et al. 2007; Ruffino et al. 2014; Prabhat et al. 2020). The resulting calcium increase would support eggshell formation (Sinclair-Black et al. 2023). However, given the *ad libitum* feeding conditions of the birds, it remains unclear why the control group did not already show physiological changes in May associated with reproductive activity, even if at lower magnitude than in the supplemented groups.

Plasma potassium increased in control females in May, although the underlying mechanism remains unclear. This effect was also attenuated by glucose and prevented by methylglyoxal supplementation. Potassium levels can be influenced by acid base balance, including the mild alkalosis associated with egg formation, as well as by changes in plasma proteins or lipids. In this context, oestrogen-induced lipemia during egg production could explain the observed electrolyte shifts in treated birds (see overview in Jones 1999). However, the effects of carbohydrate metabolism on plasma electrolyte regulation remain poorly understood, especially in passerines.

Methylglyoxal supplementation reduced plasma phosphate levels in May. This may be related to an impairment of energy metabolism induced by methylglyoxal, as hypophosphatemia is known to be associated with insulin resistance in some mammal species (see e.g. Pollack et al. 1934; DeFronzo and Lang 1980; Zhou et al. 1991), although its applicability to birds remains unclear. However, the observed sex and season-dependent effects are difficult to explain. It is finally worth noting that we detected a significant positive effect of the dilution factor on both sodium and calcium levels. Although for calcium the higher dilution factor only avoided detecting levels normally detected at lower dilution levels, for sodium, nevertheless, higher dilution actually increased its values, indicating that the reverse osmosis water was not sufficiently pure in sodium, and thus rendering these results unreliable.

### Glucose increases liver damage, while methylglyoxal paradoxically reduces plasma cell damage markers: hormesis or chronic disease?

TBA levels were higher in November in the control and methylglyoxal groups than in May. This phenomenon may be related to the more severe damage suffered by the birds during this period, including increased apoptosis of blood cells and higher production of mitochondrial superoxide in non-apoptotic blood cells, accompanied by higher mortality, as previously reported (Moreno Borrallo et al. 2026a, b). However, glucose supplementation led to an increase in TBA levels in May compared to the two other groups, and only compared to methylglyoxal-supplemented birds in November, suggesting glucose may induce hepatic damage more than methylglyoxal does.

Nevertheless, contrary to our expectations, methylglyoxal supplementation reduced plasma markers of cellular damage—aspartate aminotransferase (AST) and creatine kinase (CK)—in May. This contrasts with evidence from mammals, where methylglyoxal has been associated with hepatic damage (Dutta et al. 2026). One possible explanation is that the lower AST and CK levels observed in methylglyoxal-supplemented birds in May could reflect a reduced organ mass in this treatment group related to chronic pathological conditions, (e.g. sarcopenia or cirrhosis), as discussed previously in birds (Lumeij 1987, 1997). Nevertheless, appropriate species-specific pathology-related references would be needed to confirm this interpretation. Interestingly, lower plasma protein and uric acid levels in methylglyoxal-treated females could be associated with alterations protein metabolism. Alternatively, reductions in plasma uric acid levels in methylglyoxal-treated females may be caused by its use as an antioxidant compound (Simoyi et al. 2003; Tsahar et al. 2006). However, this would not explain why this is not observed in males that show higher protein carbonylation levels (Moreno Borrallo et a. 2026b). Finally, the negative and positive correlations between uric acid and body mass and age, respectively, may reflect a general negative relationship between uric acid levels and body condition, possibly linked to increased protein catabolism with age. That said, previous studies conducted on zebra finches (Zagkle et al. 2020; Levillain et al. 2026; Płachta et al. 2026) have not shown any effects of age on uric acid levels, although the variable was often treated as categorical, and its variation with body mass have been explored even less. Together with the limited statistical evidence provided by our own study regarding these relationships, these observations call for caution in interpreting such results.

A second possible explanation for the observed reduction in AST and CK values is a hormetic response, whereby low-level exposure to methylglyoxal triggers adaptive mechanisms that mitigate its cytotoxic effects (Calabrese, 2007; Costantini et al., 2010). Indeed, methylglyoxal has been proposed to participate in cell signalling pathways that induce protective responses against glycation stress (reviewed in Vašková et al. 2025). Mechanistically, such protection could involve enhanced antioxidant defences and/or increased methylglyoxal metabolism and clearance. While methylglyoxal can activate Nrf2 (a transcription factor inducing certain antioxidant defences) in mammals (Castiglione et al. 2020), Neoaves have constitutively active Nrf2 due to modifications in KEAP1, a protein that inhibits Nrf2 activity (Bollong et al. 2018). Part of this KEAP1 modification is present in protein regions that are modified by reactive compounds like methylglyoxal, so this pathway may operate differently in Neoaves, and currently available data suggests that it not likely to be amenable to methylglyoxal response, although ultimately it must be said thatNrf2 activity was not measured in this study. We suggest, however, that Nrf2 independent methylglyoxal inducible defensive pathways may still underlie the observed response. However, this general hormetic interpretation requires further validation in our system, particularly given that the same treatment increased oxidative stress and apoptosis in blood cells in zebra finches (Moreno Borrallo et al., 2026b). Alternatively, in line with the oxidative shielding hypothesis (Blount et al. 2015), the observed reduction in plasma damage markers with methylglyoxal supplementation might reflect an indirect activation of antioxidant defences in reproductive females, since methylglyoxal may have stimulated reproductive activity (as suggested by increased calcium levels, see above) Nevertheless, this hypothesis would also predict the same observation in the glucose group, although methylglyoxal more consistently increased plasma glucose levels, perhaps explaining a stronger effect.

Finally, we observed age-related decreases in plasma AST and TBA (only in the control group for the latter). These age-related decreases could be an effect of selective disappearance (Forslund and Pärt 1995), with individuals with higher hepatic damage levels having a higher probability of dying. We however observed an age-related CK increase, possibly reflecting increase in muscle damage. These patterns align with previous reports of cross-sectional age-related differences in liver and muscle metabolic function in zebra finches (Salmón et al. 2022), showing that liver ROS production per unit of oxygen consumption increases with age, whereas muscle does not, potentially at the cost of reduced ATP production efficiency. Nevertheless, the “old” birds (4 years) in that previous study are closer in age to our “young” cohort, and thus potential changes occurring at more advanced ages (e.g. ≥6 years) remain unexplored. Overall, these results suggest that liver dysfunction may have a greater impact on survival at very advanced ages, whereas muscle-related changes might be less critical in our study population. This interpretation is supported by the fact that birds had constant access to food and water in the aviary, and by the observation that flying performance—although partially affected by the treatments—did not appear to influence mortality risk in this group (Moreno Borrallo et al. 2026a).

### Supplementations induce season-dependent modulation of immune function

Glucose supplementation increased the H/L ratio in May and August, suggesting elevated physiological stress, although values remained within the physiological range reported for birds (Clark 2014). This pattern may be related to the potential of glucose supplementation to reduce food intake (Moreno Borrallo et al. 2026a), thereby modulating stress responses (Birkhead and Fletcher 1998). It may also reflect the known negative association between H/L ratio and body condition indexes (Birkhead and Fletcher 1998; Colominas-Ciuró et al. 2024; but see Ewenson et al. 2001 for a more limited relationship). Conversely, methylglyoxal reduced the H/L ratio in November, when baseline stress was elevated, suggesting a context-dependent modulation of stress physiology. This is consistent with its previously reported reduction of apoptosis during this high-stress period (Moreno Borrallo et al. 2026b). Thus, methylglyoxal effects may depend on the initial physiological state, with different outcomes when certain stress markers are already elevated.

Absolute cell counts provide further insight into treatment effects on immune cell dynamics, helping to clarify patterns observed in the H/L ratio. Methylglyoxal increased heterophils and lymphocytes in February and May but decreased both in August, whereas glucose only increased heterophils in August. These compensatory shifts help explaining why methylglyoxal’s effects on H/L ratio were not always evident when considering cell proportions alone. The May increase in H/L ratio in glucose-supplemented birds, despite unchanged absolute cell counts, suggests subtle proportional shifts in immune cell populations that are not fully captured by relative cell measures alone. Finally, compared to glucose, methylglyoxal increased monocyte counts in February, i.e., the season when monocyte counts are at their lowest and decreased them in August compared to control, i.e., a season when monocyte counts are higher. This pattern may suggest a tendency towards regression to the mean in monocyte responses under methylglyoxal supplementation.

Seasonal H/L ratio fluctuations could be related to temperature, as previous studies in zebra finches have shown increases in H/L ratio under both chronic high and low temperature exposure (Colominas-Ciuró et al. 2024; Udino et al. 2024a; Udino et al. 2024b), but not under acute heat exposure (Xie et al. 2017). However, this would predict higher ratios in February, when ambient temperatures were lowest (**ESM3**) and well below the thermoneutral zone for zebra finches (Calder 1964; Wojciechowski et al. 2021). Therefore, additional regulatory factors should be involved in our case. For example, the female-specific increase in H/L ratio in May could be linked to reproductive demands, as previously reported in great tits (*Parus major*) (reviewed in Skwarska 2018). Seasonal variability in this species also parallels our findings in zebra finches, with reduced immunity and H/L ratios in winter and increases in spring and summer (reviewed in Skwarska 2018).

### Glucose and especially methylglyoxal supplementation induce plasma glucose increase, but measuring technique matters

Both supplementations effectively increased plasma glucose levels in May. In November, this effect was observed only in the methylglyoxal group, indicating a stronger and more consistent response to methylglyoxal across sampling periods compared to glucose. This confirms the efficacy of our supplementation protocol. Notably, the impact of methylglyoxal on avian glycemia had not been previously documented, although it has been shown to increase hepatic gluconeogenesis in mice (Dutta et al. 2026), likely from lactate, one of the main products of its metabolism (reviewed in e.g. Vašková et al. 2025). Furthermore, glucose supplementation does not consistently affect blood glucose levels in birds (Basile et al. 2022). Accordingly, we observed no changes in whole blood glucose (Moreno Borrallo et al. 2026a), suggesting that plasma glucose measurements are more sensitive than whole blood glucose for detecting treatment-induced changes in glycemia. This is likely the case because whole blood glucose may be buffered by cell-associated glucose utilization and haematocrit changes. However, what is most surprising is that in our previous manuscript (Moreno Borrallo et al. 2026a), we found no effect of the treatments on plasma glucose levels (assessed in a subgroup of birds at different months), nor did we find the positive correlation between plasma glucose levels and body mass that is currently being reported.

Detection of supplementation effects may depend on measurement technique. We therefore also measured plasma glucose levels with a portable glucometer in most samples from May (48 out of 50), in parallel with the veterinary analyser. In a previous study, glucometer readings had been shown to correlate well with a colorimetric glucose assay kit, using plasma samples from several bird species across a wide range of glucose levels (Moreno Borrallo et al. 2025). Here, glucometer readings were consistently higher than those from the veterinary analyser, with greater discrepancies at lower concentrations: the glucometer overestimated plasma glucose by 83.4 mg/dL at the average veterinary analyser value in this dataset. A linear model comparing centred veterinary analyser values to glucometer values showed a significant intercept (α ± SE: 390.646 ± 6.618 mg/dL) and slope (β ± SE: 0.376 ± 0.123, P = 0.004), indicating a systematic bias between methods. This methodological difference may explain why no group differences in plasma glucose levels were previously detected (Moreno Borrallo et al. 2026a). Therefore, caution should be exercised in future studies when comparing glucose concentration data obtained using different analytical techniques. Indeed, adequate species-specific validation is also recommended. For now, studies trying to validate the use of point-of-care devices designed for humans or veterinary use (normally for mammals) on many different species, including birds (Stoot et al. 2014) have rendered mixed evidence of their adequacy, with variation depending on species and brand used (see e.g. for birds Acierno et al. 2009; Acierno et al. 2012; Anderson et al. 2022; Hiebert et al. 2022; Mohsenzadeh et al. 2014).

## Conclusion

This study explores how chronic glucose and methylglyoxal supplementation reshapes physiological regulation in zebra finches in a strongly sex- and season-dependent manner, particularly suggesting an alteration of reproductive-related metabolism, and differential damage effects of glucose and methylglyoxal. Indeed, supplementation also modified plasma glucose, electrolyte balance and immune traits, indicating that glucose-derived metabolites influence multiple physiological systems simultaneously.

While glucose supplementation elevated stress markers (H/L ratio) during the reproductive and summer seasons, and potentially induced hepatic function impairment (as indicated by increased total bile acid levels), methylglyoxal exhibited paradoxical effects on tissue integrity markers, despite its pro-oxidative properties, possibly through hormetic mechanisms. Nevertheless, this hypothesis would require more precise experimental validation with different dosages. Other effects of methylglyoxal suggest that it may disrupt protein dynamics and that it potentially interfere with energy metabolism via alteration of phosphate homeostasis. In **Figure 10** we show a summary of the results.

**Figure 10.**
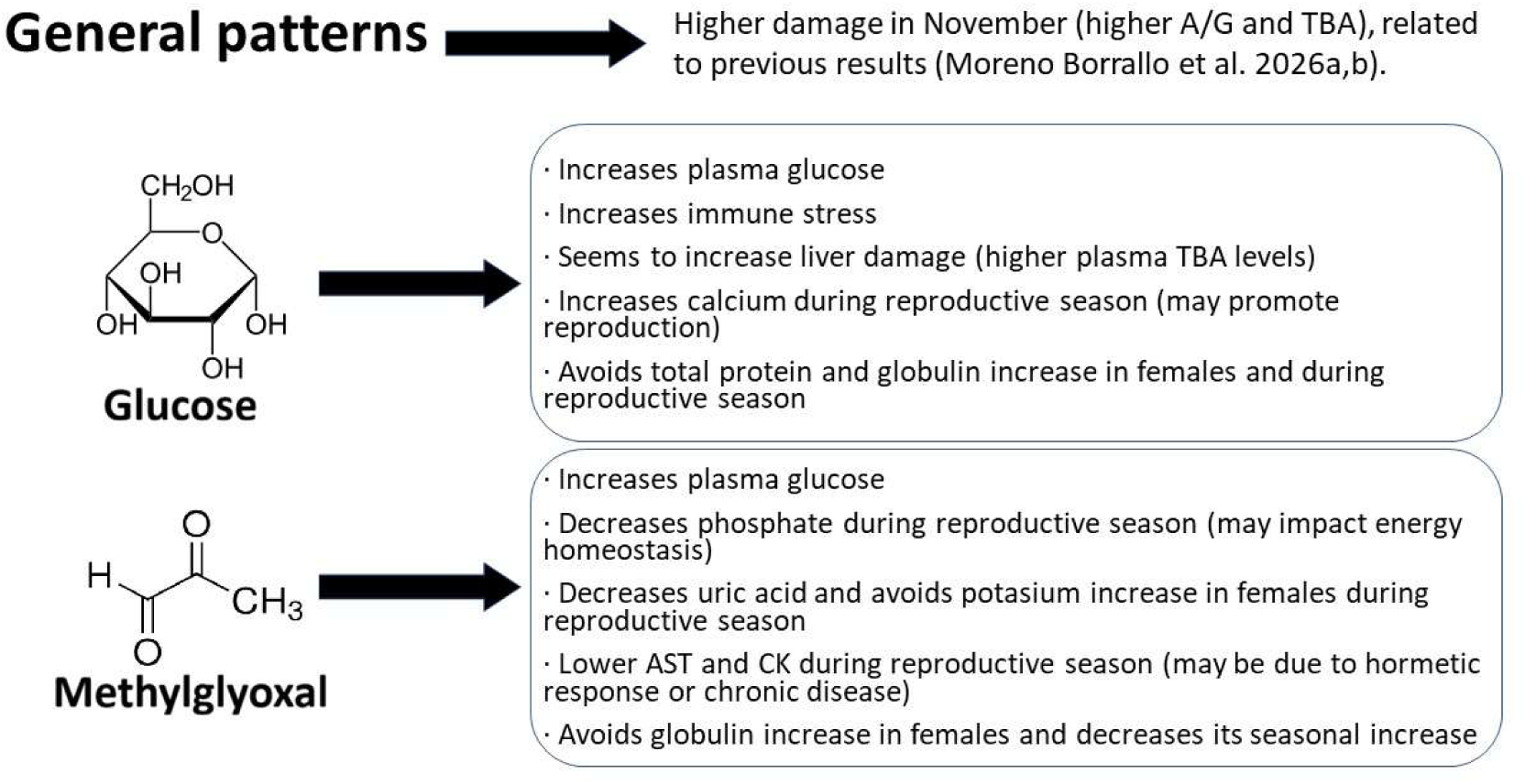
Summary of the experiment results. Results suggest generally higher damage in November, and both treatments increase plasma glucose levels, potentially inducing physiological effects promoting reproductive activation. Glucose supplementation seems to increases hepatic damage and immune stress markers. Effects of methylglyoxal may be more related to protein and phosphate metabolism.

Overall, our findings suggest that some physiological responses of birds to sustained glucose and methylglyoxal exposure may differ from those described in mammals, highlighting the need for further studies to precisely elucidate the underlying mechanisms. The marked dependence of treatment effects on sex and season highlights the complex interplay between nutrition, reproduction, and oxidative balance in avian physiology. Our study provides new insights into the mechanisms that may contribute to avian tolerance to chronically elevated glucose concentrations, while underscoring the need for standardized reference values in avian haematology and further investigation into how birds maintain metabolic homeostasis under glucose levels that would be considered pathological in mammals. We also acknowledge that our study has certain limitations that should be taken into account, including the lack of adequate validation of measurements taken using the veterinary analyzer. Furthermore, the largely exploratory nature of this study involved a large number of statistical comparisons and the inclusion of numerous variables (e.g. sex, age and body mass), which could increase the false discovery rate (FDR). Finally, the current limitations of existing statistical procedures for handling biderictionally anallytically censored data (i.e. left and right censored) restrict the possible inferences (e.g. estimates of mortality hazard). Thus, we call to new developments to facilitate the treatment of similar biochemical data, highlighting especially the limitations that arise from the use of these devices, especially to unwary ecophysiologists, lurked by the apparent ease they may seem to promise.

## Supporting information

ESM1 - Birds list

ESM2 - Extra results

ESM3 - Weather

## Acknowledgments

We would like to thank the Scil Veterinary Company for lending us the Element RC® machine during the experiment, and its representative for their kind assistance with its use, as well as Ascensia Diabetes Care for the donation of glucometer device and strips that were used in our work. We thank Helène Gachot-Neveu, David Bock, Aurélie Hranitzky and Nicolas Spanier for their work in the animal facility. We also thank Max Planck Institute for Biological Intelligence (Pöcking-Seewiesen, Bavaria, Germany) and Bielefeld University (Bielefeld, Germany) for providing part of the birds used in this study.

## Ethics statement

This study adhered to all legal and ethical regulations and was authorized by the French Ministry of Secondary Education and Research (APAFIS #32475-2021071910541808 v5). The work was funded by an ANR grant (AGEs – ANR21-CE02-0009) and the 2020 CNRS – University of Toronto Joint Call for PhD Mobility Funding Programme.

## AI use statement

AI was used for assistance in some code generation, mainly for graphs, and for recommendations on statistical approaches, always under human oversight and judgement. Given the exploratory nature of the study, AI also assisted partly in literature search and exploring possible alternative explanations for results (e.g. brainstorming). This never implied dealing with raw data and human criterium is accountable for all final decisions.

## Data availability statement

The data used for this manuscript is available as Electronic supplementary material. The R code is also available online as **ESM4**.

## Authors contribution statement

A.M.B., F.C. and F.B. obtained funding and contributed to the devise of the experiment and to the writing. A.M.B. led the writing, performed most of the animal experimentation, the analyses with the veterinary analyser, the validation of the glucose measurements and the statistical analyses. F.C. developed and performed the cytometry measurements and A.M.B. and F.C. performed the blood sampling.

