## Supplementary material for "Glucose and methylglyoxal alter plasma protein dynamics, immune traits and glucose homeostasis in a sex- and season-dependent manner in zebra finches": ESM2 - Extra results

#### Traces and posterior distributions of brms models

##### Albumin

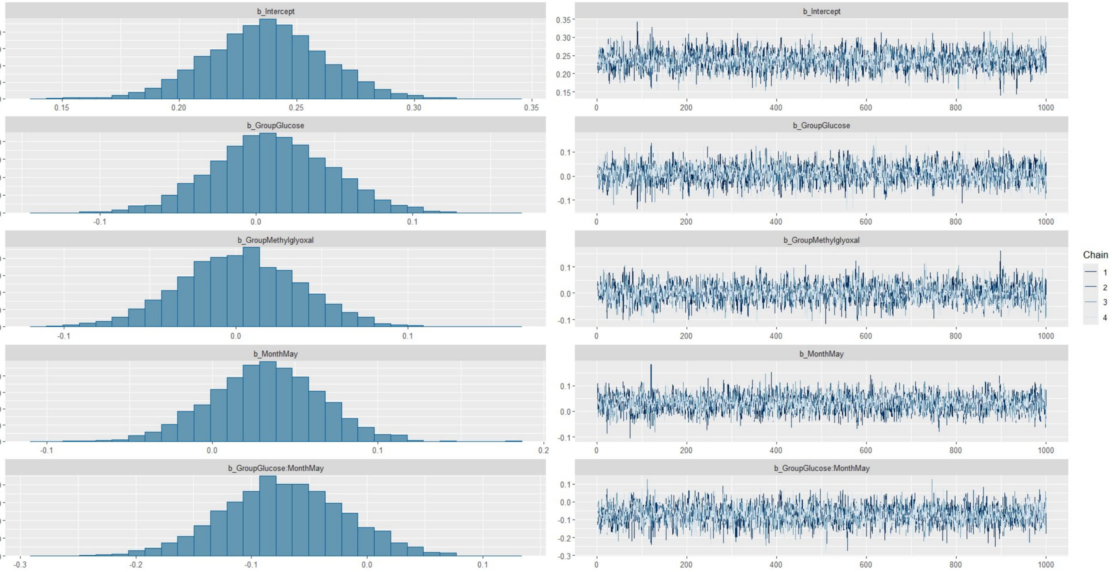


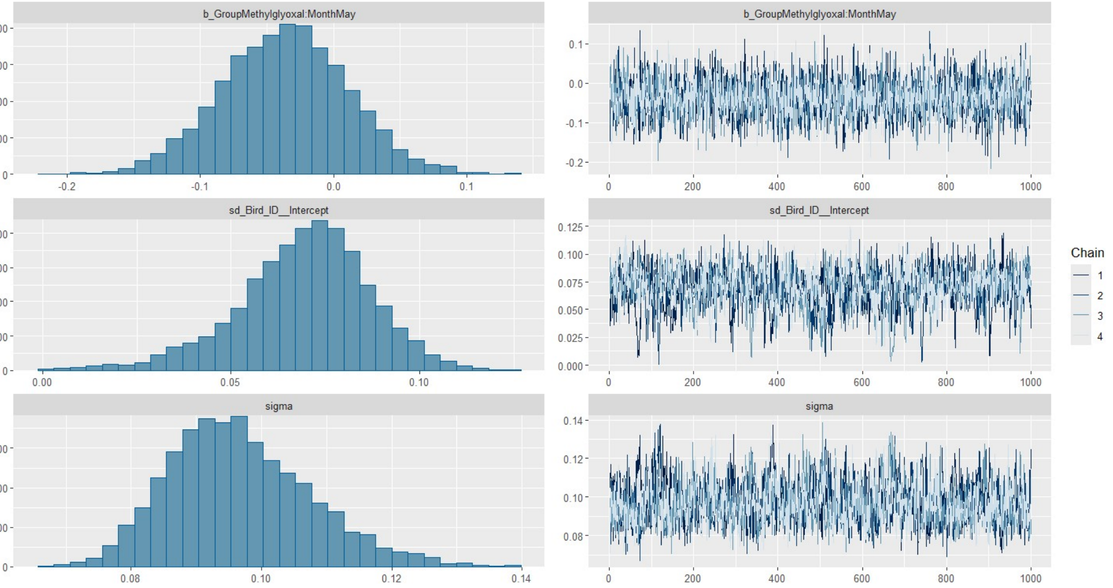

#### **Globulin**

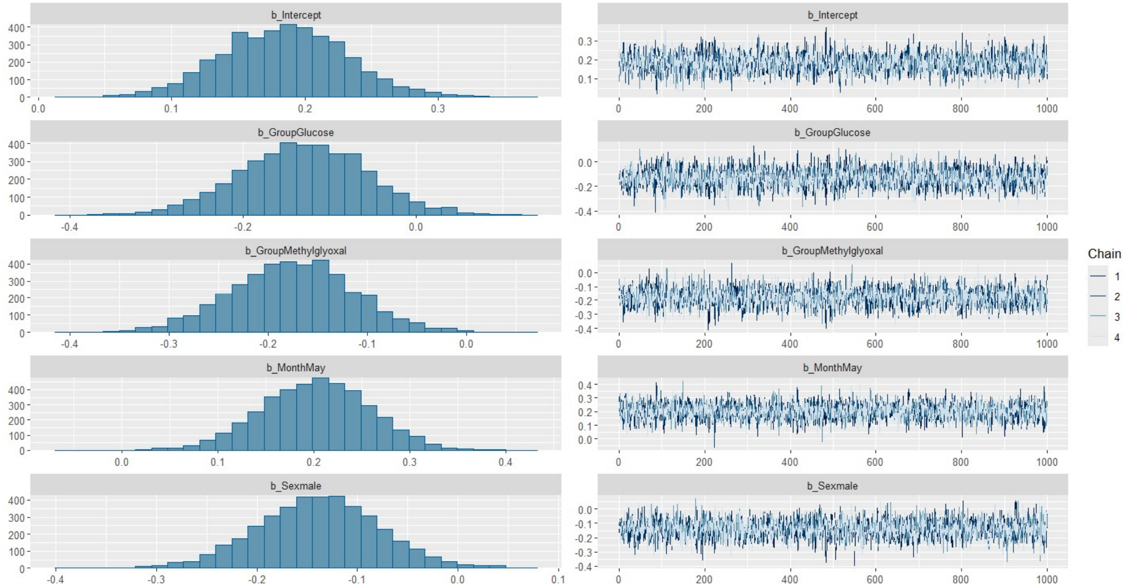
b_Intercept
b_GroupGlucose
b_GroupMethylglyoxal
b_MonthMay
b_Sexmale
Chain
1
2
3
4

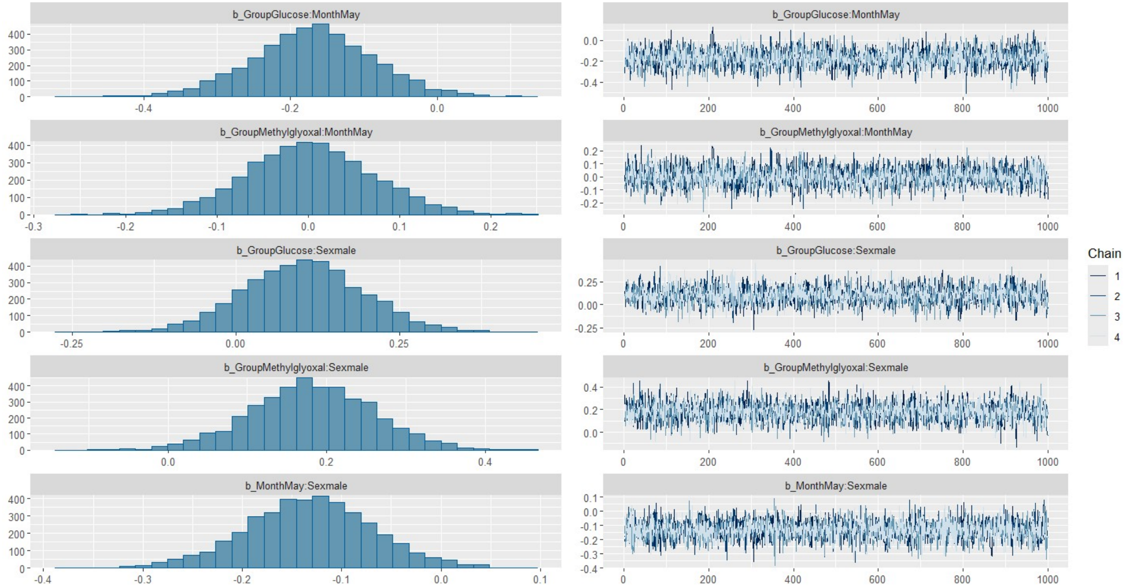
b_GroupGlucose:MonthMay
b_GroupMethylglyoxal:MonthMay
b_GroupGlucose:Sexmale
b_GroupMethylglyoxal:Sexmale
b_MonthMay:Sexmale
Chain
1
2
3
4

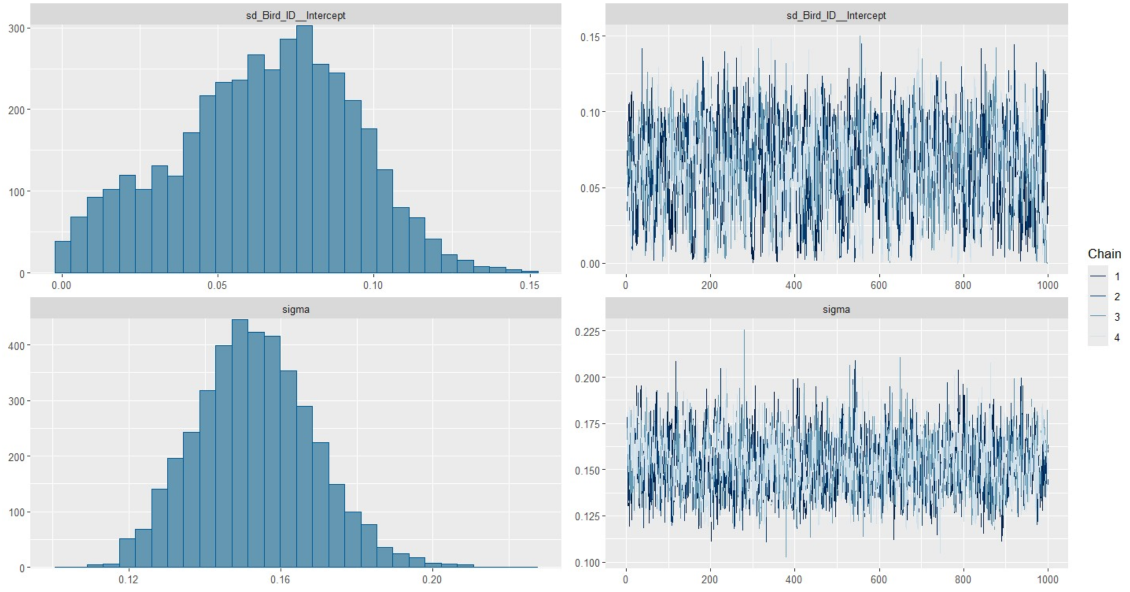
sd_Bird_ID__Intercept
sigma
Chain
1
2
3
4

#### **Total protein**

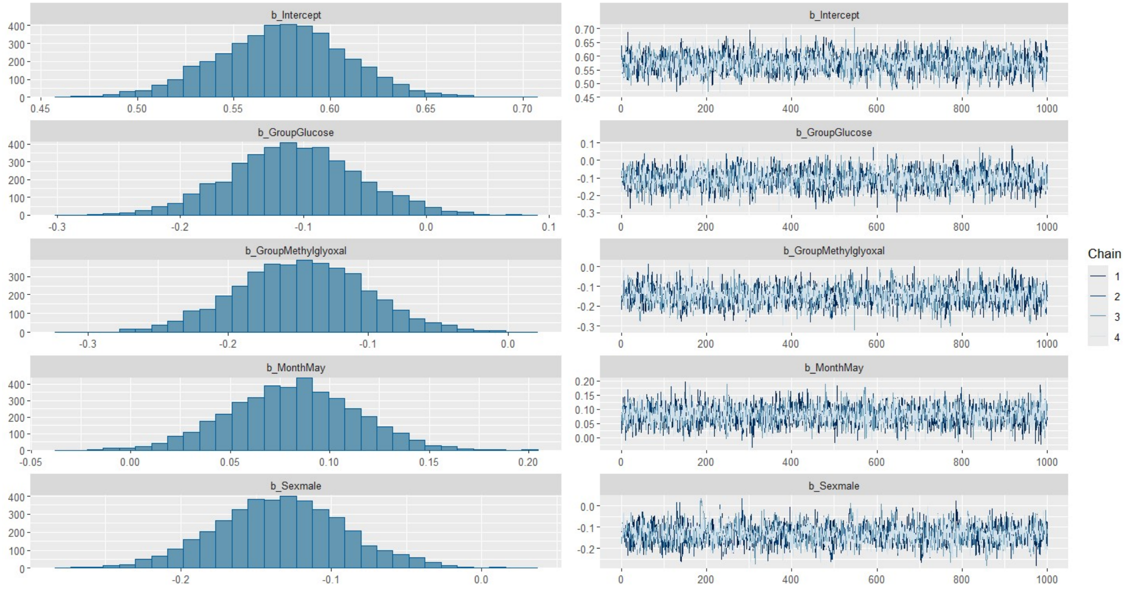


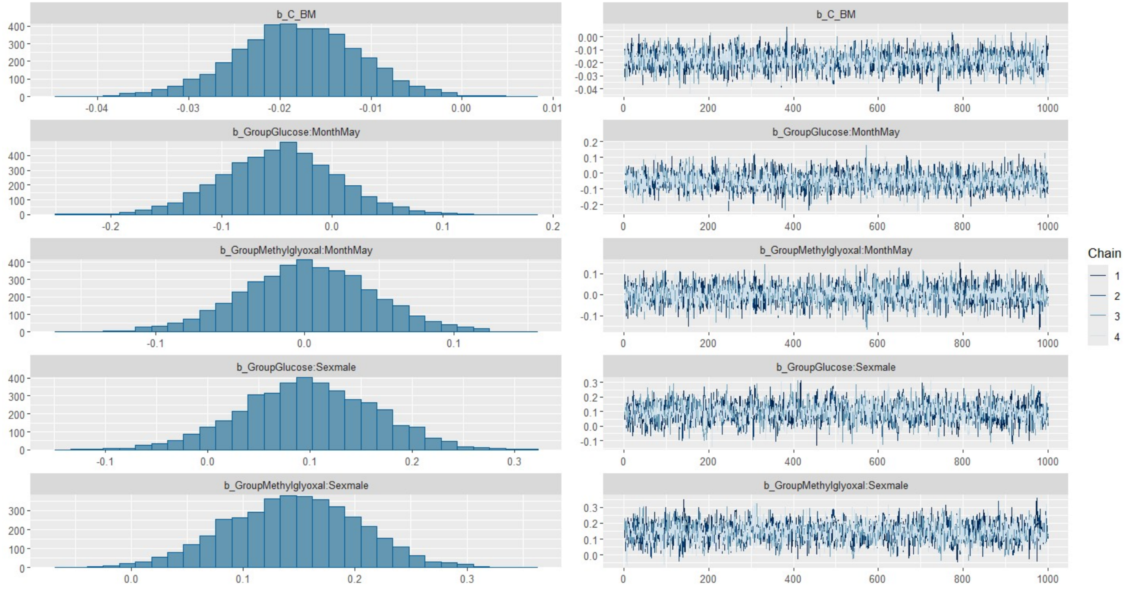


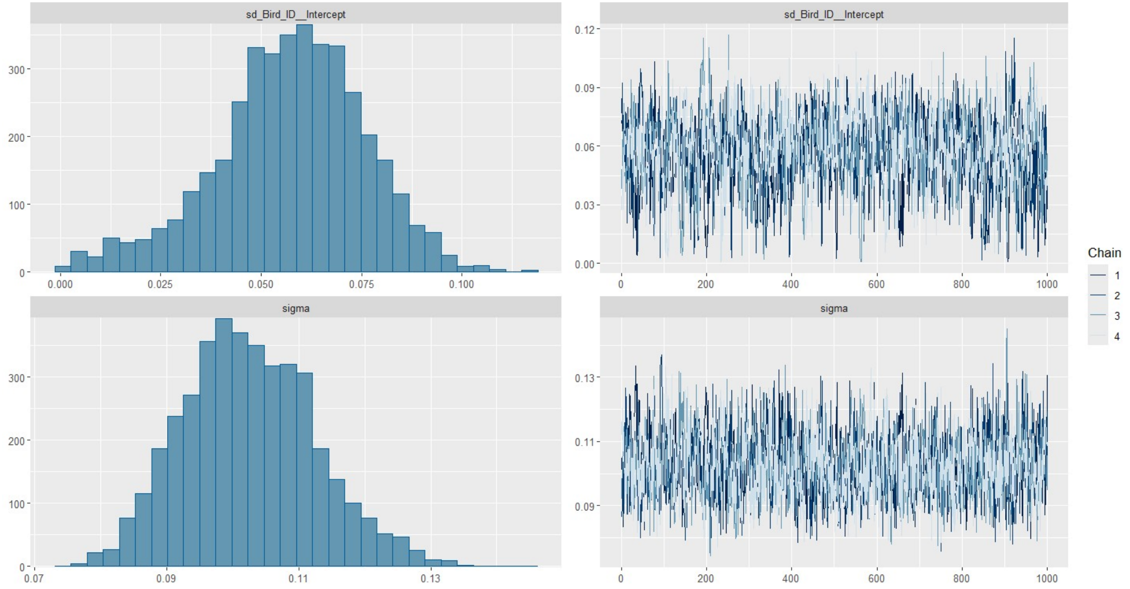

#### A/G ratio

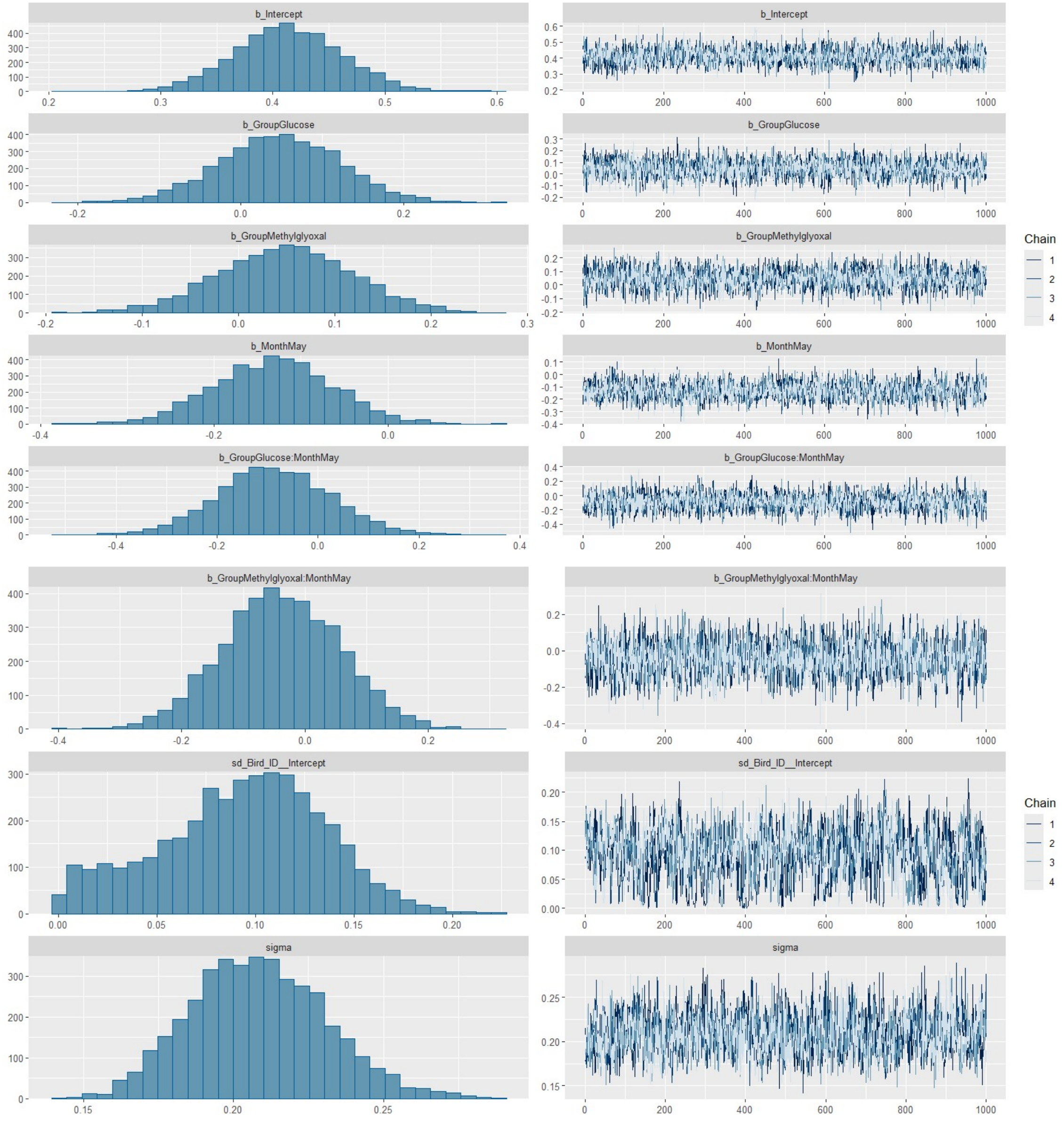
b_Intercept
b_GroupGlucose
b_GroupMethylglyoxal
b_MonthMay
b_GroupGlucose:MonthMay
b_GroupMethylglyoxal:MonthMay
sd_Bird_ID__Intercept
sigma
Chain
1
2
3
4

**AST**

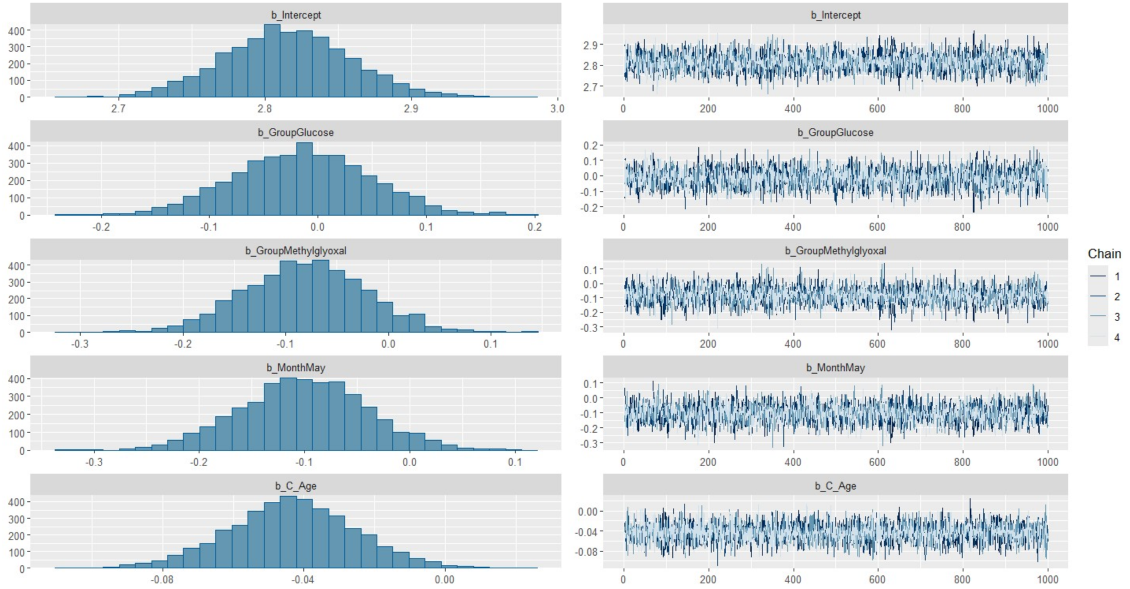
b_Intercept
b_GroupGlucose
b_GroupMethylglyoxal
b_MonthMay
b_C_Age
Chain
1
2
3
4

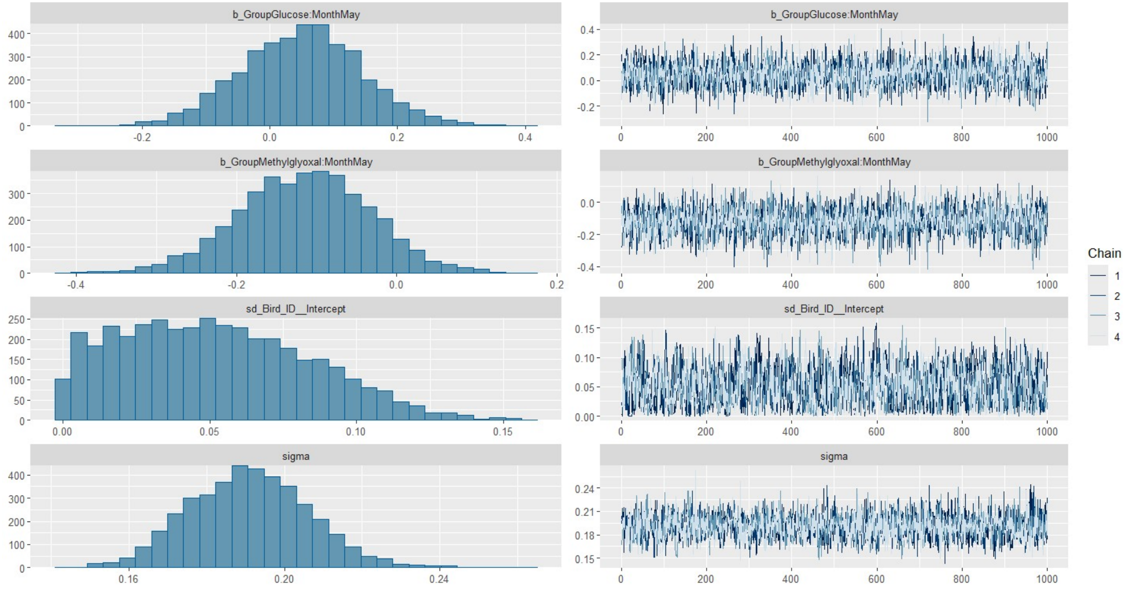
b_GroupGlucose:MonthMay
b_GroupMethylglyoxal:MonthMay
sd_Bird_ID__Intercept
sigma
Chain
1
2
3
4

**CK**

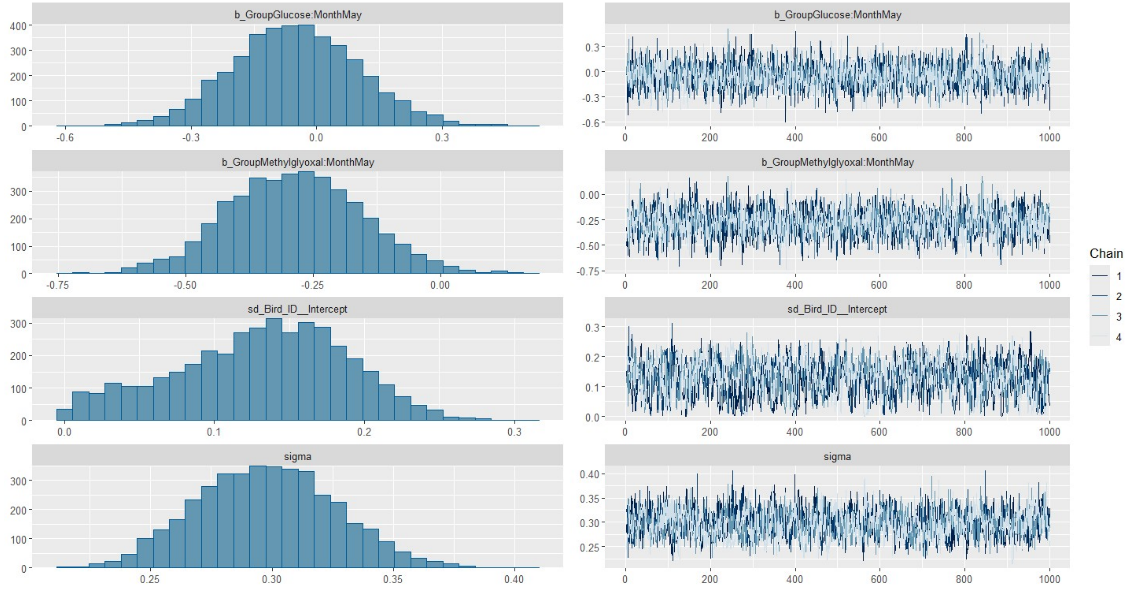
b_GroupGlucose:MonthMay
b_GroupMethylglyoxal:MonthMay
sd_Bird_ID__Intercept
sigma
Chain
1
2
3
4

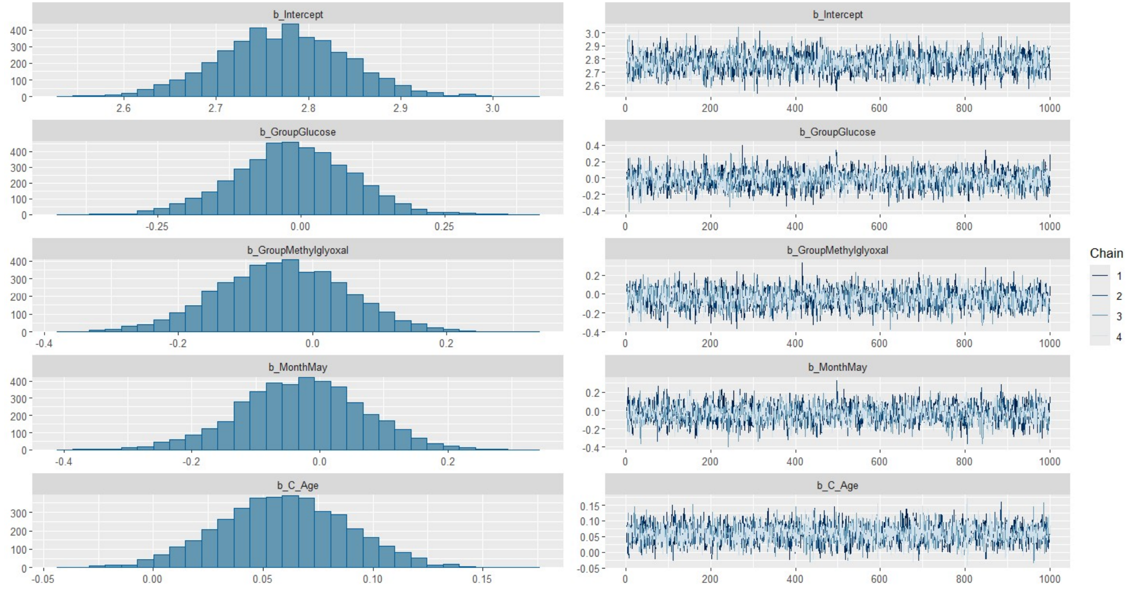
b_Intercept
b_GroupGlucose
b_GroupMethylglyoxal
b_MonthMay
b_C_Age
Chain
1
2
3
4

#### **Glucose**

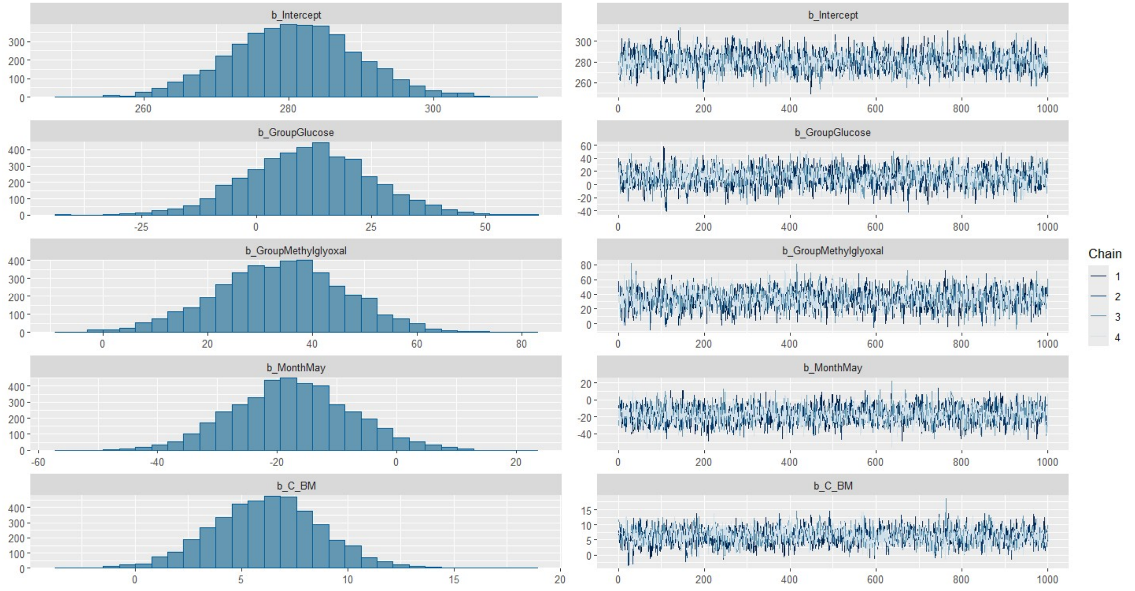

b_Intercept
b_GroupGlucose
b_GroupMethylglyoxal
b_MonthMay
b_C_BM
Chain
1
2
3
4


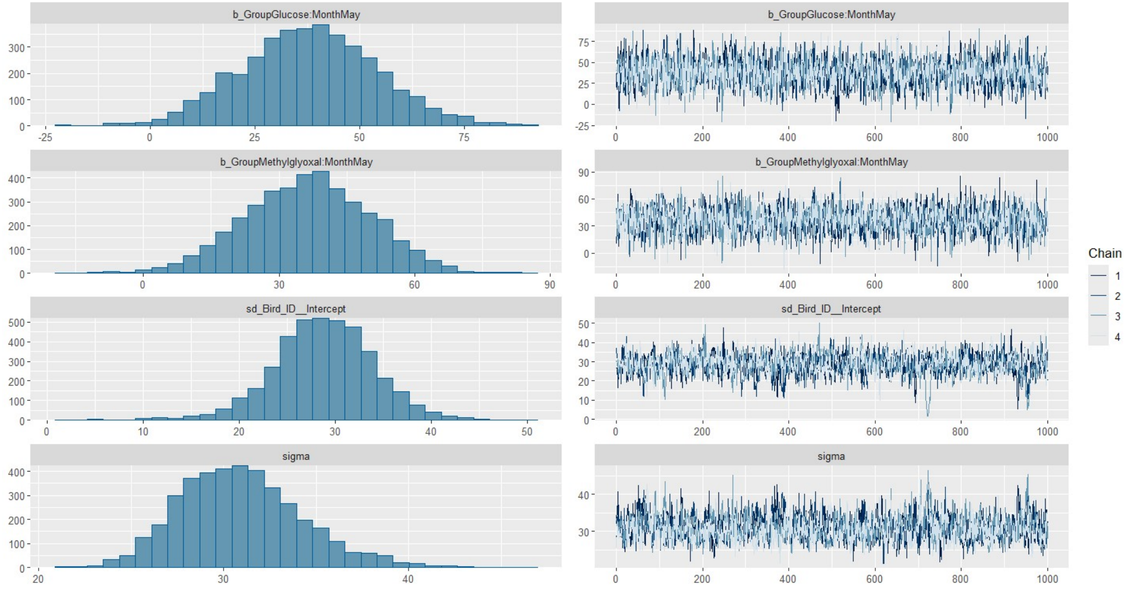

b_GroupGlucose:MonthMay
b_GroupMethylglyoxal:MonthMay
sd_Bird_ID__Intercept
sigma
Chain
1
2
3
4

#### **Total bile acids**

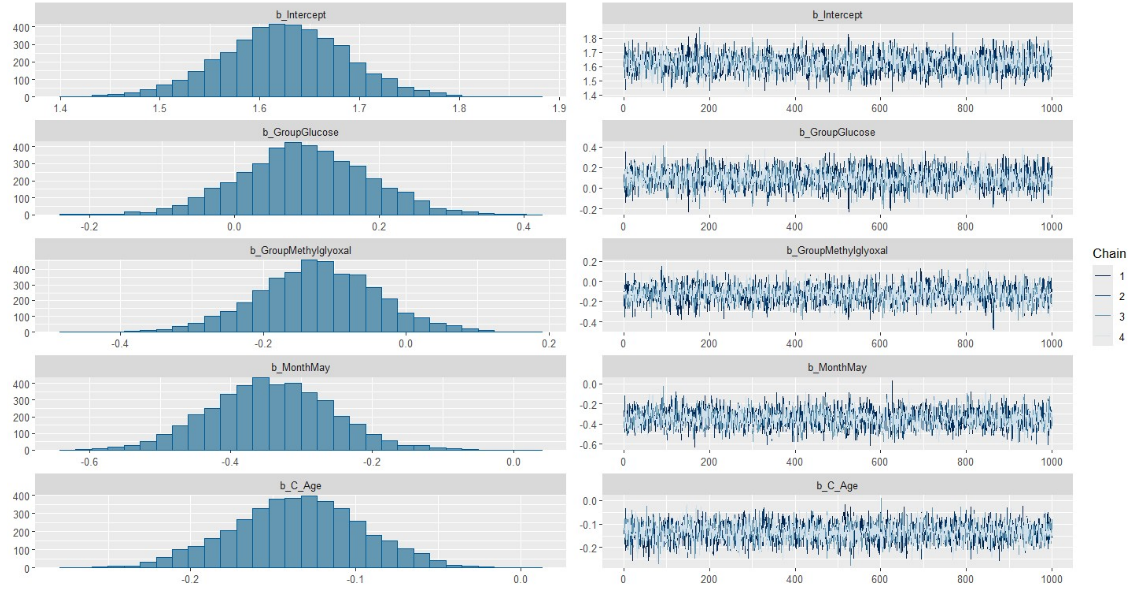


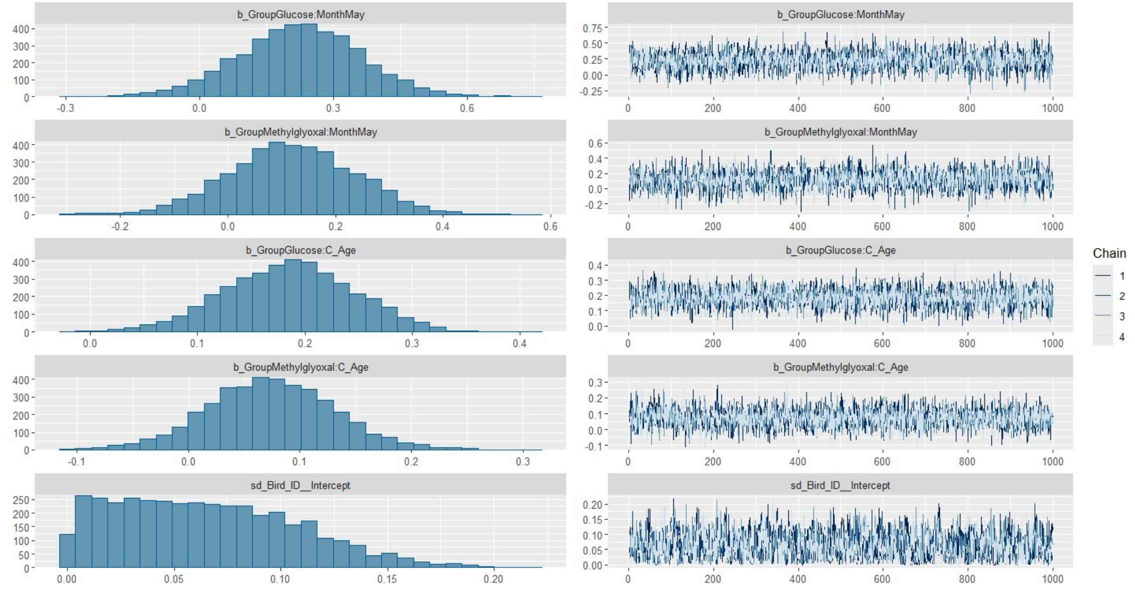


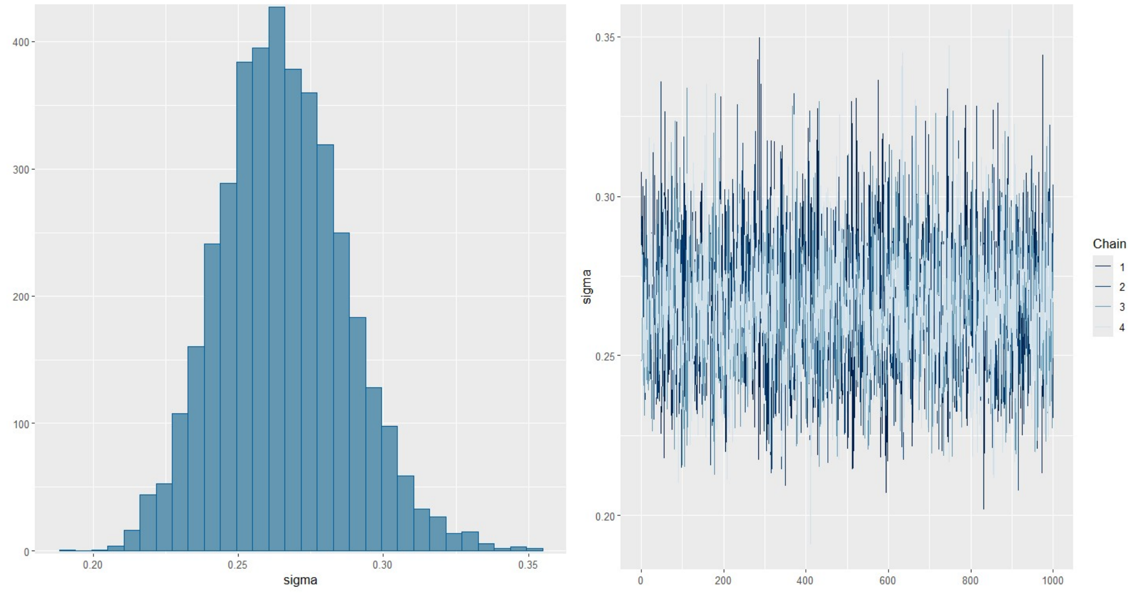

#### Uric acid

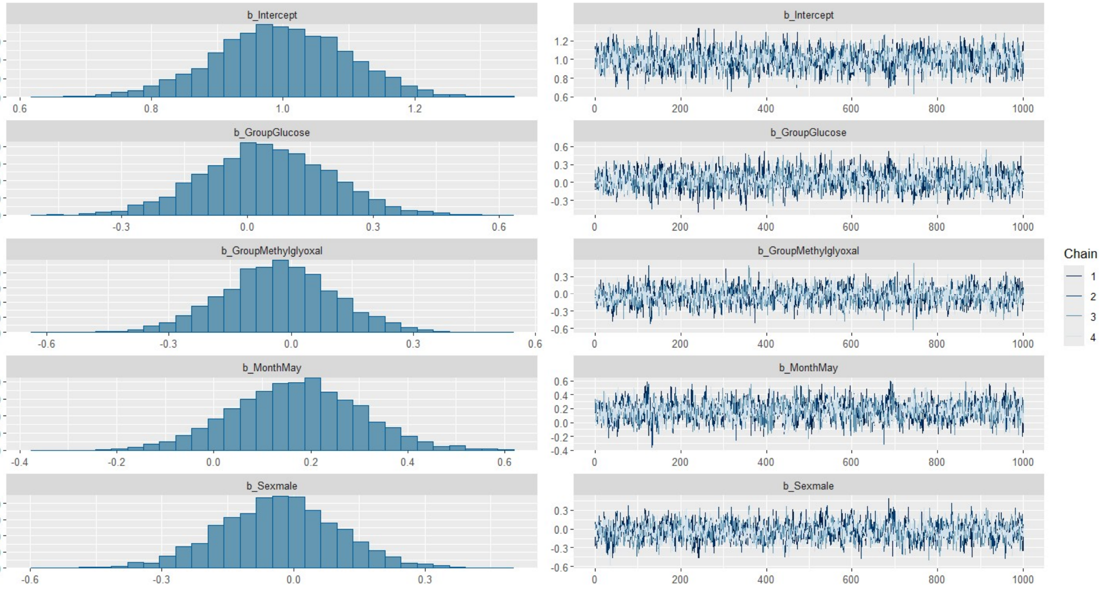
b_Intercept
b_GroupGlucose
b_GroupMethylglyoxal
b_MonthMay
b_Sexmale
Chain
1
2
3
4

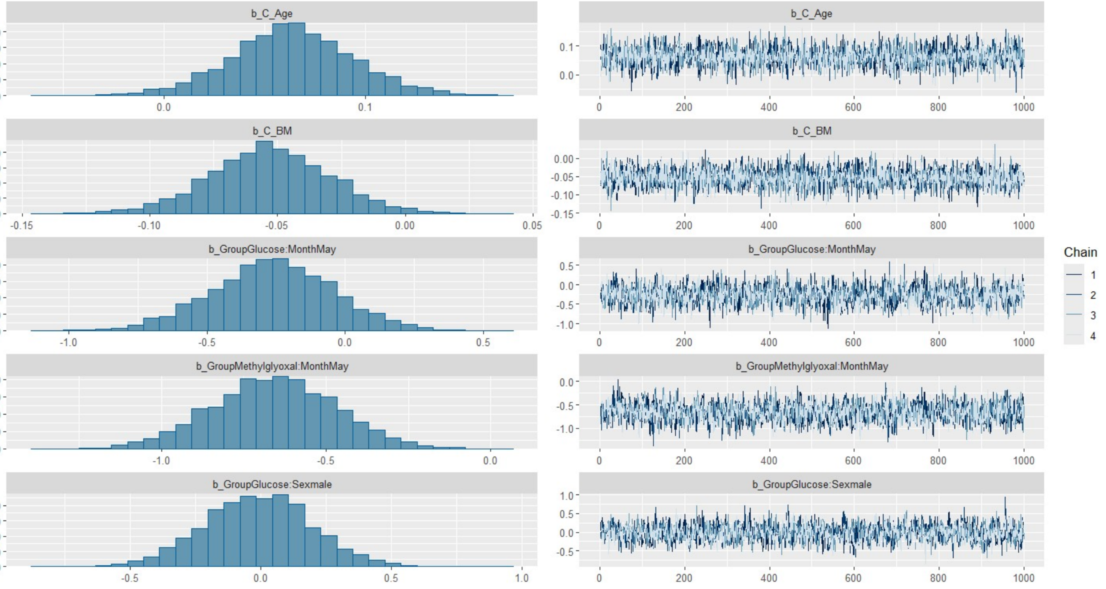
b_C_Age
b_C_BM
b_GroupGlucose:MonthMay
b_GroupMethylglyoxal:MonthMay
b_GroupGlucose:Sexmale
Chain
1
2
3
4

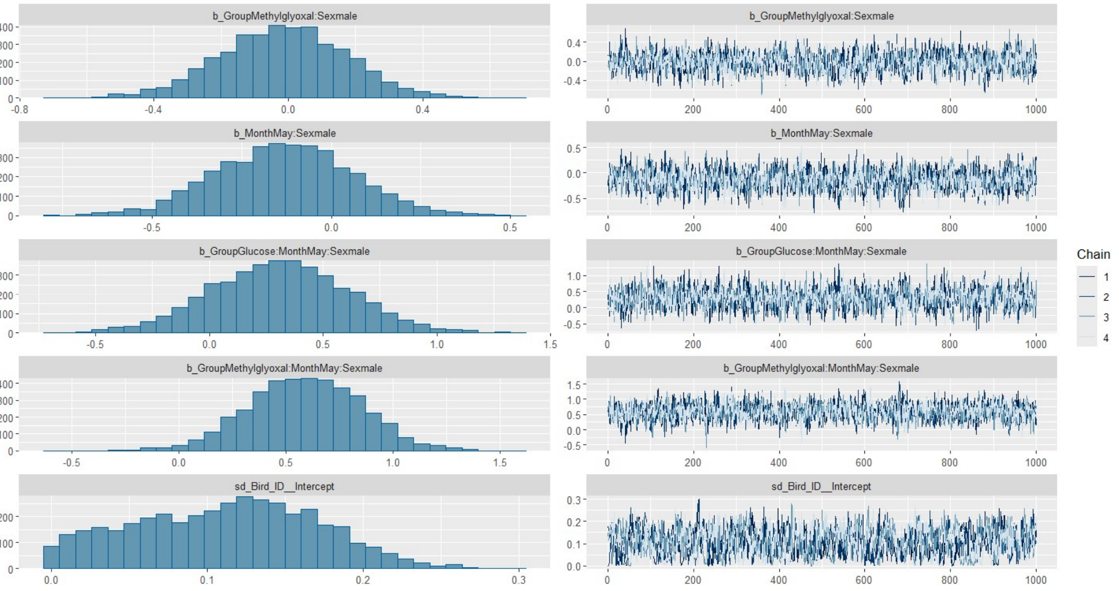
b_GroupMethylglyoxal:Sexmale
b_MonthMay:Sexmale
b_GroupGlucose:MonthMay:Sexmale
b_GroupMethylglyoxal:MonthMay:Sexmale
sd_Bird_ID__Intercept
Chain
1
2
3
4

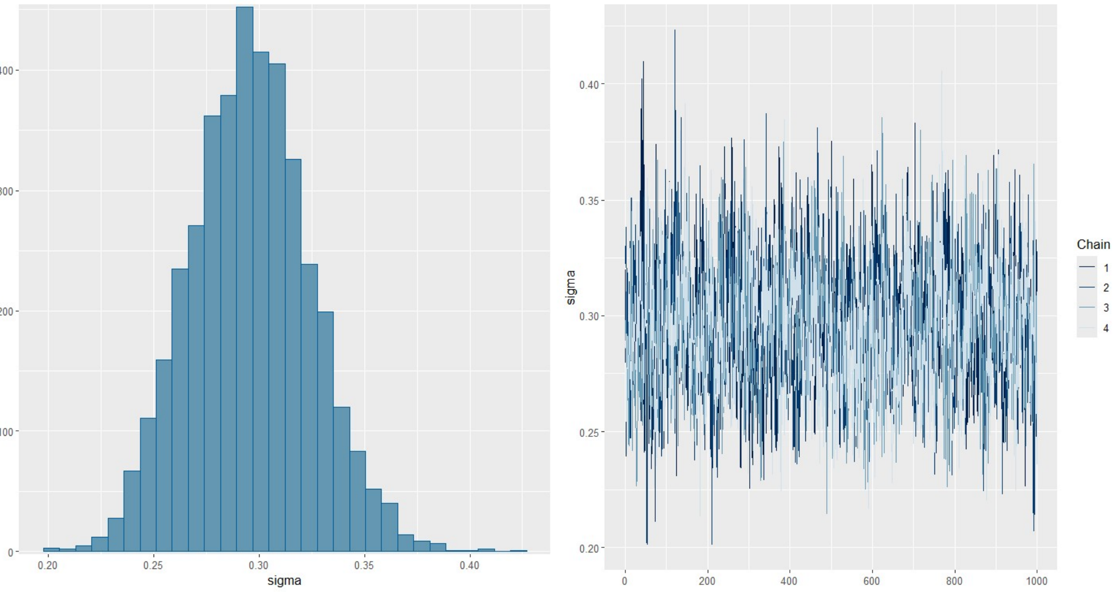
sigma
Chain
1
2
3
4

### Phosphates

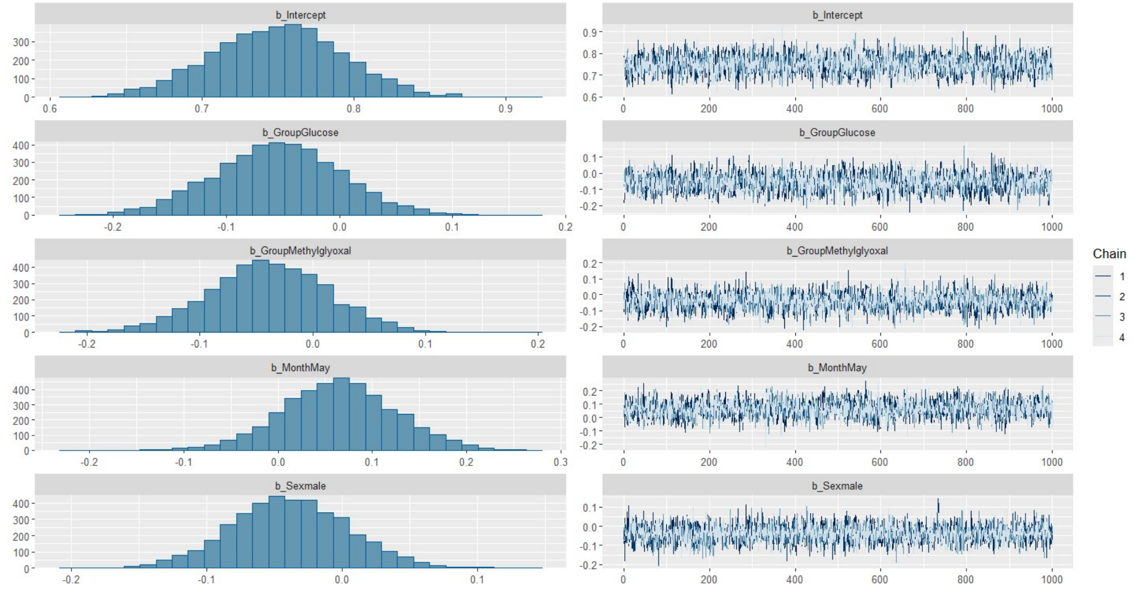
b_Intercept
b_GroupGlucose
b_GroupMethylglyoxal
b_MonthMay
b_Sexmale
Chain
1
2
3
4

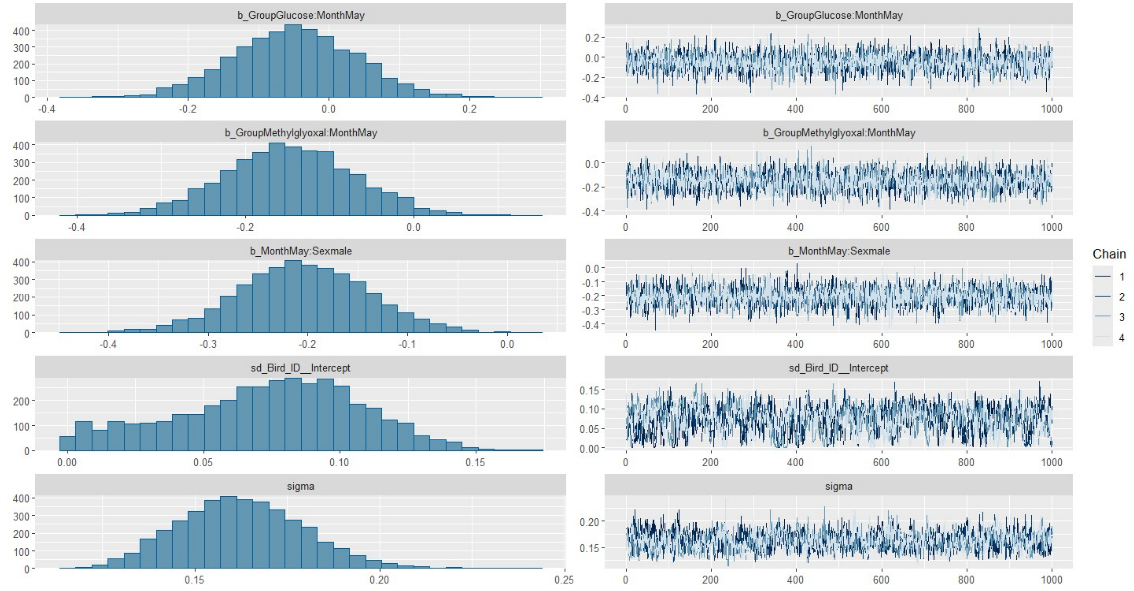
b_GroupGlucose:MonthMay
b_GroupMethylglyoxal:MonthMay
b_MonthMay:Sexmale
sd_Bird_ID__Intercept
sigma
Chain
1
2
3
4

## $Ca^{2+}$

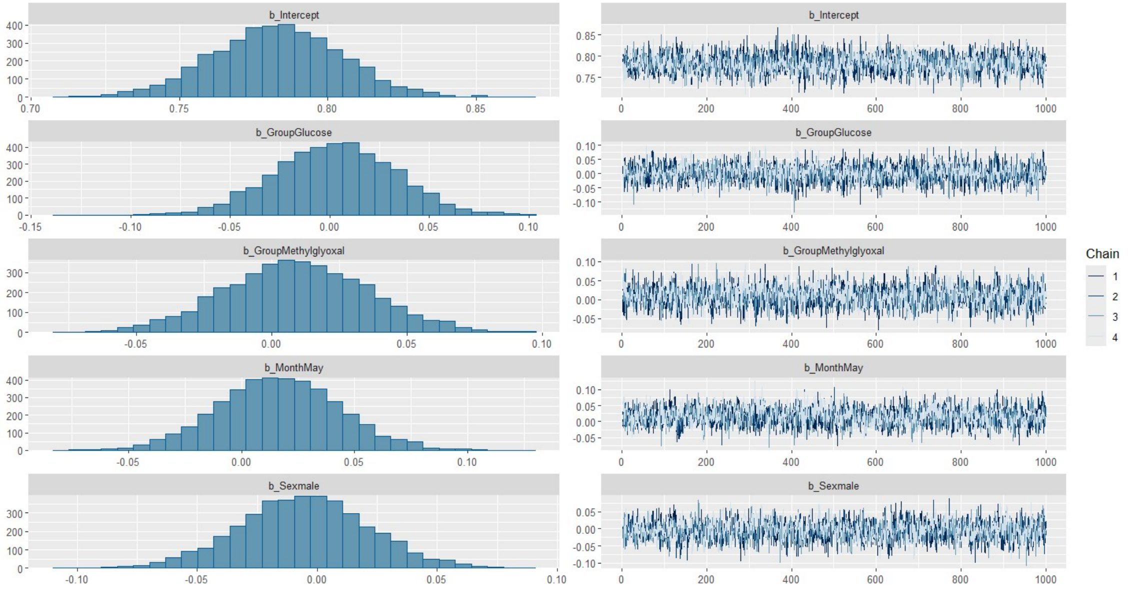
b_Intercept
b_GroupGlucose
b_GroupMethylglyoxal
b_MonthMay
b_Sexmale
Chain
1
2
3
4

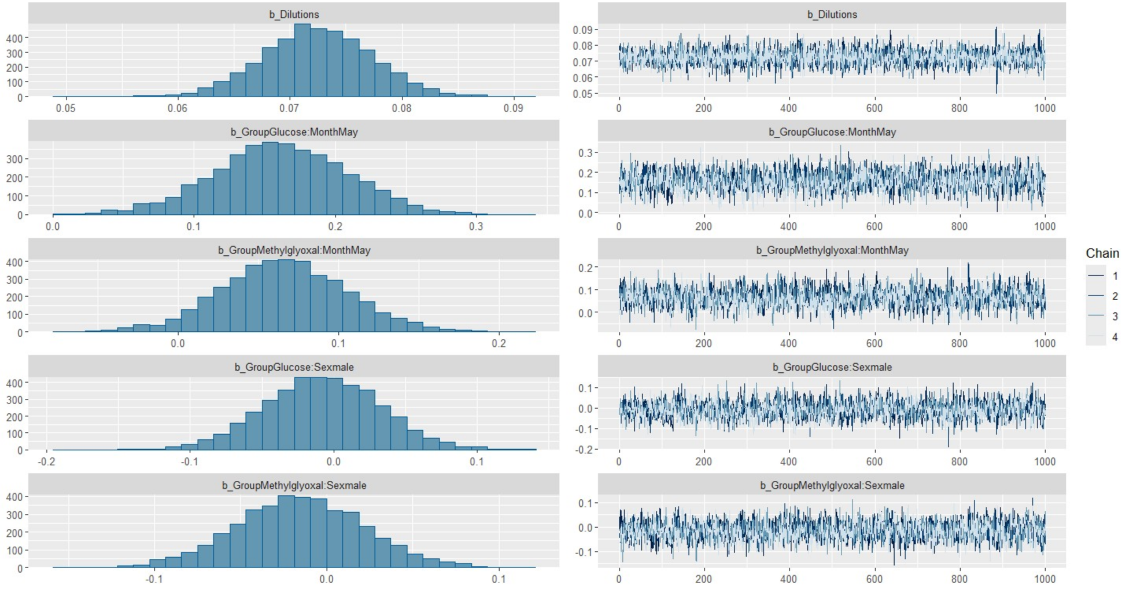
b_Dilutions
b_GroupGlucose:MonthMay
b_GroupMethylglyoxal:MonthMay
b_GroupGlucose:Sexmale
b_GroupMethylglyoxal:Sexmale
Chain
1
2
3
4

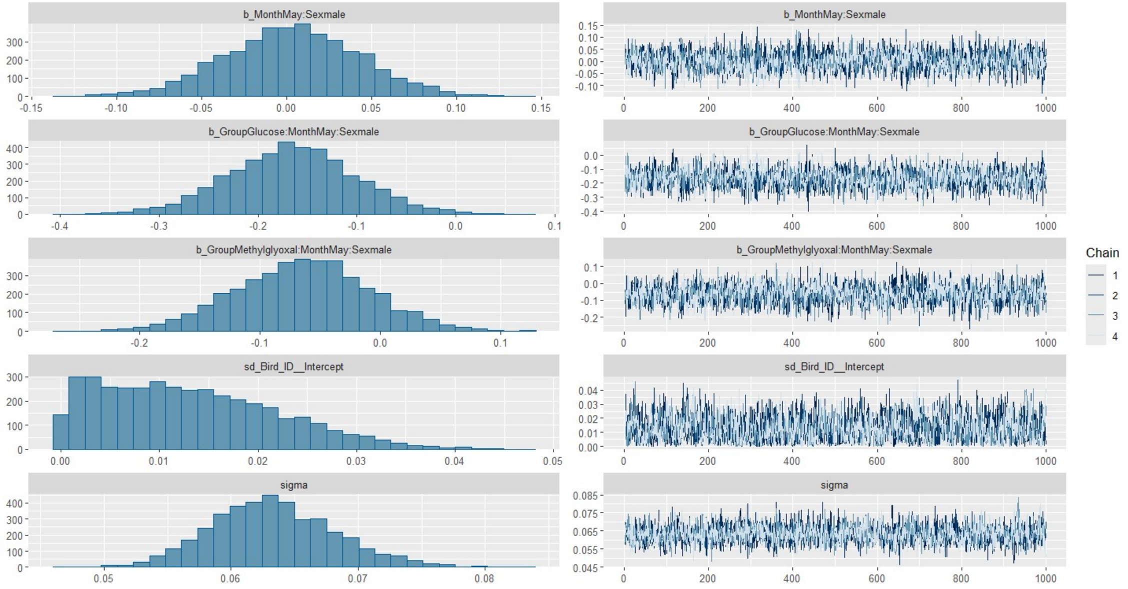
b_MonthMay:Sexmale
b_GroupGlucose:MonthMay:Sexmale
b_GroupMethylglyoxal:MonthMay:Sexmale
sd_Bird_ID__Intercept
sigma
Chain
1
2
3
4

## $K^+$

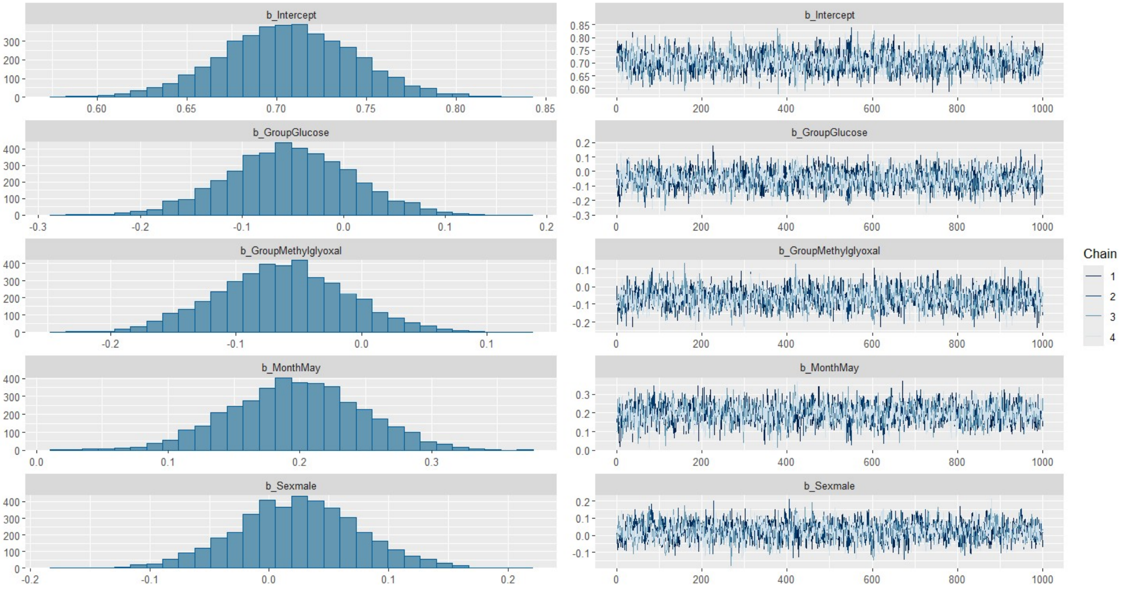
b_Intercept
b_GroupGlucose
b_GroupMethylglyoxal
b_MonthMay
b_Sexmale
Chain
1
2
3
4

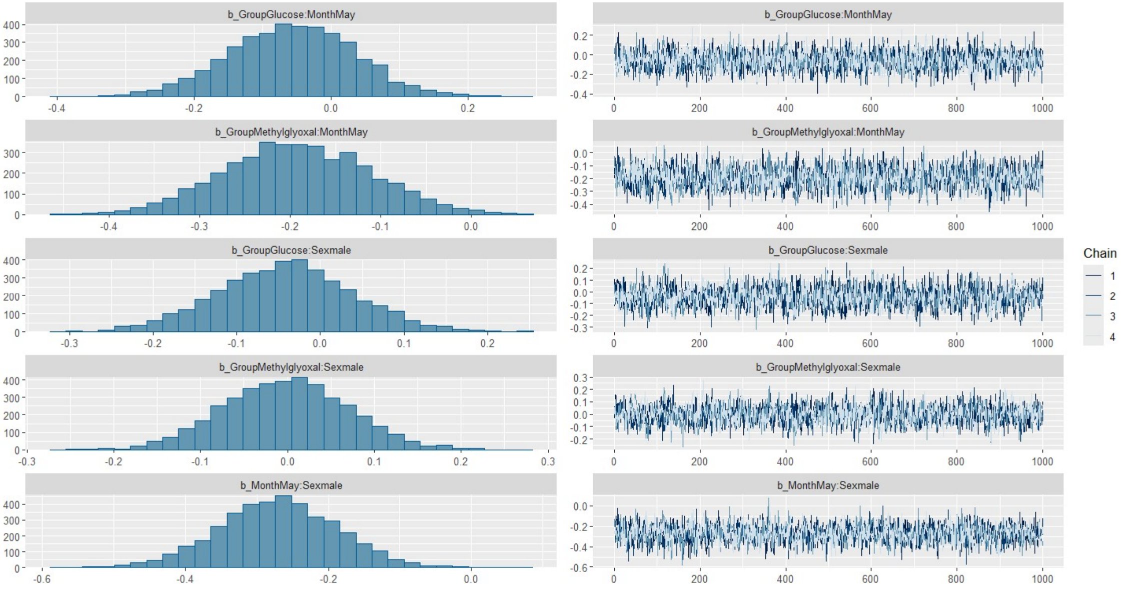
b_GroupGlucose:MonthMay
b_GroupMethylglyoxal:MonthMay
b_GroupGlucose:Sexmale
b_GroupMethylglyoxal:Sexmale
b_MonthMay:Sexmale
Chain
1
2
3
4

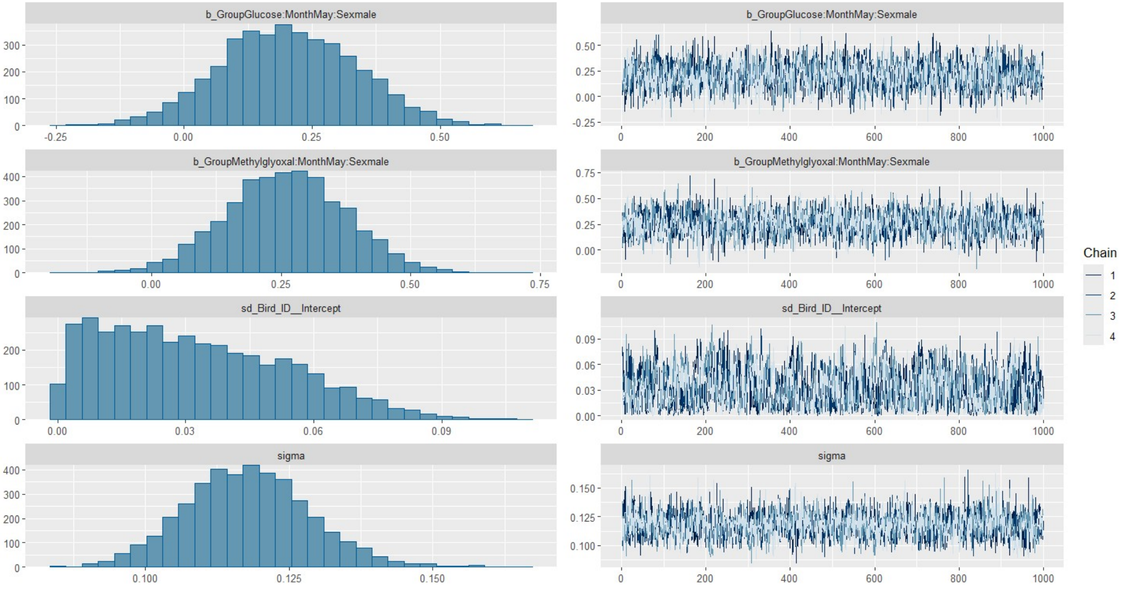
b_GroupGlucose:MonthMay:Sexmale
b_GroupMethylglyoxal:MonthMay:Sexmale
sd_Bird_ID__Intercept
sigma
Chain
1
2
3
4

## <u>$Na^+$</u>

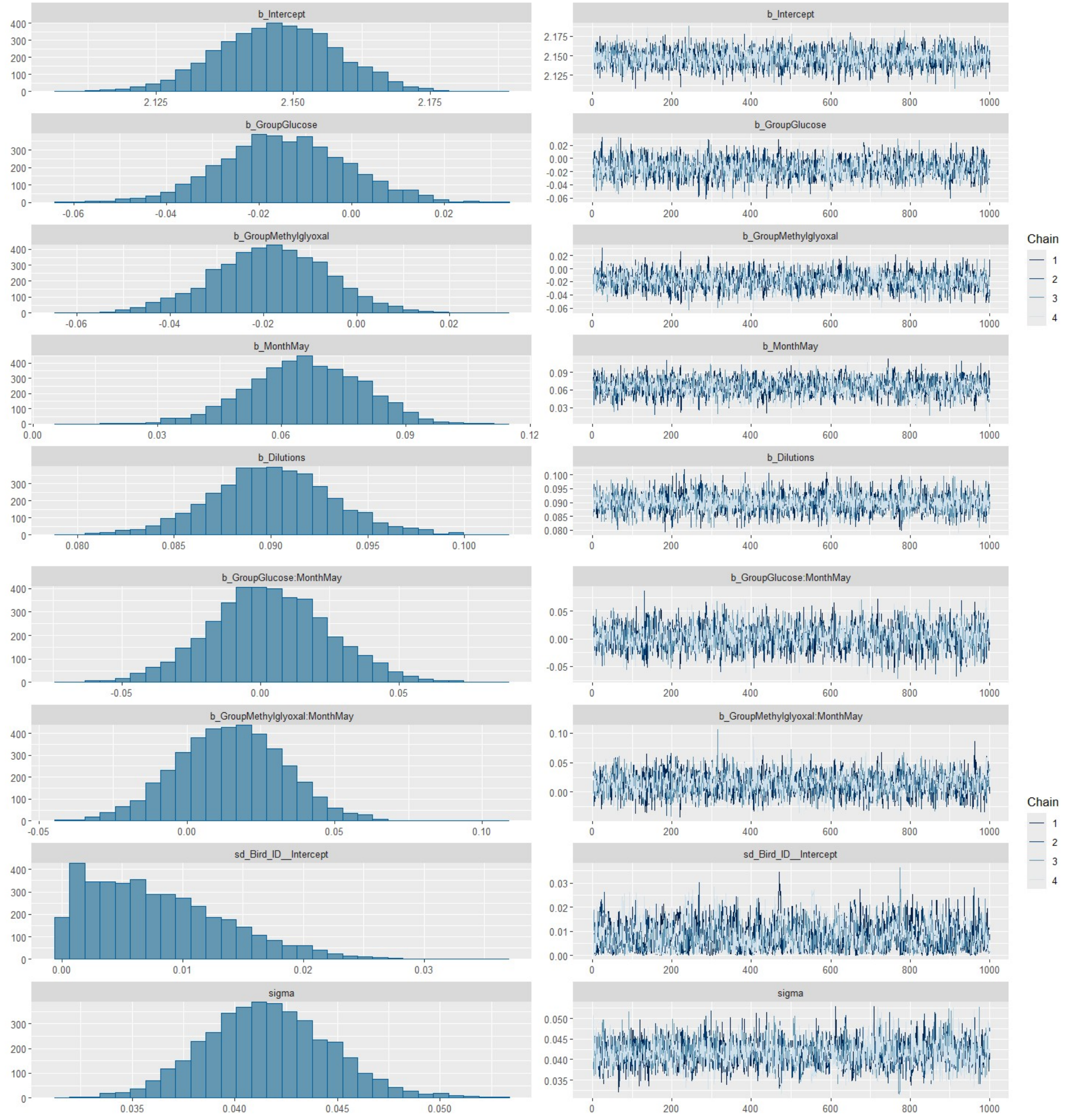


**Figure ESM2.1.** Traces and posterior distributions of the models.

**Table ESM2.1.** Outcome from the model on plasma albumin.

| | Estimate | Est.Error | l-95% CI | u-95% CI | Rhat | Bulk_ESS | Tail_ESS |
|---|---|---|---|---|---|---|---|
| **Intercept** | 0.24 | 0.02 | 0.19 | 0.28 | 1.00 | 2926 | 3130 |
| **Group Glucose** | 0.01 | 0.04 | -0.06 | 0.09 | 1.00 | 2963 | 2740 |
| **Group Methylglyoxal** | 0.00 | 0.04 | -0.07 | 0.07 | 1.00 | 2897 | 3089 |
| **Month May** | 0.03 | 0.03 | -0.03 | 0.10 | 1.00 | 3050 | 2910 |
| **Group Glucose:Month May** | -0.07 | 0.05 | -0.18 | 0.03 | 1.00 | 3465 | 3089 |
| **Group Methylglyoxal:Month May** | -0.04 | 0.05 | -0.13 | 0.06 | 1.00 | 3290 | 3031 |

**Table ESM2.2.** Outcome of loo on the sets of albumin models with the same observations (**A** with age and **B** without age, given the absence of age values for few individuals). ELP differences and their SE are presented. The higher numbers represent simpler models. The simplest model is selected if it is not different from the next most simple one (i.e. the SE of the next one is higher than its ELPD difference).

A

| Model | elpd_diff | se_diff |
|---|---|---|
| **modALB5** | 0.0 | 0.0 |
| **modALB6** | -0.3 | 0.8 |
| **modALB4** | -0.5 | 0.6 |
| **modALB3** | -2.4 | 0.8 |
| **modALB2** | -3.2 | 1.0 |
| **modALB1** | -5.1 | 0.8 |
| **modALB** | -8.1 | 1.6 |

B

| Model | elpd_diff | se_diff |
|---|---|---|
| **modALB9** | 0.0 | 0.0 |
| **modALB10** | 0.0 | 1.3 |
| **modALB8** | -0.2 | 0.8 |
| **modALB7** | -0.9 | 1.3 |

**Table ESM2.3.** Outcome from the model on plasma albumin to globulin ratio (A/G).

| | Estimate | Est.Error | l-95% CI | u-95% CI | Rhat | Bulk_ESS | Tail_ESS |
|---|---|---|---|---|---|---|---|
| **Intercept** | 0.41 | 0.05 | 0.32 | 0.51 | 1.00 | 2450 | 2740 |
| **GroupGlucose** | 0.05 | 0.07 | -0.10 | 0.19 | 1.00 | 2389 | 2493 |
| **GroupMethylglyoxal** | 0.05 | 0.07 | -0.09 | 0.18 | 1.00 | 2612 | 2709 |
| **MonthMay** | -0.13 | 0.07 | -0.26 | 0.00 | 1.00 | 2329 | 2435 |
| **GroupGlucose:MonthMay** | -0.09 | 0.11 | -0.30 | 0.14 | 1.00 | 2742 | 2836 |
| **GroupMethylglyoxal:MonthMay** | -0.04 | 0.09 | -0.22 | 0.15 | 1.00 | 2280 | 2861 |

**Table ESM2.4.** Outcome of loo on the sets of A/G models with the same observations (**A** with age and **B** without age, given the absence of age values for few individuals). ELP differences and their SE are presented. The higher numbers represent simpler models. The simplest model is selected if it is not different from the next most simple one (i.e. the SE of the next one is higher than its ELPD difference).

A

| | elpd_diff | se_diff |
|---|---|---|
| **modA_G5** | 0.0 | 0.0 |
| **modA_G3** | -0.6 | 1.5 |
| **modA_G4** | -0.9 | 0.9 |
| **modA_G2** | -1.4 | 1.6 |
| **modA_G1** | -2.6 | 1.6 |
| **modA_G** | -3.9 | 1.6 |

B

| | elpd_diff | se_diff |
|---|---|---|
| **modA_G8** | 0.0 | 0.0 |
| **modA_G9** | -0.6 | 1.4 |
| **modA_G10** | -0.7 | 1.8 |
| **modA_G7** | -1.9 | 1.0 |
| **modA_G6** | -3.3 | 2.3 |

**Table ESM2.5.** Outcome from the model on plasma globulin.

| | Estimate | Est.Error | l-95% CI | u-95% CI | Rhat | Bulk_ESS | Tail_ESS |
|---|---|---|---|---|---|---|---|
| **Intercept** | 0.18 | 0.05 | 0.09 | 0.28 | 1.00 | 2668 | 3035 |
| **Group Glucose** | -0.13 | 0.07 | -0.27 | 0.01 | 1.00 | 2715 | 2975 |
| **Group Methylglyoxal** | -0.18 | 0.06 | -0.30 | -0.05 | 1.00 | 2951 | 3108 |
| **Month May** | 0.20 | 0.06 | 0.09 | 0.31 | 1.00 | 3517 | 3089 |
| **Sex male** | -0.14 | 0.06 | -0.26 | -0.02 | 1.00 | 2710 | 2814 |
| **Group Glucose:Month May** | -0.17 | 0.08 | -0.34 | -0.01 | 1.00 | 4181 | 3276 |
| **Group Methylglyoxal:Month May** | 0.00 | 0.07 | -0.13 | 0.14 | 1.00 | 3837 | 3168 |
| **Group Glucose:Sex male** | 0.10 | 0.09 | -0.08 | 0.29 | 1.00 | 2823 | 2912 |
| **Group Methylglyoxal:Sex male** | 0.18 | 0.08 | 0.03 | 0.34 | 1.00 | 2934 | 3317 |
| **Month May:Sex male** | -0.13 | 0.07 | -0.26 | -0.00 | 1.00 | 4835 | 2931 |

**Table ESM2.6.** Outcome of loo on the sets of plasma globulin models with the same observations (**A** with age and **B** without age, given the absence of age values for few individuals). ELP differences and their SE are presented. The higher numbers represent simpler models. The simplest model is selected if it is not different from the next most simple one (i.e. the SE of the next one is higher than its ELPD difference).

A

| | elpd_diff | se_diff |
|---|---|---|
| **modGLOB5** | 0.0 | 0.0 |
| **modGLOB4** | -0.7 | 1.1 |
| **modGLOB3** | -1.4 | 1.2 |
| **modGLOB2** | -3.3 | 1.7 |
| **modGLOB1** | -4.2 | 1.7 |
| **modGLOB** | -6.0 | 2.2 |

B

| | elpd_diff | se_diff |
|---|---|---|
| **modGLOB6** | 0.0 | 0.0 |
| **modGLOB7** | -0.5 | 1.6 |

**Table ESM2.7.** Outcome from the model on total plasma proteins.

| | Estimate | Est.Error | l-95% CI | u-95% CI | Rhat | Bulk_ESS | Tail_ESS |
|---|---|---|---|---|---|---|---|
| **Intercept** | 0.58 | 0.03 | 0.51 | 0.64 | 1.00 | 2267 | 2441 |
| **Group Glucose** | -0.11 | 0.05 | -0.21 | -0.00 | 1.00 | 2362 | 2752 |
| **Group Methylglyoxal** | -0.15 | 0.05 | -0.24 | -0.05 | 1.00 | 2217 | 2502 |
| **Month May** | 0.08 | 0.03 | 0.02 | 0.14 | 1.00 | 3441 | 3260 |
| **Sex male** | -0.13 | 0.04 | -0.22 | -0.05 | 1.00 | 2135 | 2051 |
| **C_BM** | -0.02 | 0.01 | -0.03 | -0.00 | 1.00 | 3213 | 2647 |
| **Group Glucose:Month May** | -0.05 | 0.05 | -0.15 | 0.06 | 1.00 | 4390 | 3274 |
| **Group Methylglyoxal:Month May** | 0.00 | 0.04 | -0.09 | 0.09 | 1.00 | 3748 | 3215 |
| **Group Glucose:Sex male** | 0.10 | 0.07 | -0.03 | 0.23 | 1.00 | 2067 | 2459 |
| **Group Methylglyoxal:Sex male** | 0.14 | 0.06 | 0.03 | 0.26 | 1.00 | 2222 | 2431 |

**Table ESM2.8.** Outcome of loo on the sets of total plasma protein models with the same observations (**A** with age and **B** without age, given the absence of age values for few individuals). ELP differences and their SE are presented. The higher numbers represent simpler models. The simplest model is selected if it is not different from the next most simple one (i.e. the SE of the next one is higher than its ELPD difference).

A

| | elpd_diff | se_diff |
|---|---|---|
| **modTP5** | 0.0 | 0.0 |
| **modTP3** | -0.7 | 1.0 |
| **modTP4** | -0.8 | 0.9 |
| **modTP2** | -2.3 | 1.3 |
| **modTP1** | -3.6 | 1.5 |
| **modTP** | -4.4 | 2.2 |

**B**

| | elpd_diff | se_diff |
|---|---|---|
| **modTP6** | 0.0 | 0.0 |
| **modTP7** | -0.4 | 1.7 |

**Table ESM2.9.** Outcome from the model on plasma AST.

| | Estimate | Est.Error | l-95% CI | u-95% CI | Rhat | Bulk_ESS | Tail_ESS |
|---|---|---|---|---|---|---|---|
| **Intercept** | 2.82 | 0.04 | 2.73 | 2.90 | 1.00 | 3492 | 2951 |
| **Group Glucose** | -0.01 | 0.06 | -0.14 | 0.10 | 1.00 | 3555 | 2842 |
| **Group Methylglyoxal** | -0.08 | 0.06 | -0.20 | 0.03 | 1.00 | 3724 | 3136 |
| **Month May** | -0.10 | 0.06 | -0.22 | 0.02 | 1.00 | 3113 | 2853 |
| **C_Age** | -0.04 | 0.02 | -0.08 | -0.01 | 1.00 | 6333 | 2821 |
| **Group Glucose:Month May** | 0.05 | 0.10 | -0.14 | 0.24 | 1.00 | 3449 | 3140 |
| **Group Methylglyoxal:Month May** | -0.12 | 0.08 | -0.28 | 0.04 | 1.00 | 3439 | 2768 |

**Table ESM2.10.** Outcome of loo on the sets of plasma AST models with the same observations (**A** with age and **B** without age, given the absence of age values for few individuals). ELP differences and their SE are presented. The higher numbers represent simpler models. The simplest model is selected if it is not different from the next most simple one (i.e. the SE of the next one is higher than its ELPD difference).

| | elpd_diff | se_diff |
|---|---|---|
| **modAST9** | 0.0 | 0.0 |
| **modAST8** | -0.2 | 1.6 |
| **modAST7** | -0.8 | 2.4 |
| **modAST6** | -2.1 | 2.5 |
| **modAST5** | -2.7 | 2.9 |
| **modAST4** | -2.7 | 2.9 |
| **modAST3** | -4.8 | 2.9 |
| **modAST2** | -5.4 | 3.0 |
| **modAST1** | -7.2 | 2.9 |
| **modAST** | -8.4 | 3.2 |

**Table ESM2.11.** Outcome from the model on plasma CK.

| | Estimate | Est.Error | l-95% CI | u-95% CI | Rhat | Bulk_ESS | Tail_ESS |
|---|---|---|---|---|---|---|---|
| **Intercept** | 2.77 | 0.07 | 2.64 | 2.91 | 1.00 | 2656 | 3147 |
| **Group Glucose** | -0.01 | 0.10 | -0.22 | 0.18 | 1.00 | 2932 | 3010 |
| **Group Methylglyoxal** | -0.05 | 0.10 | -0.24 | 0.14 | 1.00 | 2595 | 3171 |
| **Month May** | -0.03 | 0.10 | -0.23 | 0.15 | 1.00 | 2511 | 2739 |
| **C_Age** | 0.06 | 0.03 | 0.00 | 0.12 | 1.00 | 3264 | 2933 |
| **Group Glucose:Month May** | -0.05 | 0.15 | -0.34 | 0.25 | 1.00 | 2677 | 3180 |
| **Group Methylglyoxal:Month May** | -0.29 | 0.13 | -0.54 | -0.02 | 1.00 | 2321 | 2576 |

**Table ESM2.12.** Outcome of loo on the sets of plasma CK models with the same observations. ELP differences and their SE are presented. The higher numbers represent simpler models. The simplest model is selected if it is not different from the next most simple one (i.e. the SE of the next one is higher than its ELPD difference).

| | elpd_diff | se_diff |
|---|---|---|
| **modCK8** | 0.0 | 0.0 |
| **modCK9** | 0.0 | 1.2 |
| **modCK7** | -0.3 | 1.4 |
| **modCK6** | -0.9 | 2.2 |
| **modCK5** | -1.8 | 2.5 |
| **modCK4** | -2.5 | 2.6 |
| **modCK3** | -3.5 | 2.7 |
| **modCK2** | -5.0 | 2.8 |
| **modCK1** | -5.5 | 3.8 |
| **modCK** | -7.1 | 4.1 |

**Table ESM2.13.** Outcome from the model on plasma glucose.

| | Estimate | Est.Error | l-95% CI | u-95% CI | Rhat | Bulk_ESS | Tail_ESS |
|---|---|---|---|---|---|---|---|
| **Intercept** | 280.77 | 8.86 | 263.49 | 298.38 | 1.00 | 1633 | 2124 |
| **Group Glucose** | 11.40 | 13.57 | -15.13 | 38.46 | 1.00 | 1885 | 2717 |
| **Group Methylglyoxal** | 33.95 | 12.51 | 8.63 | 58.01 | 1.00 | 1922 | 2807 |
| **Month May** | -17.40 | 9.98 | -36.81 | 2.66 | 1.00 | 2523 | 2609 |
| **C_BM** | 6.20 | 2.55 | 1.22 | 11.27 | 1.00 | 2256 | 2527 |
| **Group Glucose:Month May** | 38.04 | 16.26 | 7.15 | 70.36 | 1.00 | 2840 | 2874 |
| **Group Methylglyoxal:Month May** | 36.33 | 13.98 | 8.90 | 62.65 | 1.00 | 2641 | 3001 |

**Table ESM2.14.** Outcome of loo on the sets of plasma glucose models with the same observations. ELP differences and their SE are presented. The higher numbers represent simpler models. The simplest model is selected if it is not different from the next most simple one (i.e. the SE of the next one is higher than its ELPD difference).

| | elpd_diff | se_diff |
|---|---|---|
| **modGLU8** | 0.0 | 0.0 |
| **modGLU7** | 0.0 | 0.7 |
| **modGLU6** | -0.3 | 1.6 |
| **modGLU3** | -0.6 | 2.4 |
| **modGLU5** | -0.9 | 2.4 |
| **modGLU4** | -1.4 | 2.2 |
| **modGLU2** | -3.1 | 2.6 |
| **modGLU1** | -4.1 | 2.6 |
| **modGLU** | -7.4 | 2.8 |

**Table ESM2.15.** Outcome from the model on uric acid.

| | Estimate | Est.Error | l-95% CI | u-95% CI | Rhat | Bulk_ESS | Tail_ESS |
|---|---|---|---|---|---|---|---|
| **Intercept** | 1.00 | 0.10 | 0.80 | 1.19 | 1.00 | 1774 | 2466 |
| **Group Glucose** | 0.05 | 0.15 | -0.25 | 0.34 | 1.00 | 1990 | 2810 |
| **Group Methylglyoxal** | -0.03 | 0.14 | -0.31 | 0.24 | 1.00 | 1758 | 2975 |
| **Month May** | 0.17 | 0.13 | -0.09 | 0.43 | 1.00 | 1667 | 2342 |
| **Sex male** | -0.03 | 0.14 | -0.30 | 0.24 | 1.00 | 1485 | 2242 |
| **C_Age** | 0.06 | 0.03 | 0.00 | 0.13 | 1.00 | 4235 | 3182 |
| **C_BM** | -0.05 | 0.02 | -0.10 | -0.01 | 1.00 | 4363 | 2766 |
| **Group Glucose:Month May** | -0.26 | 0.22 | -0.71 | 0.18 | 1.00 | 2038 | 2748 |
| **Group Methylglyoxal:Month May** | -0.66 | 0.19 | -1.03 | -0.29 | 1.00 | 1708 | 2319 |
| **Group Glucose:Sex male** | -0.01 | 0.21 | -0.41 | 0.39 | 1.00 | 1701 | 2671 |
| **Group Methylglyoxal:Sex male** | -0.02 | 0.19 | -0.40 | 0.36 | 1.00 | 1542 | 2473 |
| **Month May:Sex male** | -0.13 | 0.19 | -0.51 | 0.24 | 1.00 | 1533 | 2366 |
| **Group Glucose:Month May:Sex male** | 0.31 | 0.30 | -0.30 | 0.90 | 1.00 | 1840 | 2450 |
| **Group Methylglyoxal:Month May:Sex male** | 0.59 | 0.26 | 0.08 | 1.09 | 1.00 | 1556 | 2426 |

**Table ESM2.16.** Outcome of loo on the sets of plasma uric acid models with the same observations. ELP differences and their SE are presented. The higher numbers represent simpler models. The simplest model is selected if it is not different from the next most simple one (i.e. the SE of the next one is higher than its ELPD difference).

| | elpd_diff | se_diff |
|---|---|---|
| **modUA2** | 0.0 | 0.0 |
| **modUA4** | -0.5 | 2.4 |
| **modUA3** | -0.7 | 1.9 |
| **modUA1** | -0.9 | 0.6 |
| **modUA** | -1.7 | 0.7 |

**Table ESM2.17.** Outcome from the model on total bile acid.

| | Estimate | Est.Error | l-95% CI | u-95% CI | Rhat | Bulk_ESS | Tail_ESS |
|---|---|---|---|---|---|---|---|
| **Intercept** | 1.62 | 0.06 | 1.50 | 1.75 | 1.00 | 3034 | 3094 |
| **Group Glucose** | 0.10 | 0.09 | -0.08 | 0.27 | 1.00 | 3378 | 2992 |
| **Group Methylglyoxal** | -0.12 | 0.08 | -0.29 | 0.04 | 1.00 | 3208 | 3276 |
| **Month May** | -0.34 | 0.09 | -0.51 | -0.17 | 1.00 | 2954 | 3179 |
| **C_Age** | -0.14 | 0.04 | -0.21 | -0.06 | 1.00 | 3370 | 3111 |
| **Group Glucose:Month May** | 0.22 | 0.14 | -0.05 | 0.49 | 1.00 | 3607 | 3193 |
| **Group Methylglyoxal:Month May** | 0.12 | 0.12 | -0.11 | 0.34 | 1.00 | 3167 | 3013 |
| **Group Glucose:C_Age** | 0.19 | 0.06 | 0.06 | 0.30 | 1.00 | 4126 | 3217 |
| **Group Methylglyoxal:C_Age** | 0.07 | 0.06 | -0.04 | 0.19 | 1.00 | 4076 | 3445 |

**Table ESM2.18.** Outcome of loo on the sets of plasma total bile acid models with the same observations. ELP differences and their SE are presented. The higher numbers represent simpler models. The simplest model is selected if it is not different from the next most simple one (i.e. the SE of the next one is higher than its ELPD difference).

| | elpd_diff | se_diff |
|---|---|---|
| **modTBA5** | 0.0 | 0.0 |
| **modTBA7** | -0.2 | 2.6 |
| **modTBA6** | -0.3 | 1.4 |
| **modTBA4** | -0.5 | 1.3 |
| **modTBA8** | -1.0 | 2.7 |
| **modTBA3** | -2.3 | 1.3 |
| **modTBA2** | -3.5 | 1.6 |
| **modTBA1** | -4.4 | 1.5 |
| **modTBA** | -6.0 | 1.7 |

**Table ESM2.19.** Outcome from the model on plasma phosphates.

| | Estimate | Est.Error | l-95% CI | u-95% CI | Rhat | Bulk_ESS | Tail_ESS |
|---|---|---|---|---|---|---|---|
| **Intercept** | 0.75 | 0.04 | 0.66 | 0.83 | 1.00 | 2911 | 3235 |
| **Group Glucose** | -0.05 | 0.06 | -0.16 | 0.06 | 1.00 | 3070 | 2861 |
| **Group Methylglyoxal** | -0.04 | 0.05 | -0.15 | 0.06 | 1.00 | 2750 | 2257 |
| **Month May** | 0.06 | 0.06 | -0.05 | 0.18 | 1.00 | 2812 | 2868 |
| **Sex male** | -0.04 | 0.04 | -0.12 | 0.05 | 1.00 | 3495 | 3362 |
| **Group Glucose:Month May** | -0.05 | 0.09 | -0.22 | 0.12 | 1.00 | 3342 | 2864 |
| **Group Methylglyoxal:Month May** | -0.15 | 0.08 | -0.30 | -0.00 | 1.00 | 3063 | 3007 |
| **Month May:Sex male** | -0.20 | 0.07 | -0.34 | -0.08 | 1.00 | 3651 | 2838 |

**Table ESM2.20.** Outcome of loo on the sets of plasma phosphates models with the same observations (**A** with age and **B** without age, given the absence of age values for few individuals). ELP differences and their SE are presented. The higher numbers represent simpler models. The simplest model is selected if it is not different from the next most simple one (i.e. the SE of the next one is higher than its ELPD difference).

**A**

| | elpd_diff | se_diff |
|---|---|---|
| **modPHOS3** | 0.0 | 0.0 |
| **modPHOS6** | -0.5 | 2.4 |
| **modPHOS4** | -1.0 | 1.2 |
| **modPHOS2** | -1.2 | 0.7 |
| **modPHOS5** | -1.5 | 2.1 |
| **modPHOS1** | -2.1 | 1.1 |
| **modPHOS** | -5.3 | 1.0 |

**B**

| | elpd_diff | se_diff |
|---|---|---|
| **modPHOS8** | 0.0 | 0.0 |
| **modPHOS7** | -1.7 | 1.5 |

**Table ESM2.21.** Outcome from the model on plasma calcium.

| | Estimate | Est.Error | l-95% CI | u-95% CI | Rhat | Bulk_ESS | Tail_ESS |
|---|---|---|---|---|---|---|---|
| **Intercept** | 0.78 | 0.02 | 0.74 | 0.83 | 1.00 | 2302 | 3295 |
| **Group Glucose** | 0.00 | 0.03 | -0.06 | 0.06 | 1.01 | 2244 | 2501 |
| **Group Methylglyoxal** | 0.01 | 0.03 | -0.04 | 0.06 | 1.00 | 2132 | 2767 |
| **Month May** | 0.02 | 0.03 | -0.04 | 0.07 | 1.01 | 1306 | 2482 |
| **Sex male** | -0.01 | 0.03 | -0.06 | 0.05 | 1.00 | 1964 | 2682 |
| **Dilutions** | 0.07 | 0.00 | 0.06 | 0.08 | 1.00 | 6277 | 3016 |
| **Group Glucose:Month May** | 0.16 | 0.05 | 0.06 | 0.25 | 1.01 | 1765 | 2850 |
| **Group Methylglyoxal:Month May** | 0.07 | 0.04 | -0.01 | 0.15 | 1.01 | 1855 | 2574 |
| **Group Glucose:Sex male** | -0.01 | 0.04 | -0.09 | 0.07 | 1.01 | 1970 | 2348 |
| **Group Methylglyoxal:Sex male** | -0.02 | 0.04 | -0.09 | 0.06 | 1.00 | 2087 | 2828 |
| **Month May:Sex male** | 0.00 | 0.04 | -0.07 | 0.08 | 1.01 | 1504 | 2742 |
| **Group Glucose:Month May:Sex male** | -0.17 | 0.06 | -0.29 | -0.04 | 1.01 | 1835 | 2188 |
| **Group Methylglyoxal:Month May:Sex male** | -0.06 | 0.06 | -0.18 | 0.04 | 1.01 | 1559 | 3019 |

**Table ESM2.22.** Outcome of loo on the sets of plasma calcium models with the same observations. ELP differences and their SE are presented. The higher numbers represent simpler models. The simplest model is selected if it is not different from the next most simple one (i.e. the SE of the next one is higher than its ELPD difference).

| | elpd_diff | se_diff |
|---|---|---|
| **modCa5** | 0.0 | 0.0 |
| **modCa4** | -1.8 | 1.3 |
| **modCa3** | -2.3 | 1.6 |
| **modCa2** | -2.9 | 1.9 |
| **modCa1** | -3.6 | 2.1 |
| **modCa** | -4.4 | 2.0 |

**Table ESM2.23.** Outcome from the model on plasma potassium.

| | Estimate | Est.Error | l-95% CI | u-95% CI | Rhat | Bulk_ESS | Tail_ESS |
|---|---|---|---|---|---|---|---|
| **Intercept** | 0.71 | 0.04 | 0.63 | 0.78 | 1.00 | 1694 | 2545 |
| **Group Glucose** | -0.05 | 0.06 | -0.17 | 0.07 | 1.00 | 1918 | 2448 |
| **Group Methylglyoxal** | -0.06 | 0.05 | -0.16 | 0.04 | 1.00 | 1814 | 2529 |
| **Month May** | 0.20 | 0.05 | 0.10 | 0.30 | 1.00 | 1842 | 2237 |
| **Sex male** | 0.02 | 0.05 | -0.08 | 0.13 | 1.00 | 1572 | 2214 |
| **Group Glucose:Month May** | -0.06 | 0.09 | -0.24 | 0.12 | 1.00 | 2202 | 2475 |
| **Group Methylglyoxal:Month May** | -0.19 | 0.08 | -0.35 | -0.03 | 1.00 | 1977 | 2717 |
| **Group Glucose:Sex male** | -0.04 | 0.08 | -0.20 | 0.12 | 1.00 | 1803 | 2417 |
| **Group Methylglyoxal:Sex male** | -0.00 | 0.07 | -0.14 | 0.14 | 1.00 | 1685 | 2252 |
| **Month May:Sex male** | -0.27 | 0.08 | -0.44 | -0.11 | 1.00 | 1708 | 2333 |
| **Group Glucose:Month May:Sex male** | 0.20 | 0.13 | -0.06 | 0.46 | 1.00 | 2018 | 2621 |
| **Group Methylglyoxal:Month May:Sex male** | 0.26 | 0.12 | 0.03 | 0.48 | 1.00 | 1851 | 2742 |

**Table ESM2.24.** Outcome of loo on the sets of plasma potassium models with the same observations. ELP differences and their SE are presented. The higher numbers represent simpler models. The simplest model is selected if it is not different from the next most simple one (i.e. the SE of the next one is higher than its ELPD difference).

| | elpd_diff | se_diff |
|---|---|---|
| **modK5** | 0.0 | 0.0 |
| **modK4** | -1.5 | 1.1 |
| **modK3** | -2.5 | 1.3 |
| **modK2** | -3.1 | 1.3 |
| **modK1** | -4.3 | 1.3 |
| **modK** | -6.3 | 1.4 |

**Table ESM2.25.** Outcome from the model on plasma sodium.

| | Estimate | Est.Error | l-95% CI | u-95% CI | Rhat | Bulk_ESS | Tail_ESS |
|---|---|---|---|---|---|---|---|
| **Intercept** | 2.15 | 0.01 | 2.13 | 2.17 | 1.00 | 3919 | 3763 |
| **Group Glucose** | -0.01 | 0.01 | -0.04 | 0.01 | 1.00 | 2746 | 2911 |
| **Group Methylglyoxal** | -0.02 | 0.01 | -0.04 | 0.01 | 1.00 | 2691 | 2984 |
| **Month May** | 0.07 | 0.01 | 0.04 | 0.09 | 1.00 | 2370 | 2835 |
| **Dilutions** | 0.09 | 0.00 | 0.08 | 0.10 | 1.00 | 5272 | 2931 |
| **Group Glucose:Month May** | 0.00 | 0.02 | -0.04 | 0.05 | 1.00 | 2730 | 3163 |
| **Group Methylglyoxal:Month May** | 0.01 | 0.02 | -0.02 | 0.05 | 1.00 | 2566 | 2638 |

**Table ESM2.26.** Outcome of loo on the sets of plasma sodium models with the same observations (**A** with age and **B** without age, given the absence of age values for few individuals). ELP differences and their SE are presented. The higher numbers represent simpler models. The simplest model is selected if it is not different from the next most simple one (i.e. the SE of the next one is higher than its ELPD difference).

A

| | elpd_diff | se_diff |
|---|---|---|
| **modNa4** | 0.0 | 0.0 |
| **modNa5** | -0.2 | 1.3 |
| **modNa3** | -1.0 | 0.6 |
| **modNa2** | -2.1 | 0.9 |
| **modNa1** | -3.7 | 0.9 |
| **modNa** | -5.9 | 1.2 |

B

| | elpd_diff | se_diff |
|---|---|---|
| **modNa10** | 0.0 | 0.0 |
| **modNa9** | -1.0 | 0.1 |
| **modNa8** | -1.8 | 0.6 |
| **modNa7** | -3.2 | 1.5 |
| **modNa6** | -3.6 | 2.7 |

**Table ESM2.27.** Outcome from the model on H/L ratio.

| | Estimate | Standard Error | df | t value | P-value |
|---|---|---|---|---|---|
| **Intercept** | 0.649501 | 0.040123 | 200.000000 | 16.188 | $< 2*10^{-16}$ |
| **Group Glucose** | 0.060613 | 0.056360 | 200.000000 | 1.075 | 0.28346 |
| **Group Methylglyoxal** | -0.151353 | 0.048771 | 200.000000 | -3.103 | 0.00219 |
| **Month February** | -0.299909 | 0.057370 | 200.000000 | -5.228 | $4.31*10^{-7}$ |
| **Month May** | -0.126138 | 0.060150 | 200.000000 | -2.097 | 0.03724 |
| **Month August** | -0.267599 | 0.060502 | 200.000000 | -4.423 | $1.60*10^{-5}$ |
| **Sex male** | 0.008004 | 0.042664 | 200.000000 | 0.188 | 0.85138 |
| **Group Glucose:Month February** | -0.001317 | 0.083460 | 200.000000 | -0.016 | 0.98742 |
| **Group Methylglyoxal:Month February** | 0.154378 | 0.070997 | 200.000000 | 2.174 | 0.03085 |
| **Group Glucose:Month May** | 0.109716 | 0.088462 | 200.000000 | 1.240 | 0.21633 |
| **Group Methylglyoxal:Month May** | 0.139287 | 0.073926 | 200.000000 | 1.884 | 0.06100 |
| **Group Glucose:Month August** | 0.154656 | 0.091290 | 200.000000 | 1.694 | 0.09180 |
| **Group Methylglyoxal:Month August** | 0.194767 | 0.074976 | 200.000000 | 2.598 | 0.01008 |
| **Month February:Sex male** | 0.090738 | 0.062746 | 200.000000 | 1.446 | 0.14971 |
| **Month May:Sex male** | -0.118180 | 0.065968 | 200.000000 | -1.791 | 0.07473 |
| **Month August:Sex male** | -0.080889 | 0.067580 | 200.000000 | -1.197 | 0.23274 |

**Table ESM2.28.** Outcome from the model on monocytes counts.

| | Estimate | Standard Error | df | t value | P-value |
|---|---|---|---|---|---|
| **Intercept** | 2.417 | $5.819*10^{-2}$ | $1.995*10^{2}$ | 41.528 | $< 2*10^{-16}$ |
| **Group Glucose** | $1.991*10^{-2}$ | $9.330*10^{-2}$ | $1.994*10^{2}$ | 0.213 | 0.83124 |
| **Group Methylglyoxal** | 0.1251 | $8.709*10^{-2}$ | $1.999*10^{2}$ | 1.436 | 0.15255 |
| **Month February** | -0.5739 | $8.203*10^{-2}$ | $1.530*10^{2}$ | -6.996 | $7.72*10^{-11}$ |
| **Month May** | -0.6084 | $8.553*10^{-2}$ | $1.588*10^{2}$ | -7.114 | $3.65*10^{-11}$ |
| **Month August** | -0.2736 | $8.567*10^{-2}$ | $1.592*10^{2}$ | -3.194 | 0.00169 |
| **C_WBC** | $1.563*10^{-4}$ | $1.257*10^{-5}$ | $1.961*10^{2}$ | 12.432 | $< 2*10^{-16}$ |
| **Group Glucose:Month February** | -0.1736 | 0.1356 | $1.554*10^{2}$ | -1.280 | 0.20245 |
| **Group Methylglyoxal:Month February** | $-6.715*10^{-3}$ | 0.1198 | $1.591*10^{2}$ | -0.056 | 0.95535 |
| **Group Glucose:Month May** | $-3.579*10^{-2}$ | 0.1432 | $1.623*10^{2}$ | -0.250 | 0.80299 |
| **Group Methylglyoxal:Month May** | $4.641*10^{-2}$ | 0.1235 | $1.641*10^{2}$ | 0.376 | 0.70763 |
| **Group Glucose:Month August** | $-3.043*10^{-2}$ | 0.1467 | $1.647*10^{2}$ | -0.207 | 0.83591 |
| **Group Methylglyoxal:Month August** | -0.3851 | 0.1296 | $1.687*10^{2}$ | -2.970 | 0.00341 |

**Table ESM2.29.** Outcome from the model on heterophils counts.

| | Estimate | Standard Error | df | t value | P-value |
|---|---|---|---|---|---|
| **Intercept** | 2.396 | $5.829*10^{-2}$ | 200 | 41.111 | $< 2*10^{-16}$ |
| **GroupGlucose** | 0.1361 | $9.344*10^{-2}$ | 200 | 1.456 | 0.14686 |
| **GroupMethylglyoxal** | $-1.01510^{-2}$ | $8.728*10^{-2}$ | 200 | -0.116 | 0.90752 |
| **MonthFebruary** | -0.8709 | $8.360*10^{-2}$ | 200 | -10.418 | $< 2*10^{-16}$ |
| **MonthMay** | -0.8069 | $8.704*10^{-2}$ | 200 | -9.270 | $< 2*10^{-16}$ |
| **MonthAugust** | -0.8918 | $8.717*10^{-2}$ | 200 | -10.230 | $< 2*10^{-16}$ |
| **C_WBC** | $1.028*10^{-4}$ | $1.264*10^{-5}$ | 200 | 8.127 | $4.5*10^{-14}$ |
| **GroupGlucose:MonthFebruary** | -0.2327 | 0.1381 | 200 | -1.685 | 0.09360 |
| **GroupMethylglyoxal:MonthFebruary** | 0.2297 | 0.1219 | 200 | 1.885 | 0.06092 |
| **GroupGlucose:MonthMay** | $-9.108*10^{-2}$ | 0.1457 | 200 | -0.625 | 0.53249 |
| **GroupMethylglyoxal:MonthMay** | 0.3436 | 0.1255 | 200 | 2.737 | 0.00676 |
| **GroupGlucose:MonthAugust** | 0.1886 | 0.1490 | 200 | 1.265 | 0.20719 |
| **GroupMethylglyoxal:MonthAugust** | -0.4091 | 0.1316 | 200 | -3.108 | 0.00216 |

**Table ESM2.30.** Outcome from the model on lymphocytes counts.

| | Estimate | Standard Error | df | t value | P-value |
|---|---|---|---|---|---|
| **Intercept** | 2.806 | $5.511*10^{-2}$ | 200 | 50.920 | $< 2*10^{-16}$ |
| **GroupGlucose** | $2.350*10^{-2}$ | $8.835*10^{-2}$ | 200 | 0.266 | 0.790553 |
| **GroupMethylglyoxal** | $3.665*10^{-2}$ | $8.252*10^{-2}$ | 200 | 0.444 | 0.657441 |
| **MonthFebruary** | -0.3254 | $7.904*10^{-2}$ | 200 | -4.117 | $5.62*10^{-5}$ |
| **MonthMay** | -0.4134 | $8.229*10^{-2}$ | 200 | -5.024 | $1.12*10^{-6}$ |
| **MonthAugust** | -0.3888 | $8.242*10^{-2}$ | 200 | -4.718 | 4.48e-06 |
| **C_WBC** | $1.735*10^{-4}$ | $1.195*10^{-5}$ | 200 | 14.516 | $< 2*10^{-16}$ |
| **GroupGlucose:MonthFebruary** | -0.2187 | 0.1306 | 200 | -1.675 | 0.095514 |
| **GroupMethylglyoxal:MonthFebruary** | $8.851*10^{-2}$ | 0.1152 | 200 | 0.768 | 0.443227 |
| **GroupGlucose:MonthMay** | -0.1861 | 0.1377 | 200 | -1.352 | 0.178015 |
| **GroupMethylglyoxal:MonthMay** | 0.2382 | 0.1187 | 200 | 2.007 | 0.046148 |
| **GroupGlucose:MonthAugust** | $-5.027*10^{-2}$ | 0.1409 | 200 | -0.357 | 0.721654 |
| **GroupMethylglyoxal:MonthAugust** | -0.4179 | 0.1244 | 200 | -3.358 | 0.000939 |

The dilution factor had a positive effect on plasma calcium levels (**Figure ESM2.2.A**; α ± SE: 0.78 ± 0.02, $CI_{95}$[0.74, 0.83]; β ± SE: 0.07 ± 0.00, $CI_{95}$[0.06, 0.08]), but especially on sodium (**Figure ESM2.2.B**; α ± SE: 2.15 ± 0.01, $CI_{95}$[2.12, 2.17]; β ± SE: 0.09 ± 0.00, $CI_{95}$[0.08, 0.1]), rendering the values of the latter unreliable, so that the higher May levels observed for all groups in this electrolyte (**Figure ESM2.3**; November - May: -0.072, $CI_{95}$[-0.089, -0.055]) are potentially an artefact.

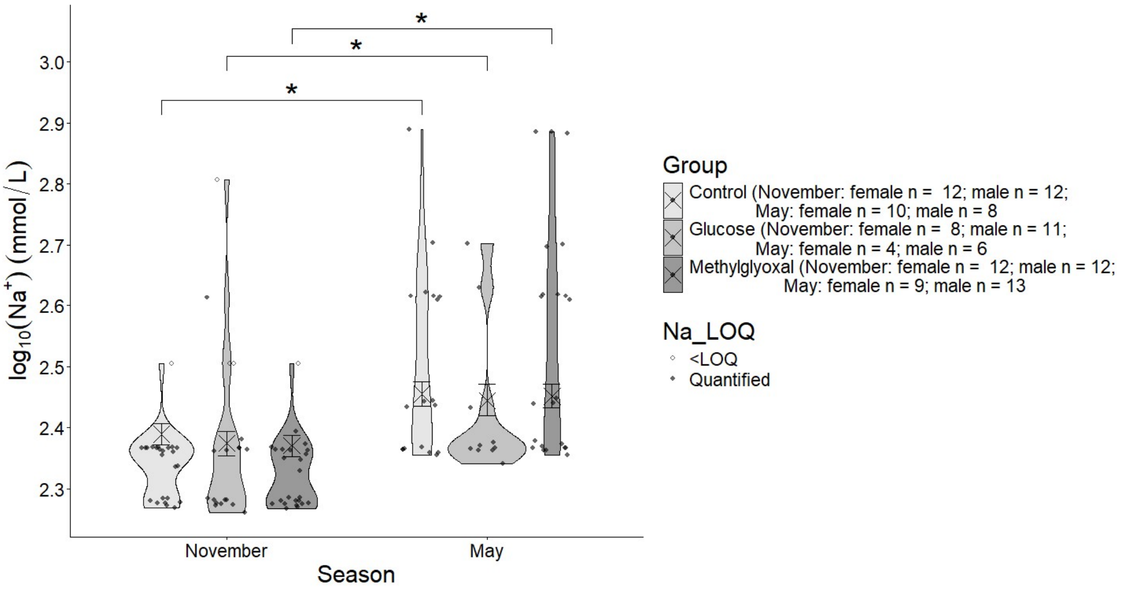


**Figure ESM2.2~~3~~.** Variation in ($log_{10}$) plasma sodium concentration across treatments and months. Crosses and error bars represent model-estimated marginal means ±95% CI. Annotations indicating the support of the effects are based on pairwise contrasts performed separately within each month and treatment.

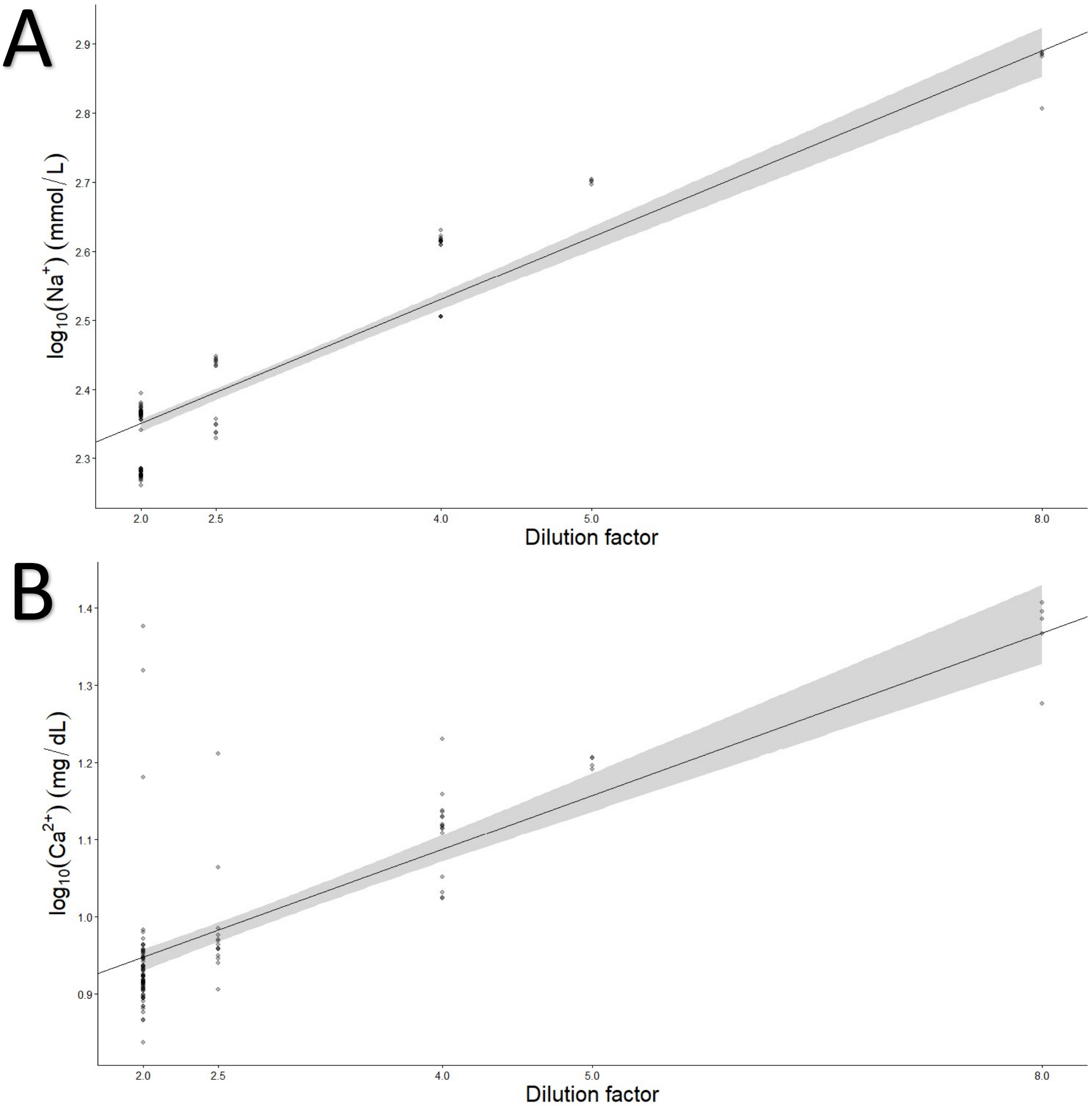


**Figure ESM2.~~2~~3.** Effects of sample dilution on measured ($log_{10}$) plasma (**A**) sodium and (**B**) calcium levels. Crosses and error bars represent model-estimated marginal means ±95% CI. Grey areas in plots **A** and **B** represent estimated ±95% CI.

### Seasonal effects on leukocytes counts

#### Heterophiles

Seasonally, all months showed lower heterophil counts than November across all treatment groups (**Figure ESM2.3.A**; Control: November - February: marginal difference ± SE: 0.871 ± 0.086, $t = 10.07$, $P < 0.0001$; November - May: 0.807 ± 0.09, $t = 8.954$, $P < 0.0001$; November - August: 0.892 ± 0.09, $t = 9.88$, $P < 0.0001$; Glucose: November - February: marginal difference ± SE: 1.104 ± 0.114, $t = 9.66$, $P < 0.0001$; November - May: 0.898 ± 0.122, $t = 7.387$, $P < 0.0001$; November - August: 0.703 ± 0.126, $t = 5.594$, $P < 0.0001$; Methylglyoxal: November - February: 0.641 ± 0.095, $t = 6.761$, $P < 0.0001$; November - May: 0.463 ± 0.096, $t = 4.816$, $P < 0.0001$;

November - August: 1.301 ± 0.099, t = 13.097, P < 0.0001). For the methylglyoxal group, August levels were lower than those in February (February - August: 0.659 ± 0.094, t = 7.016, P < 0.0001) and May (May - August: 0.838 ± 0.097, t = 8.619, P < 0.0001). For the glucose group, August levels were higher than those in February (February - August: -0.4 ± 0.132, t = -3.024, P = 0.015).

#### Lymphocytes

Seasonally, lymphocyte levels decreased after November for both control (**Figure ESM2.3.B**; November - February: 0.325 ± 0.082, t = 3.979, P = 0.001; November - May: 0.413 ± 0.085, t = 4.852, P < 0.0001; November - August: 0.389 ± 0.085, t = 4.556, P = 0.0001) and glucose group (November - February: 0.544 ± 0.108, t = 5.037, P < 0.0001; November - May: 0.599 ± 0.115, t = 5.216, P < 0.0001; November - August: 0.439 ± 0.119, t = 3.695, P = 0.002). For methylglyoxal-supplemented birds, lymphocyte counts were lower in February (November - February: 0.237 ± 0.09, t = 2.641, P = 0.044) and August (November - August: 0.807 ± 0.094, t = 8.59, P < 0.0001). August showed lower lymphocyte count than February (February - August: 0.57 ± 0.089, t = 6.412, P < 0.0001) and May (May - August: 0.631 ± 0.092, t = 6.873, P < 0.0001) within the methylglyoxal group.

#### Monocytes

Seasonally, control birds showed a decrease in monocyte counts after November (Main text - **Figure 10.C**; November - February: 0.574 ± 0.085, t = 6.759, P < 0.0001; November - May: 0.608 ± 0.089, t = 6.869, P < 0.0001; November - August: 0.274 ± 0.089, t = 3.083, P = 0.013), followed by an increase in August compared to February and May (February - August: -0.3 ± 0.09, t = -3.321, P = 0.006; May - August: -0.335 ± 0.093, t = -3.586, P = 0.002). Methylglyoxal-supplemented birds also showed a decrease after November (November - February: 0.581 ± 0.093, t = 6.219, P < 0.0001; November - May: 0.562 ± 0.095, t = 5.931, P < 0.0001; November - August: 0.659 ± 0.098, t = 6.728, P < 0.0001), without a later increase. For the glucose group, monocyte counts decreased in February and May compared to November (November - February: 0.747 ± 0.112, t = 6.658, P < 0.0001; November - May: 0.644 ± 0.12, t = 5.384, P < 0.0001), with August levels higher than those in February (February - August: -0.443 ± 0.13, t = -3.406, P = 0.004).

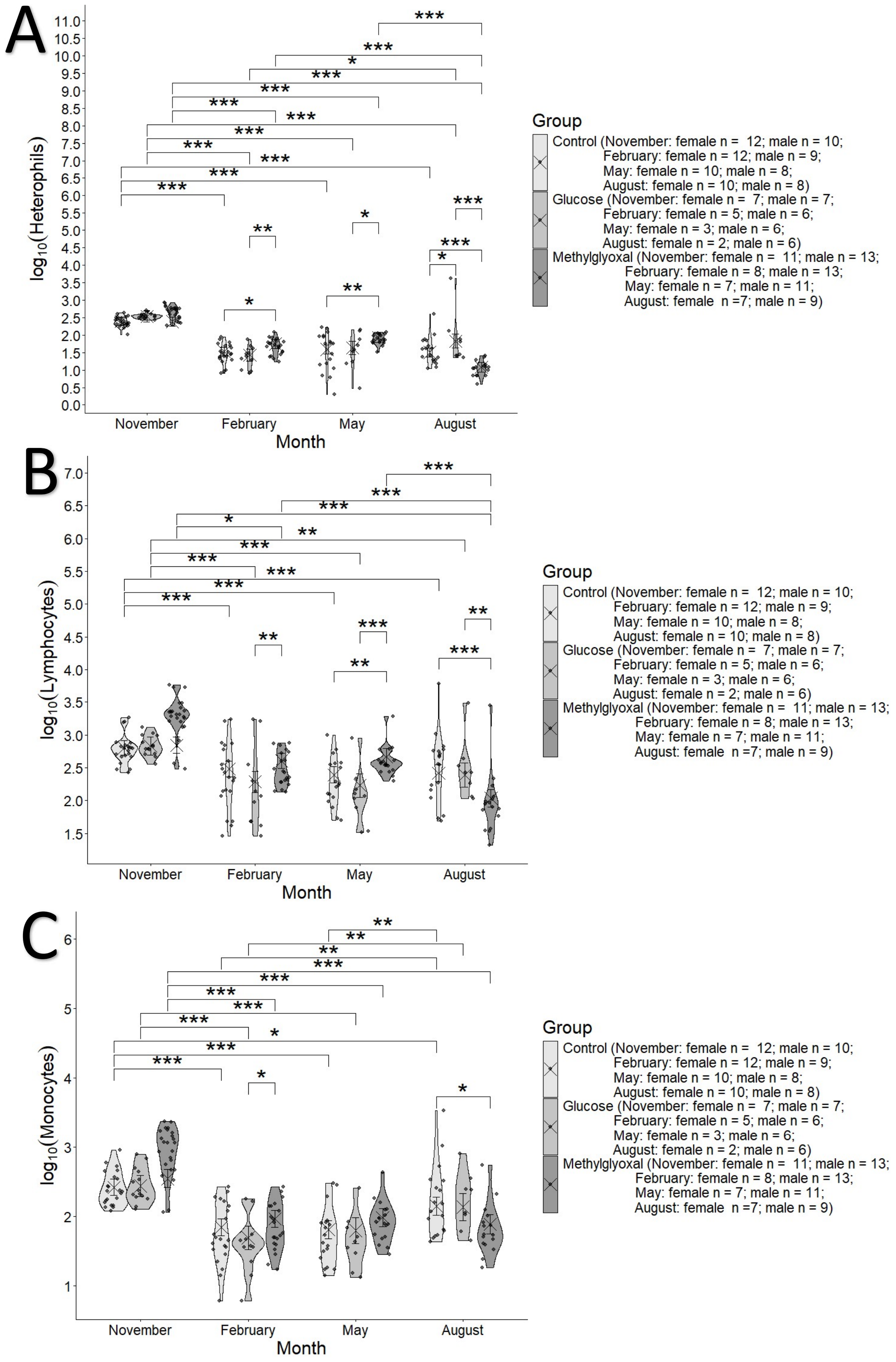


**Figure ESM2.4.** Variation in ($log_{10}$) (**A**) heterophils, (**B**) lymphocyte and (**C**) monocyte number, adjusting by total (centred) leukocyte number in the models, across treatments and months. Crosses and error bars represent model-estimated marginal means ±95% CI. Significance annotations are based on pairwise contrasts performed separately within each month and treatment. The number of asterisks represent the level of significance (* P < 0.05; ** P < 0.01; *** P < 0.001).

#### Box 1. Structure of the final models included in the manuscript

##### Bayesian general linear mixed models fit with brms()

log10(ALB)| cens(ALB_QL) ~ Group*Month + (1|Bird_ID)

log10(GLOB)|cens(GLOB_DL_mod) ~ Group*Month*Sex-(Group:Month:Sex) + (1|Bird_ID)

log10(TP)|cens(TP_QL) ~ Group*Month*Sex-(Group:Month:Sex)-(Month:Sex) + C_BM + (1|Bird_ID)

log10(A_G)|cens(A_G_DL_mod) ~ Group*Month + (1|Bird_ID)

log10(AST)|cens(AST_UDL) ~ Group*Month + C_Age + (1|Bird_ID)

log10(TBA)|cens(TBA_QL) ~ Group*Month+ Group*C_Age + (1|Bird_ID)

log10(CK) ~ Group*Month + C_Age + (1|Bird_ID)

log10(UA)| cens(UA_QL) ~ Group*Month*Sex + C_Age + C_BM + (1|Bird_ID)

GLU ~ Group*Month + C_BM + (1|Bird_ID)

log10(Ca) ~ Group*Month*Sex + Dilutions + (1|Bird_ID)

log10(PHOS)|cens(PHOS_QL) ~ Group*Month*Sex -(Group:Month:Sex) -Group:Sex + (1|Bird_ID)

log10(K)|cens(K_QL) ~ Group*Month*Sex + (1|Bird_ID)

log10(Na)|cens(Na_QL) ~ Group*Month + Dilutions + (1|Bird_ID)

##### Frequentist general linear mixed models fit with lmer()

sqrt(H_L) ~ Group*Month+Month*Sex + (1| Bird_ID)

log10(Monocytes) ~ Group*Month + C_WBC + (1| Bird_ID)

log10(Heterophils) ~ Group*Month + C_WBC + (1| Bird_ID)

log10(Lymphocytes) ~ Group*Month+C_WBC + (1| Bird_ID)
