## Supplementary material for "Glucose and methylglyoxal alter plasma protein dynamics, immune traits and glucose homeostasis in a sex- and season-dependent manner in zebra finches": ESM3 - Weather

### ESM 2. Weather variations

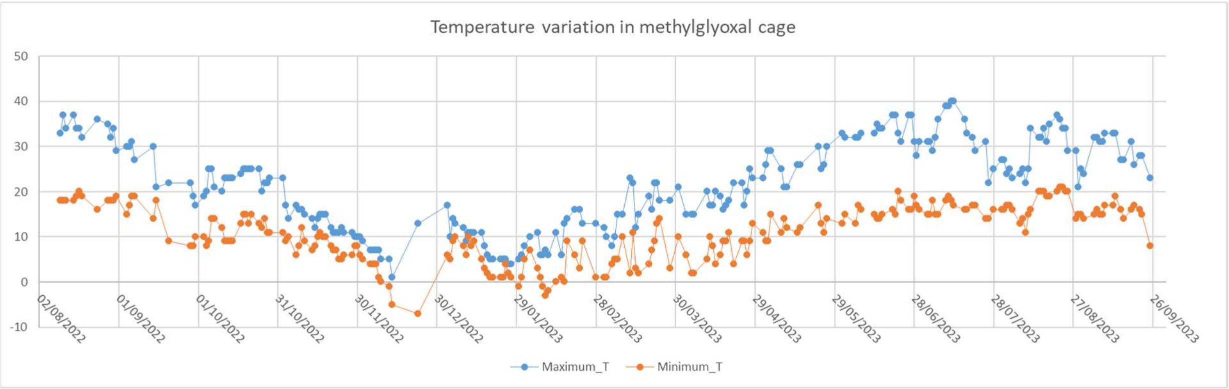


**Figure ESM3.1.** Minimum and maximum temperature variation along the year of the experiment in the cage from methylglyoxal supplementation group.

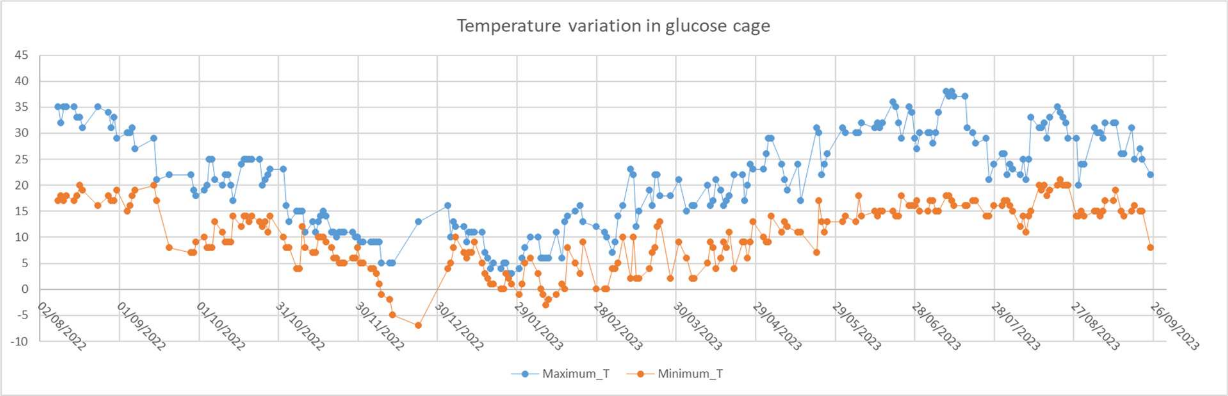


**Figure ESM3.2.** Minimum and maximum temperature variation along the year of the experiment in the cage from glucose supplementation group.

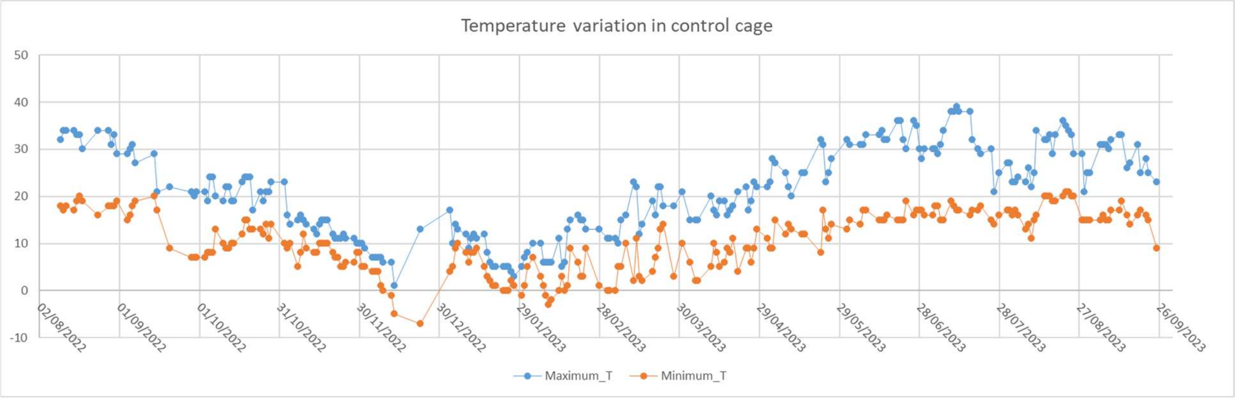


**Figure ESM3.3.** Minimum and maximum temperature variation along the year of the experiment in the cage from control group.

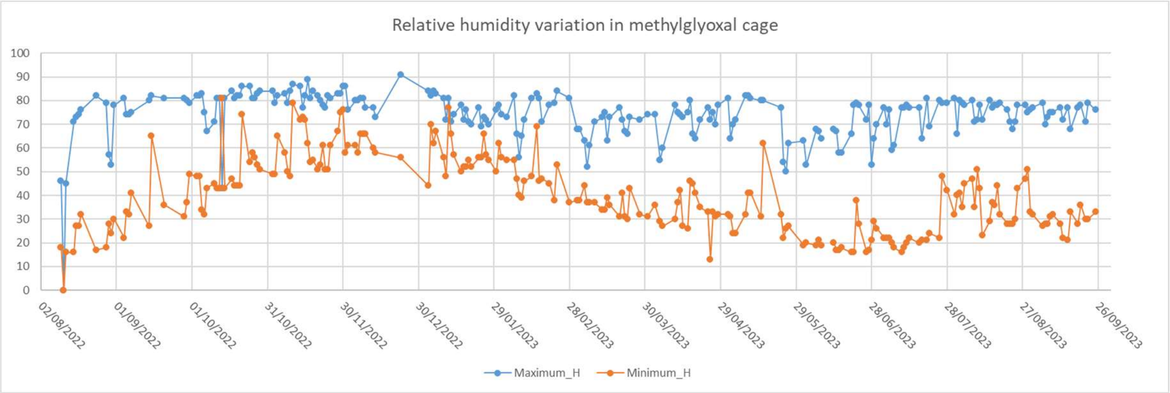


**Figure ESM3.4.** Minimum and maximum relative humidity variation along the year of the experiment in the cage from methylglyoxal supplementation group.

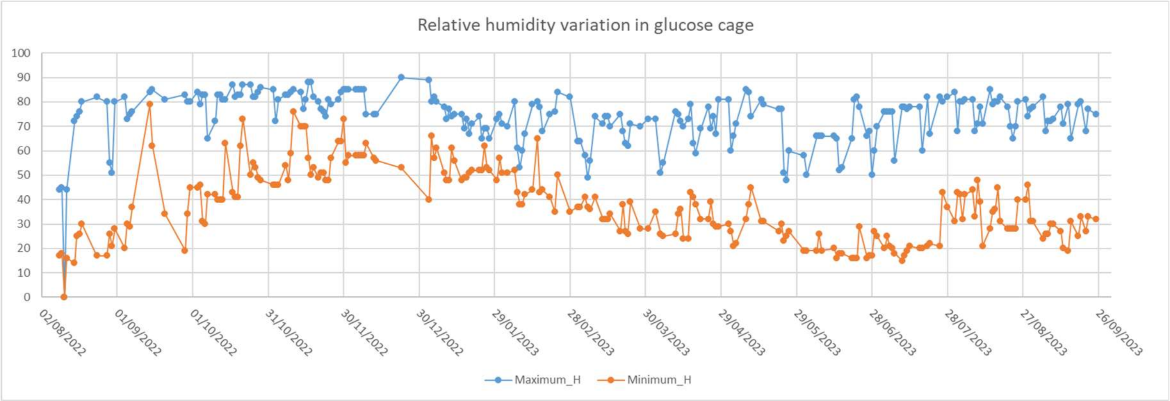


**Figure ESM3.5.** Minimum and maximum relative humidity variation along the year of the experiment in the cage from glucose supplementation group.

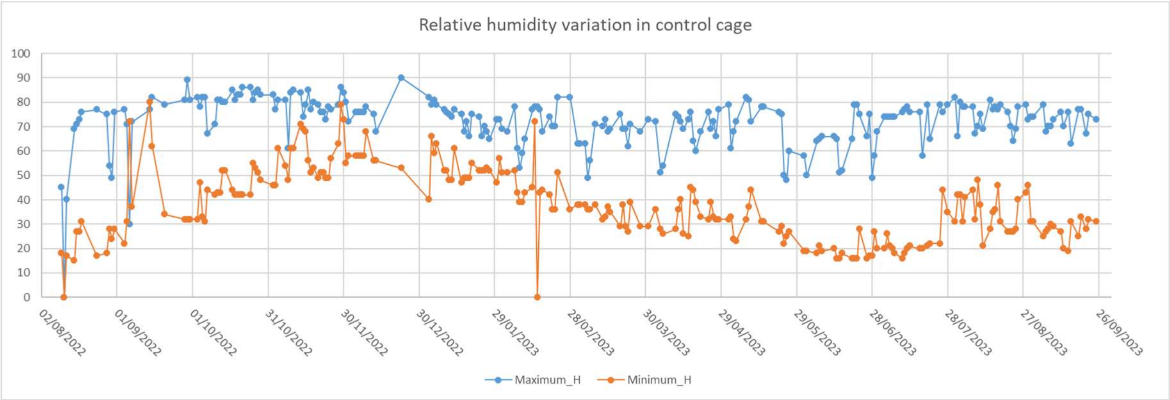


**Figure ESM3.6.** Minimum and maximum relative humidity variation along the year of the experiment in the cage from control supplementation group.
